# Disrupting cellular memory to overcome drug resistance

**DOI:** 10.1101/2022.06.16.496161

**Authors:** Guillaume Harmange, Raúl A. Reyes Hueros, Dylan Schaff, Benjamin Emert, Michael Saint-Antoine, Shivani Nellore, Mitchell E. Fane, Gretchen M. Alicea, Ashani T. Weeraratna, Abhyudai Singh, Sydney M. Shaffer

## Abstract

Plasticity enables cells to change their gene expression state in the absence of a genetic change. At the single-cell level, these gene expression states can persist for different lengths of time which is a quantitative measurement referred to as gene expression memory. Because plasticity is not encoded by genetic changes, these cell states can be reversible, and therefore, are amenable to modulation by disrupting gene expression memory. However, we currently do not have robust methods to find the regulators of memory or to track state switching in plastic cell populations. Here, we developed a lineage tracing-based technique to quantify gene expression memory and to identify single cells as they undergo cell state transitions. Applied to human melanoma cells, we quantified long-lived fluctuations in gene expression that underlie resistance to targeted therapy. Further, we identified the PI3K and TGF-β pathways as modulators of these state dynamics. Applying the gene expression signatures derived from this technique, we find that these expression states are generalizable to in vivo models and present in scRNA-seq from patient tumors. Leveraging the PI3K and TGF-β pathways as dials on memory between plastic states, we propose a “ pretreatment” model in which we first use a PI3K inhibitor to modulate the expression states of the cell population and then apply targeted therapy. This plasticity informed dosing scheme ultimately yields fewer resistant colonies than targeted therapy alone. Taken together, we describe a technique to find modulators of gene expression memory and then apply this knowledge to alter plastic cell states and their connected cell fates.

## Introduction

Gene expression memory describes the length of time that a particular expression state exists in an individual cell or lineage of cells. Gene expression memory can exist over a range of different timescales and is ultimately a quantitative measurement (Angel et al. 2011; Shaffer et al. 2020). For cancer, an intermediate timescale of memory has been associated with a number of important phenotypes including, stemness and differentiation (Chaligne et al. 2021; Gupta et al. 2011), metastasis (Kaur et al. 2022), and therapy resistance (Shaffer et al. 2020; Torre et al. 2021; Sharma et al. 2010; Oren et al. 2021). In these examples, the cellular state underlying the phenotype is stable enough to persist through multiple cell divisions, but is ultimately not permanent, and thus amenable to switching states. For cancer therapy resistance, drug-naive cells seemingly randomly fluctuate between two states, one which is susceptible to targeted therapy and another which would become resistant if the drug is applied, termed “ primed” for drug resistance (Emert et al. 2021; Shaffer et al. 2017). However, the molecular drivers underlying these heritable, but reversible, memory states remain unknown.

Because drug resistance can emerge from cells that are in a primed gene expression state, one intriguing possibility to prevent resistance is to transform primed cells into drug-susceptible cells. However, we currently do not know the molecular cues that trigger cells to switch between these states. Knowing these cues would make it possible to therapeutically target state switching pathways to drive cells out of the primed gene expression state and sensitize them to therapy. Such an approach requires deep characterization of the processes underlying state switching, which remains exceedingly difficult with current methods.

Currently, there are limited techniques that will reveal memory and state switching in single cells. Our previous work inferred cellular memory from bulk RNA-seq measurements (Shaffer et al. 2020), but failed to capture drivers of switching between memory states. Meanwhile, scRNA-seq can capture the heterogeneity of a population, but it fails to capture the timescales for which different states have been present in individual cells, and lacks the resolution to capture transitioning gene expression states. A number of computational and experimental techniques have been developed to resolve time in single-cells on short timescales, on the order of hours (La Manno et al. 2018; Qiu et al. 2020), but few exist for the longer timescales of days to weeks as needed to track cellular memory. Recent advances in high-throughput cellular barcoding technologies have made it possible to track cellular lineages across any length of time (Bhang et al. 2015). Pairing cellular barcoding with scRNA-seq enables us to now match cellular lineages with their transcriptome (Biddy et al. 2018; Oren et al. 2021; Weinreb et al. 2020; Emert et al. 2021), and thus is an ideal tool for tracking transcriptional states across lineages to measure cellular memory.

Here, we developed an ultra-sensitive technique for measuring memory that is built from cell barcoding and scRNA-seq called scMemorySeq. Our experimental design uses a precisely controlled number of cell divisions to capture lineages that have undergone switching between drug-susceptible and primed states in cancer. We applied scMemorySeq to drug naive melanoma cells and found that the drug-susceptible and primed states are two very distinct, high-memory states that appear unrelated based only on scRNA-seq alone. However, through lineage tracing with cell barcoding, we identified lineages containing cells from both states, directly showing that single cells can switch between states. By analyzing lineages that lose memory of their state, we identified TGF-β and PI3K as pathways controlling state switching. Ultimately, we found that by initially disrupting the primed state through PI3K inhibition and then applying BRAF inhibitor in combination with a MEK inhibitor (BRAFi/MEKi), we could reduce the frequency of drug resistance. Taken together, we demonstrate the feasibility of molecularly targeting memory and state switching to eliminate gene expression states in cancer that prime cells for undesirable phenotypes.

## Results

We first sought to identify the molecular pathways underlying cellular memory in the drug-susceptible and primed cells in melanoma (Fig. 1A). We developed a technique called scMemorySeq that specifically captures heritable gene expression states and switching between plastic states using cellular barcoding and scRNA-seq. In our experimental design, the cellular barcodes enable high-throughput tracking of cells over any desired period of time and the scRNA-seq reveals the transcriptional states of every cell. Ultimately, we infer cellular memory by examining gene expression across lineages of related cells. For genes or sets of genes that have memory over the experimental timescale, all cells from the same lineage should be in the same end state and match the initial state (Fig. 1B). However, if cellular states change over the experimental timescale, then the end-state will contain cells with multiple gene expression states, some of which are different from the initial state. We applied this technique to BRAF-mutant melanoma cells in which a rare subpopulation of cells are resistant to BRAF inhibitors. We transduced WM989 melanoma cells with a high-complexity viral barcode library consisting of a transcribed 100 base pair semi-random barcode sequence in the 3’ UTR of GFP (Emert et al. 2021) (Fig. 1C). In this experimental model, our phenotype of interest (cells that are primed for resistance) is rare. To improve our ability to capture primed cells, and to establish the initial state of lineages, we sorted cells based on known primed cell markers (EGFR+ and NGFR+) and also isolated a matched control population (Fig. 1C). To measure cellular memory, we allowed the cells to expand through roughly 4 doublings (12-14 days). Over this time, we expect that most cells will maintain memory of their initial gene expression state, but that a subset of cells will lose memory of their initial state, thereby capturing the reversion of these expression states. At the endpoint, we harvested these cells for scRNA-seq and performed a PCR side reaction to specifically amplify the lineage barcodes from each cell.

**Figure 1:**
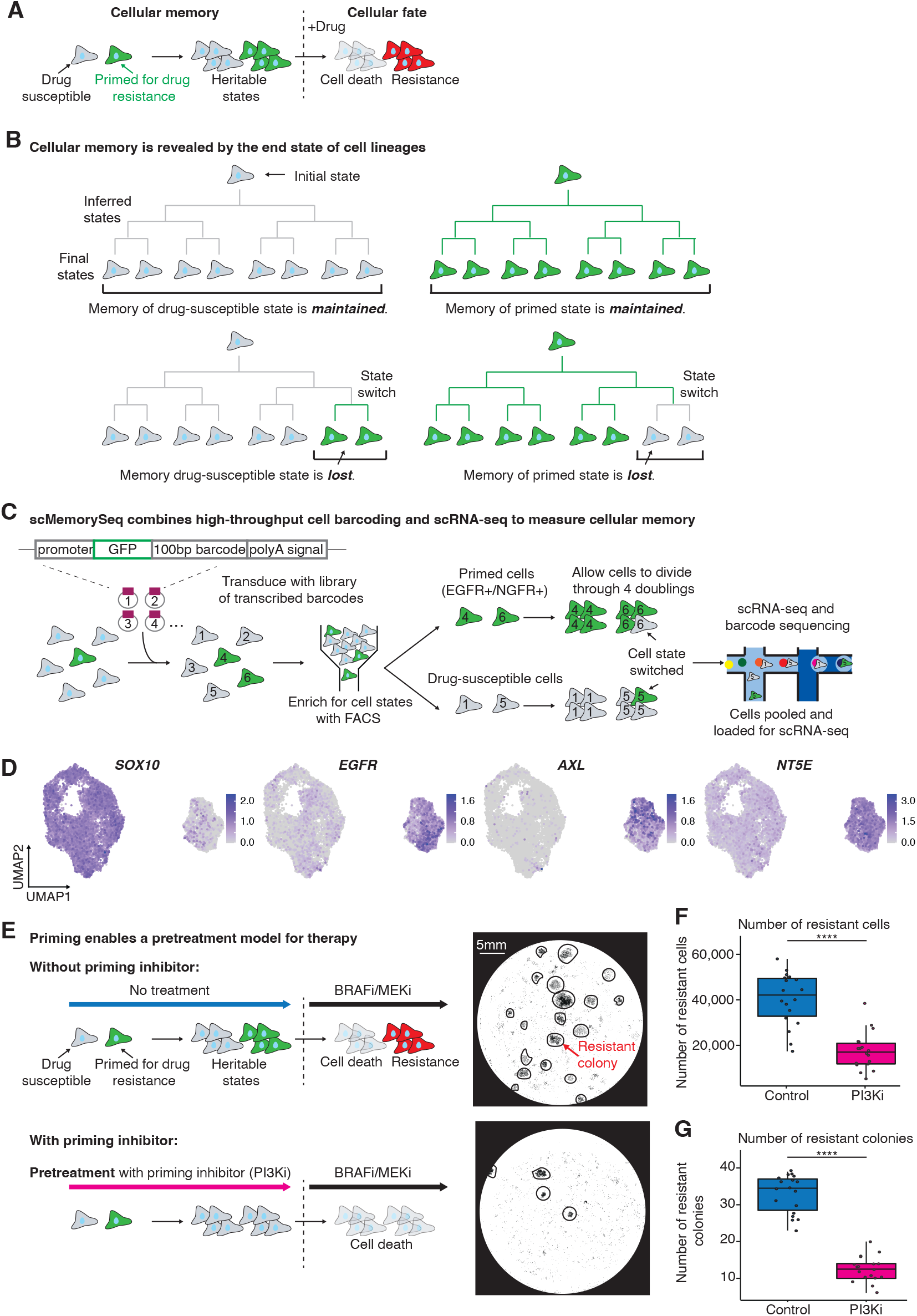
Cells primed for drug resistance have memory and can be targeted to modulate the resistance phenotype. **A**. Model for how the heritable primed state leads to resistance to targeted therapy. Cells primed for drug resistance (in green) proliferate and pass on their gene expression state through cell division, which demonstrates the concept of gene expression memory. These primed cells survive treatment with BRAFi and MEKi (resistant cells in red) while the drug-susceptible cells (in gray) die. **B**. Schematic of lineages that maintain memory compared to those that lose memory. In lineages that maintain memory, the end state of all the cells is the same as the initial state. In lineages where memory is lost, the endstate comprises a mixture of cell states. **C**. Schematic of the experimental design that we used to capture the transcriptome and lineage of cells. We transduced melanoma cells (WM989) with a high-complexity library of lentiviral lineage barcodes. We then sorted a sample of primed cells (based on EGFR and NGFR) and a mixed population. This sorting step provides us with the initial state of the cells. We then allowed the cells to undergo ∼4 doublings to build out the lineages before doing scRNA-seq and barcode sequencing on the cells. **D**. UMAP plots of the per cell log_10_ normalized gene expression for the drug-susceptible cell marker *SOX10*, and the primed cell markers *EGFR, AXL*, and *NT5E*. **E**. Schematic of pretreatment model. Without a primed state inhibitor, primed cell states are heritable through cell division and primed cells become resistant upon application of drug. With a primed state inhibitor, the primed state is eliminated from the cell population during the “ pretreatment” phase. When the drug is applied, fewer cells become resistant because the primed population is eliminated. Adjacent to the schematic are example images of fixed cells stained with DAPI after 5 days of pretreatment and 4 weeks of selection with BRAFi and MEKi. Drug-resistant colonies are circled in black where readily identifiable. Scale bar in the top image represents 5mm and applies to both images. **F**. Box plot quantifying the number of drug-resistant cells from image scans like those shown in E across 3 biological replicates, each with 6 technical replicates. P values were calculated using a wilcoxon test (****: p <= 0.0001). **G**. Box plot quantifying the number of drug colonies (for each condition where distinct colonies were present) from image scans like those shown in E across 3 biological replicates, each with 6 technical replicates. P values were calculated using a wilcoxon test (****: p <= 0.0001).

We profiled a total of 12,531 (7,581 cells with barcodes) melanoma cells and found two major transcriptionally distinct populations (Fig. 1D, Supp. Fig. 1A). We found that one cluster showed much higher expression of many of the genes associated with the primed resistant state including *EGFR* and *AXL* (Shaffer et al. 2020; Emert et al. 2021; Shaffer et al. 2017), and that most of the sorted NGFR and EGFR-high cells were included in this cluster (Fig. 1D, Supp. Fig. 1B). Of note, the two clusters were sufficiently distinct that they clustered separately even when the NGFR and EGFR enriched sample was not included in the analysis (Supp. Fig. 1C). Consistent with this observation, the other cluster showed higher expression of genes associated with the drug-susceptible state including *SOX10* and *MITF* (Fig. 1D, Supp. Fig. 1D).

Because primed cell states are reversible, we sought to leverage this reversibility to overcome the resistant cell fate. We used a pretreatment dosing strategy, in which we treated with a priming inhibitor prior to the main therapy (Fig. 1E). Here, the priming inhibitor forces cells out of the primed cell state and thus fewer primed cells are present in the population to become resistant to therapy. Importantly, this dosing scheme is possible because primed cells become resistant to therapy and priming is reversible (Torre et al. 2021; Shaffer et al. 2017). To identify a priming inhibitor, we used our scRNA-seq data and found that primed cells have increased expression of many growth factors (including *FGF1, VGF, BDNF, VEGFA, VEGFC*, and *PDGFA*) with convergence on the PI3K pathway (Stommel et al. 2007). We wondered whether blocking this common downstream PI3K signaling might inhibit the primed cell phenotype. We tested pretreatment with a PI3K inhibitor (PI3Ki), GDC-0941, for 5 days followed by BRAFi/MEKi for 4 weeks. Importantly, we selected this dose of PI3Ki as it did not kill the cells and had minimal effects on the growth rate (see description in Methods). Compared to BRAFi/MEKi alone, we found that PI3Ki pretreatment decreased the number of resistance colonies and the number of resistant cells (Fig. 1E representative images, Fig. 1F, G quantification across 6 wells and 3 replicates, Supp. Video 1, Supp. Video 2). Thus, using this dosing strategy informed by the priming model, we observe a significant decrease in drug resistance, suggesting that PI3Ki could be reducing the number of primed cells.

We next explored the primed cell gene expression state to better understand its regulation. Within the cluster of cells expressing primed cell genes, we noticed significant heterogeneity in previously described marker genes including *EGFR, AXL* and *NGFR* with each gene expressed by some of the cells in the cluster, but not all of the cells (Fig. 1D, Supp. Fig. 1D). Furthermore, in a separate experiment, we profiled the chromatin accessibility of EGFR and NGFR-high cells and found epigenetic differences between these populations further confirming that there is additional heterogeneity within the primed state (Supp. Fig. 1E). We suspected that our markers from previous work (Shaffer et al. 2017) did not capture all of the cells within this cluster. In order to capture all of the primed cells, we identified a novel marker, *NT5E*, that encapsulated the entire cluster containing the primed population (Fig. 1D). To validate *NT5E* as an effective marker of primed cells, we labeled WM989 cells with NT5E antibody and sorted the top 2% of cells. We then applied the targeted BRAF inhibitor, vemurafenib, and found that the NT5E samples had 5.5 fold more resistant cells compared to a mixed control (Supp. Fig. 1F, G). Thus, we demonstrated that the marker gene *NT5E* captures the entire cluster of transcriptionally similar cells and that these cells are indeed more resistant to targeted therapy. Surprisingly, despite the fact that our previous work showed that cells can switch between the drug-susceptible and primed state (Shaffer et al. 2020), there were no cells visibly transitioning between these populations in UMAP space. The lack of transitioning cells was also observed in a replicate WM989 scRNA-seq experiment (Supp. Fig. 2A).

With the scRNA-seq data alone, it is not possible to know whether cells are rapidly fluctuating between these states or whether the states persist at the single-cell level through multiple cell divisions. To extract the time that cells remain in these gene expression states, we used the lineage barcodes. First, we used the scRNA-seq data to assign all cells to one of two states, either drug-susceptible or primed for drug resistance. We then classified each lineage into one of four categories (Fig. 2A, B): 1. Drug-susceptible lineage (containing only drug-susceptible cells), 2. Primed lineage (containing only primed cells), 3. Switching from drug-susceptible to the primed state (initial state is drug-susceptible and lineage contains drug-susceptible and primed cells), or 4. Switching from primed to drug-susceptible state (initial state is primed and lineage contains drug-susceptible and primed cells). Across the classes of lineages, we found that lineage sizes were largest for drug-susceptible lineages and smallest for primed lineages, which is consistent with scRNA-seq predictions that primed cells divide slower than drug-susceptible cells (Fig. 2A, Supp. Fig. 2B). Given the high memory observed in both the drug-susceptible and primed states, we compared the differentially expressed genes between these states captured by scMemorySeq to bulk measurements of gene expression memory performed in the same cell line from (Shaffer et al. 2020). We observed a strong correlation between the two methods on a per gene basis (Supp. Fig. 2C). To derive the rates of proliferation and state switching from this data, we used a stochastic two-state model. Using the model we estimate that primed cells proliferate at approximately half the rate of drug-susceptible cells on average. We also estimate that drug-susceptible cells switch to the primed state once every 135-233 cell divisions, while primed cells are estimated to switch to the drug-susceptible state once every 5-8 cell divisions. The large difference in switching rates is due to the rarity of the primed cell state which must have a faster switching rate to maintain a constant proportion of primed cells at steady state. Thus, on the single-cell level, the drug-susceptible state is significantly more stable than the primed state (Supp. Methods 1).

**Figure 2:**
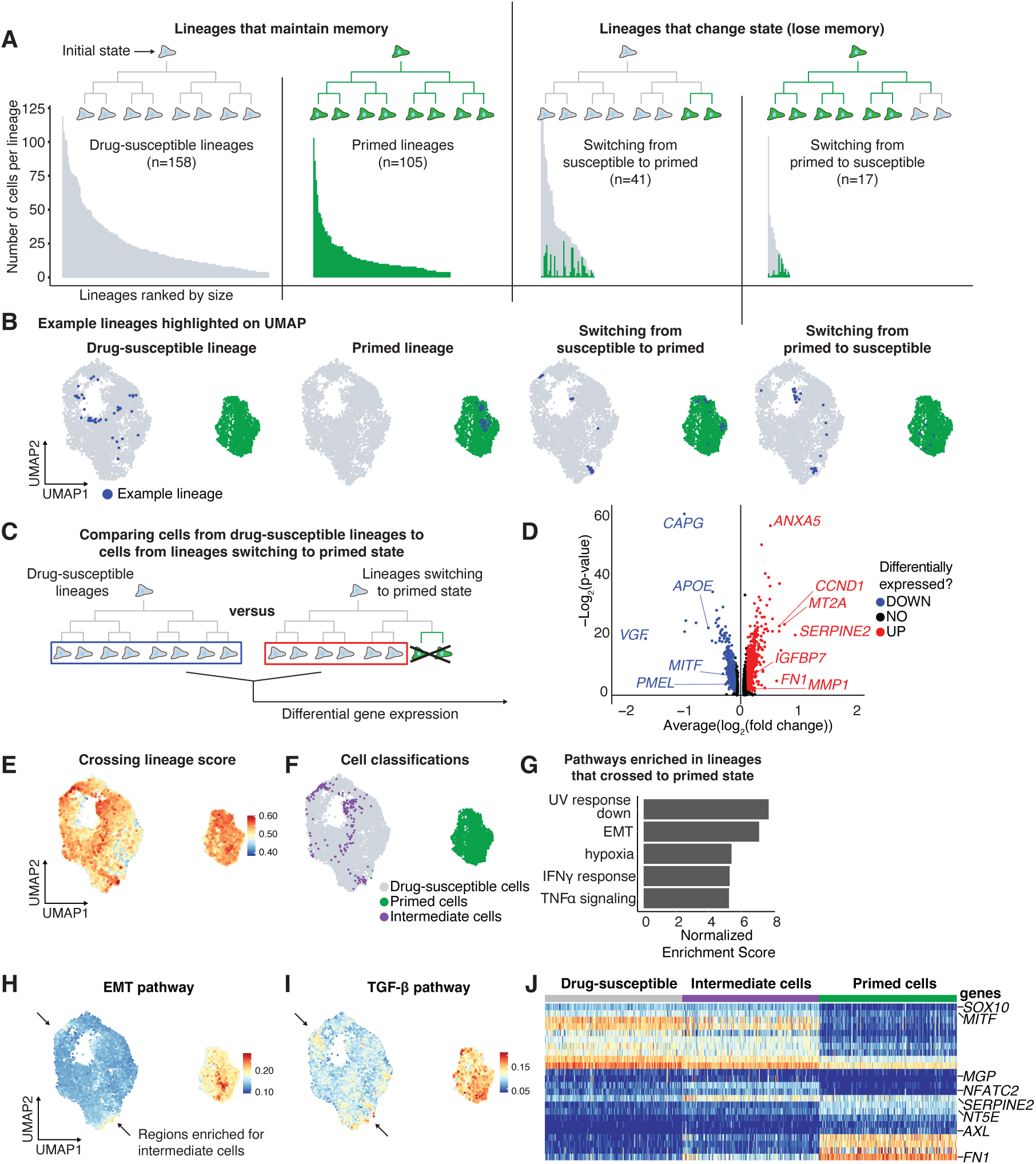
An EMT-like state is activated early in the transition to the primed cell state. **A**. Bar plots showing the size of each lineage found in the data organized based upon classification as drug-susceptible, primed, switching from susceptible to primed, or switching from primed to susceptible. Each bar represents an individual lineage, and the color of the bar indicates the state of the cell (green is primed and gray is drug-susceptible). In lineages that change state, the number of cells in each state is reflected by the colors in the bar. **B**. UMAP plots showing an example lineage from each type of lineage in the data. The cells from the example lineage are highlighted in blue. **C**. Schematic showing which cells from the drug-susceptible lineages (indicated in the blue rectangle) and which cells from lineages crossing from the drug-susceptible state to the primed state (indicated in the red rectangle) were used to identify differentially expressed genes in transitioning cells. **D**. Volcano plot representing the differential expression analysis outlined in C. Red points represent genes up regulated in crossing lineages and blue points represent genes downregulated in crossing lineages. **E**. UMAP plot showing an integrated score of the activity for the gene signature defined by crossing lineage differential expression in D (high score means the cell highly expresses the gene set) **F**. UMAP plot labeling the top 2% drug-susceptible cells expressing the crossing lineage gene set. We classify these cells as “ Intermediate cells”, and these cells are labeled in purple. **G**. Bar plot of the normalized enrichment score of the top 5 gene sets enriched in the crossing lineage gene set. **H**. UMAP plot showing which cells have high expression of the EMT pathway gene set. Arrows point to drug-susceptible cells with high EMT scores. **I**. UMAP plot showing which cells have high expression of the TGF-β signaling pathway gene set. Arrows point to drug-susceptible cells with high TGF-β signaling scores. **J**. Heatmap of the log10 normalized and scaled gene expression of cells in the drug-susceptible, intermediate, and primed state. Genese shown are the top differentially expressed genes between the primed and drug-susceptible states as well as select genes upregulated in the intermediate state.

After classifying the lineages based on whether they switch between states, we next wondered if there are transcriptional differences between lineages that undergo state-switching and lineages that do not switch. We hypothesized that these differences might exist if there was an intermediate transcriptional state as cells transition between the drug-susceptible and the drug-primed states. To uncover such a state, we compared lineages that contain only drug-susceptible cells to lineages that switch from drug-susceptible to the primed state (Fig. 2C). Importantly, this comparison requires the assumption that intermediate states would also demonstrate memory through cell division (schematic in Supp. Fig. 2D). Consistent with the presence of such an intermediate state, we found 575 genes differentially expressed between these types of lineages (based upon a differential gene expression analysis with a cutoff of 0.05 Bonferroni adjusted p-value and 0.25 log fold change cutoff) (Fig. 2D, Supp. Table 1). We then used this gene list to develop a “ crossing-lineage score”, applied this score to all cells in the data set, and then projected them into UMAP space (Fig. 2E). We identified two primary regions within the drug-susceptible cluster in UMAP space that show enrichment for cells in this intermediate expression state. We then set a threshold on this score and classified cells as “ intermediate cells” in addition to the two previously defined states, drug-susceptible and drug-primed (Fig. 2F). These intermediate cells are very rare, found at a frequency of around 2% in the population and would have not been identified using standard scRNA-seq techniques and analyses.

We next wanted to know what pathways are activated in the intermediate cell state. We first used gene set enrichment analysis on the differentially expressed gene list from Fig. 2D. The top pathways included UV response down, epithelial-to-mesenchymal transition (EMT), and response to hypoxia (Fig. 2G). We examined the activity of these different pathways in UMAP space and noted that the EMT pathway score localized in the same regions of the UMAP as the intermediate cell population (Fig. 2H). We focused on this gene set as multiple lines of evidence in melanoma suggest that an “ EMT-like” gene expression state is associated with resistance to BRAFi/MEKi (Ramsdale et al. 2015; Rambow et al. 2018; Verfaillie et al. 2015; Tang et al. 2020; Pedri et al. 2022). Another enriched pathway in our analysis was TGF-β signaling, which also showed enrichment in the same region of the UMAP as the intermediate cells (Fig. 2I). Furthermore, both the EMT and TGF-β pathways were further enriched in the primed cells beyond the levels in intermediate cells, suggesting that these are early changes as cells transition from drug-susceptible to intermediate to drug-primed (Fig. 2H, I). We also observed that these intermediate cells maintain expression of many genes associated with the drug-susceptible state, including melanocyte identity genes *SOX10*, and *MITF*, but have already begun to upregulate some of the important primed cell marker genes including *FN1* and *SERPINE2* (Fig. 2J). A few genes were uniquely expressed only in the intermediate state cells including *NFATC2*, and *MGP*. Intriguingly, *NFATC2* was previously found to be a regulator of *MITF* and melanoma dedifferentiation (Perotti et al. 2016; Aibar et al. 2017).

Having established these primed cell signatures in the WM989 cell lines, we next assessed their generalizability to tumor models and patient samples. We used RNA FISH HCRv3.0 on tumor samples derived from WM989 cells grown in NOD/SCID mice (Torre et al. 2021; Choi et al. 2018). We used probes for *NT5E* and *SOX10* on tumor sections and quantified expression across 5,600 cells. We found rare cells expressing high levels of *NT5E* mRNA scattered throughout the tissue (Fig. 3A, additional images in Supp. Fig. 2E). *SOX10* showed diffuse expression across the tissue, but many of the *NT5E*-high cells did not have *SOX10* expression, as predicted by the scRNA-seq. Altogether these *NT5E*-high *SOX10*-low cells demonstrate that the primed state exists both in vitro and in vivo (and is not an artifact of cell culture conditions). We also performed scRNA-seq on a different melanoma cell line, WM983B, and found a subpopulation population of cells with a large number of primed cell markers, including *NT5E* (Fig. 3B, C). To establish whether these cell states exist in patients tumors, we analyzed scRNA-seq data (Tirosh et al. 2016; Jerby-Arnon et al. 2018) which included 7 samples directly from human tumors and found that 5 of them had a subpopulation of cells with high expression of genes associated with the primed state (including the EMT and TGF-β gene signatures, Fig. 3D, E). Together, these data demonstrate the generalizability of the primed cell signatures as we observe them in mouse models and in patient samples.

**Figure 3:**
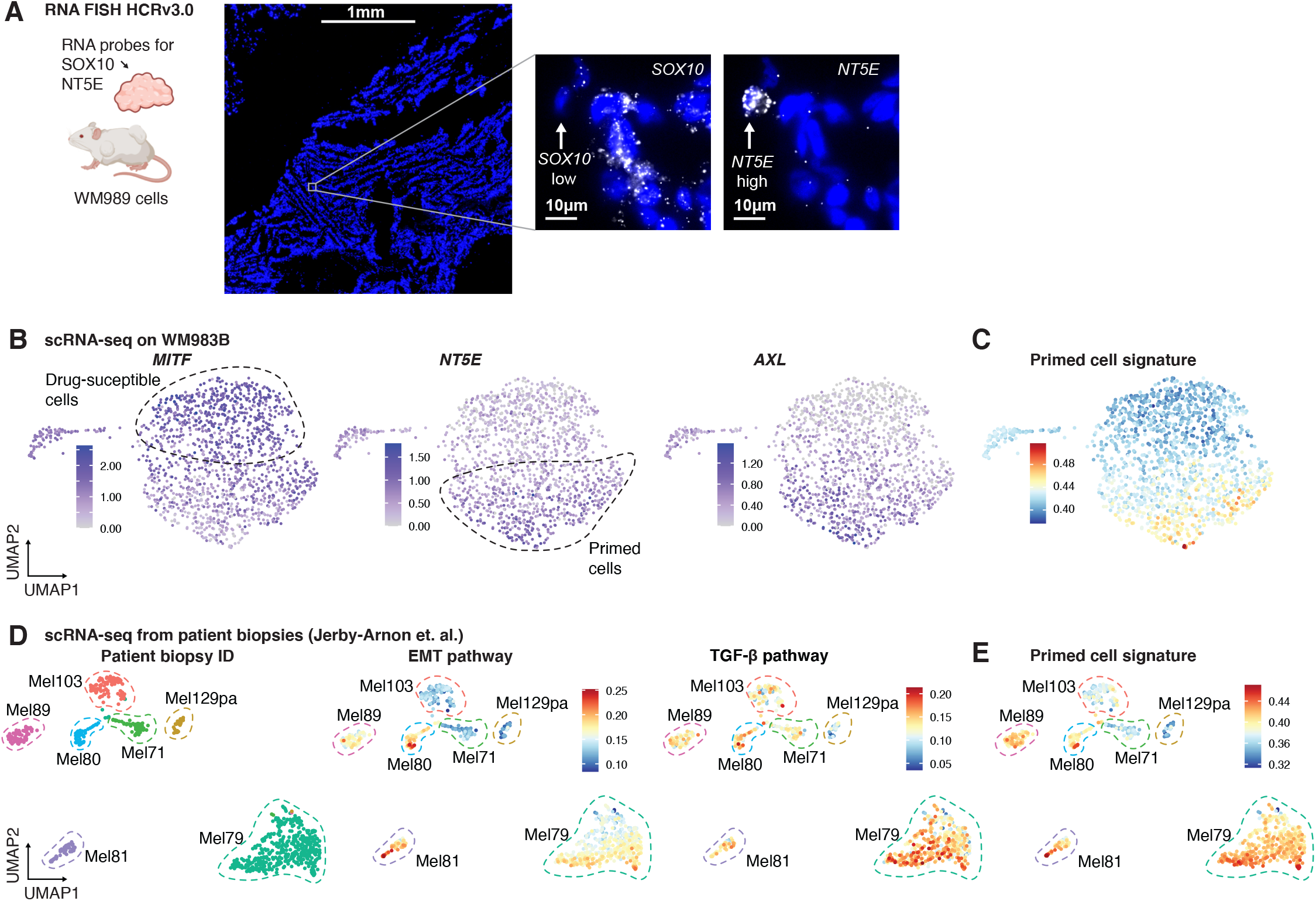
Mouse models and patient samples demonstrate expression of primed cell signatures. **A**. HCR RNA FISH of *NT5E* and *SOX10* in a mouse PDX drug naive tumor. Nuclei were stained with DAPI (blue) and RNA for each respective probe is in white. The green arrow points to an example *SOX10*-low *NT5E*-high cell. The leftmost image is a large scan of the tissue section and the scale bar corresponds to 1mm. The 2 images to the right are zoomed in on an example cell where the scale bar corresponds to 10µm. **B**. scRNA-seq on WM983B melanoma cell line. UMAP plots showing expression of *MITF, NT5E*, and *AXL* in each cell. **C**. UMAP showing WM983B scRNA-seq with each cell colored by the enrichment score for the primed cell signature derived from WM989 cells. **D**. Analysis of scRNA-seq of melanoma patient biopsies from (Tirosh et al. 2016; Jerby-Arnon et al. 2018). The first UMAP shows the different biopsies in different colors, and the other UMAP plots showing the signature score for gene sets associated with EMT and TGF-β signaling. Within each biopsy, there are cells that have variable expression of genes associated with EMT and TGF-β signaling. **E**. UMAP of patient scRNA-seq from D with the color of each point depicting the enrichment score for the primed cell signature.

Given the role of TGF-β as a potent inducer of EMT and the enrichment for the TGF-β pathway in the intermediate cells, we hypothesized that applying TGFB1 to drug-susceptible cells might drive them into the primed state (Fig. 4A) (Xu, Lamouille, and Derynck 2009). Importantly, we also observed that TGFB1 and its receptor are highly upregulated in primed cells suggesting that these cells naturally activate TGF-β signaling through autocrine or paracrine mechanisms (Supp. Fig. 2F). To test if TGFB1 induces the primed state, we treated melanoma cell lines including WM989 and WM983B with recombinant TGFB1 for 5 days and then performed flow cytometry for primed cell marker gene NT5E (Fig. 4B). We found that treatment with TGFB1 increased the percentage of cells of primed cells in WM989 from 1.98% to 19.15% and in WM983B from 10.04% to 81.56% (Fig. 4C, D). Our observed effects of TGFB1 on melanoma agree with literature showing that TGF-β signaling can induce a dedifferentiation state (Lee et al. 2020; Sun et al. 2014). The dedifferentiated state is similar to our primed cell population as these cells are also characterized by their decreased expression of melanocyte transcription factors SOX10 and MITF (Tirosh et al. 2016; Sun et al. 2014; Lee et al. 2020; Hoek et al. 2008). Furthermore, our work shows that these dedifferentiation events occur in a rare subpopulation of melanoma cells before targeted therapy whereas other studies focus on these gene expression states after treatment with targeted therapy.

**Figure 4:**
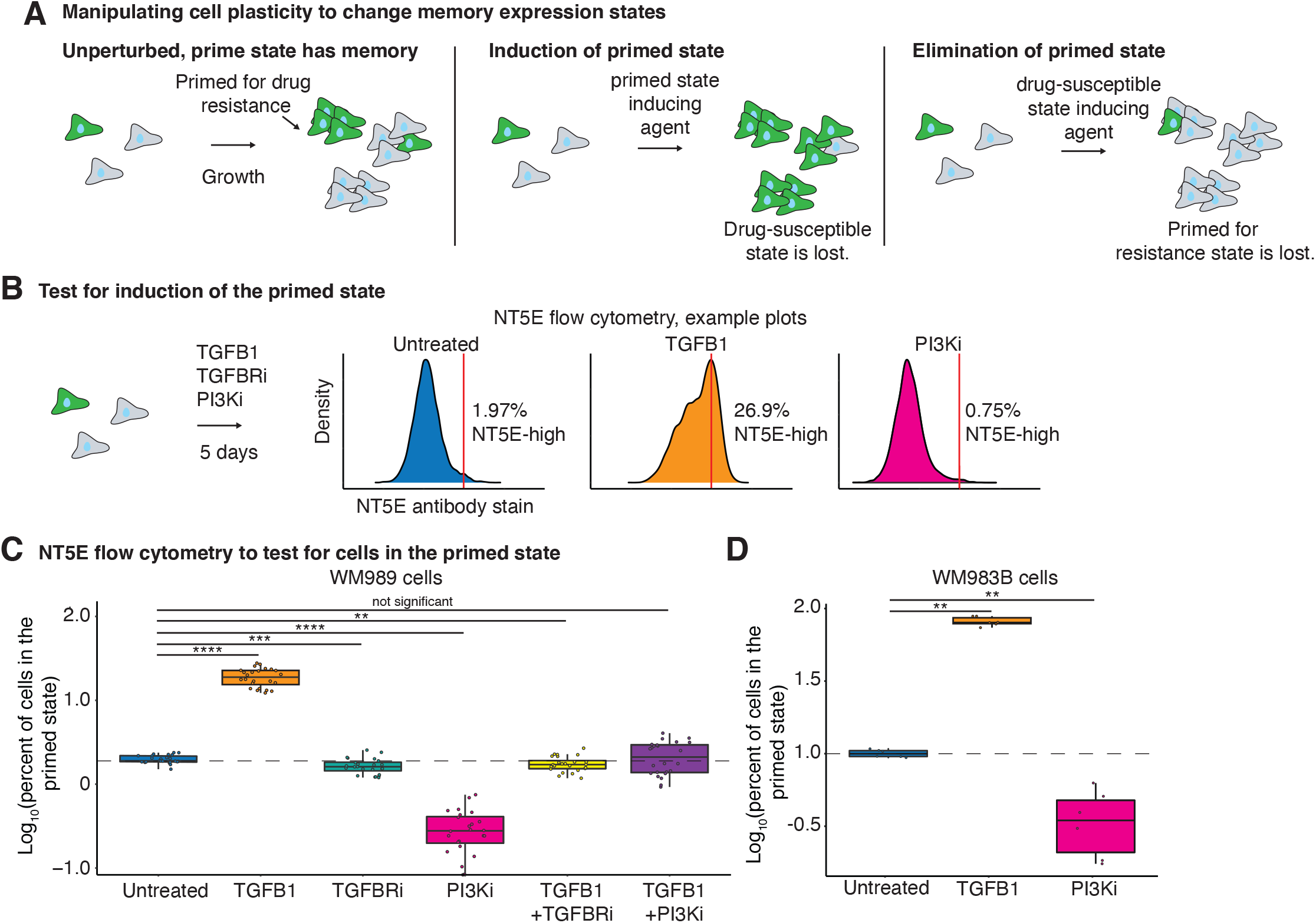
TGFB1 and PI3K inhibitor can modulate the number of primed cells in the population. **A**. Schematic showing the expected number of primed cells over time if cells are untreated, treated with a drug or protein that induces the primed state, or treated with a drug or protein that disrupts the primed state. **B**. Example flow cytometry density plots from NT5E stained WM989 cells after 5 days of their respective treatments. **C**. Box plot of the log_10_ percent of WM989 cells in the primed state after 5 days in each respective treatment. All conditions have 3 biological replicates each with 6 technical replicates. P values were calculated using a wilcoxon test (not significant: p > 0.05, **: p <= 0.01, ***: p <= 0.001, ****: p <= 0.0001). The dotted line represents the mean log_10_ percent of primed cells in the untreated condition. **D**. Box plot of the log_10_ percent of WM983B cells in the primed state after 5 days of each respective treatment. All conditions have 2 biological replicates each with 3 technical replicates. P values were calculated using a wilcoxon test (**: p <= 0.01). The dotted line marks the mean log_10_ percent of primed cells in the untreated condition.

Since TGFB1 was sufficient to induce the primed state, we wondered whether inhibiting TGF-β signaling would force cells in the opposite direction into the drug-susceptible state. We treated WM989 cells with LY2109761, a targeted TGFBR1/2 inhibitor (TGFBRi), for 5 days. Unexpectedly, we found that the TGFBRi did not change the percentage of primed cells significantly (Fig. 4C). To confirm the specificity of the inhibitor, we treated WM989 cells with both recombinant TGFB1 and TGFBRi and found that indeed the inhibitor was able to block the effects of TGFB1 (Fig. 4C). Overall this suggests that TGF-β signaling is sufficient to increase the percentage of primed cells, but is not necessary for maintaining them in the population.

To further test the functional effects of treating with TGFB1 and TGFBRi, we used the pretreatment experimental design (Fig. 1E) and applied BRAFi/MEKi following each treatment. Unexpectedly, we found that, although fewer total cells died, pretreatment with TGFB1 led to accelerated killing of the remaining drug-susceptible cells following BRAFi/MEKi compared to cells treated with BRAFi/MEKi alone (based on time-lapse microscopy, Supp. Fig. 3A-D). Furthermore, after 4 weeks of BRAFi/MEKi, we found that the pretreated TGFB1 sample had fewer resistant cells than the BRAFi/MEKi only sample. Interestingly, although there were fewer cells on the plate, they were much more evenly distributed suggesting that more of the initial cells survived BRAFi/MEKi, but that these cells were slow to proliferate. Consistent with its minimal effects on priming, we also found that TGFBRi did not change cell fate of resistance (Supp. Fig. 3A-C).

We also tested the effects of blocking PI3K on the primed state to determine whether the effects of PI3Ki on cell fate are mediated through changes in priming. We treated WM989 and WM983B cells with PI3Ki for 5 days and then performed flow cytometry for NT5E to quantify the percentage of primed cells. We found that the PI3Ki decreased the percentage of primed cells from 1.98% to 0.31% and 10.04% to 3.66% in WM989 and WM983B, respectively (Fig. 4C, D). Furthermore, we tested whether the PI3Ki is able to block the effects of TGF-β signaling by simultaneously treating WM989 cells with both TGFB1 and the PI3Ki. Indeed, we found that the PI3Ki was able to block the increase in primed cells seen when we treat with TGFB1 alone (Fig. 4C), suggesting that the TGFB1-mediated effects on priming melanoma cells acts through downstream PI3K activity.

Given that numerous growth factors were differentially expressed between the drug-susceptible and primed for resistance states, we wondered whether other ligands, in addition to TGFB1, might also be able to induce state switching. We treated WM989 cells with EGF, BDNF, and IL6 for 5 days (Supp. Fig. 3E) and found that none of these factors were able to induce the primed cell state as seen with TGFB1 despite their potential to increase PI3K signaling (Simiczyjew et al. 2019; Meng et al. 2019; Wegiel et al. 2007). Thus, we concluded that TGFB1 is a unique inducer of state switching and other growth factors do not have the same functionality.

Since TGFB1 and the PI3Ki were both able to change the percentage of cells in the primed state, we wondered how this is achieved at the single cell level. Specifically, does TGFB1 force more cells to switch into the primed state? Conversely, does the PI3Ki force cells to exit the primed state? Based on flow cytometry alone, we can conclude that the percentage of the population is shifted by these treatments; however, changes in multiple different parameters could lead to this same effect. For instance, the result that TGFB1 increases the percentage of primed cells could be explained by (1) an increase in growth rate of primed cells, (2) a selective killing of drug-susceptible cells, or (3) state-switching from the drug-susceptible into the primed state. The opposite consideration is necessary with the finding that the PI3Ki decreases the percentage of primed cells. This result could be the effects of (1) an increase in growth rate in the drug-susceptible cells, (2) a selective killing of primed cells, or (3) state-switching from the primed into the drug-susceptible state. To directly test how TGFB1 and PI3K shift the percentage of primed cells, we redesigned the scMemorySeq approach leveraging the knowledge of memory to test the effects of these perturbations. For this experimental design, we transduced WM989 cells with the high-complexity transcribed barcode library and then allowed the cells to go through 7-8 doublings (based upon experiments demonstrating that memory of these states persists on these timescales, Supp. Fig. 4A-C). We then split the cells into 4 separate plates and subjected each to a different condition (untreated, TGFB1 recombinant protein, TGFBRi, and PI3Ki) (Fig. 5A). Since cells within the same lineages have very similar gene expression states due to memory, this allows us to approximate studying how the “ same cell” will react to the different conditions. After 5 days, we harvested each sample and performed scRNA-seq and barcode-sequencing to capture the transcriptional state and barcode under each condition (Fig. 5A). With this experimental design, each barcode is represented across all the conditions, and thus, we can see the effects of that treatment by comparing the cells in each treatment to the cells in the untreated plate with the same barcode.

**Figure 5.**
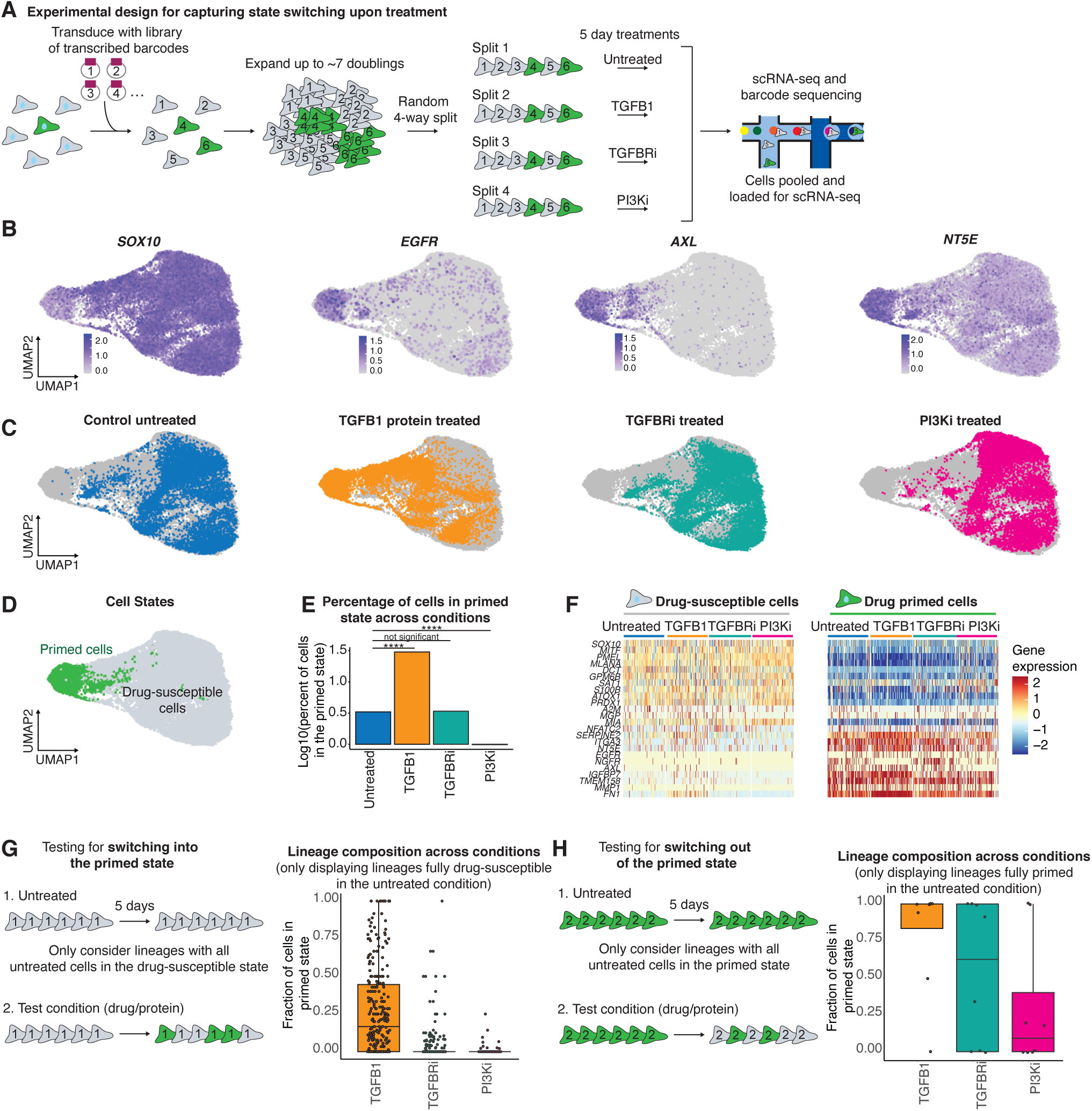
Treatment with TGFB1 induces the primed state and treatment with PI3K inhibitor induces the drug-susceptible state. **A.** Schematic of the experimental design used to determine if treatments cause state switching. We transduced melanoma cells (WM989) with lineage barcodes, allowed them to divide ∽ 7 times, and then split them across 4 plates, expecting each plate to get cells with the same barcode. Cells with the same barcodes serve as effectively as “copies“ of the same cells, therefore, by treating each barcode with all the different treatments we can determine which treatment causes state switching. We used scRNA-seq with barcode sequencing to capture both the lineage and transcriptome of the cells at the end point. **B.** UMAP plots of the log_10_ normalized gene expression of the drug-susceptible state associated marker *SOX10* and primed state associated markers *EGFR, AXL* and *NT5E*. **C.** UMAP plots highlighting cells in each condition relative to all the other sequenced cells (in gray). Cells highlighted in blue were untreated, cells highlighted in orange were treated with TGFB1, cells highlighted in teal were treated with TGFBRi, and cells highlighted in pink were treated with the PI3Ki. **D.** UMAP plot with the primed cells labeled in green and the drug-susceptible cells in gray. **E.** Bar plot quantifying the log_10_ percent of primed cells in each condition based on the defined drug-susceptible and primed cell populations in D and the location of each treatment condition shown in C. P values were calculated using chi squared tests (not significant: p > 0.05, ****: p <= 0.0001) F. Heatmaps of the log_10_ normalized and scaled gene expression of drug-susceptible and primed cells in each treatment condition. **G.** Schematic of lineage based analysis to test for state switching into the primed state. Box plots show the fraction of cells in each lineage that are in the primed state across all the conditions. The lineages shown on this plot are exclusively those that were completely drug-susceptible in the untreated sample. **H.** Schematic of lineage based analysis to test for state switching into the drug-susceptible state. Box plots show the fraction of cells in each lineage that are in the primed state across all the conditions. The lineages shown on this plot are exclusively those that were completely primed in the untreated sample.

We analyzed a total of 40,021 cells across the four different samples and found similar transcriptional states distributed across UMAP space as in previous experiments with *SOX10*-low cells clustered separately from cells with high expression of *EGFR, AXL*, and *NT5E* (Fig. 5B, Supp. Fig. 5A). Across the different treatment conditions, we found that the TGFB1 treatment shifted cells towards the primed state, while the PI3Ki shifted cells towards the drug-susceptible state (Fig. 5C). Consistent with previous experiments, the TGFBRi had minimal effects on the primed cell population. To classify cells as primed or drug-susceptible, we selected clusters 4 and 10 from the high-dimensional clustering in Seurat, as these clusters contained the majority of cells expressing the known primed cell marker genes (Fig. 5D, Supp. Fig. 5B, C). Based on this classification, we quantified the percentage of primed cells based upon the full transcriptional state captured by scRNA-seq and further confirmed the effects of TGFB1 and the PI3Ki (Fig. 5E). Importantly, the primed cell state induced by treatment with TGFB1 was transcriptionally very similar to untreated cells in the primed state (Fig. 5F). To explicitly test for differences, we performed differential gene expression to compare the TGFB1-induced and untreated primed cells. We found that some genes, including *NGFR, FGFR1, FOSL1*, and *JUN*, were induced by the TGFB1 treatment, but were not as highly expressed as in the untreated primed cells. This might be significant as *NGFR* expression has been linked to invasive properties of melanoma cells (Radke, Roßner, and Redmer 2017; Filipp, Li, and Boiko 2019). We also noted that TGFB1 seemed to induce even higher expression for many of the primed cell genes including *FN1, SERPINE1, COL1A1* and *VGF* (Supp. Fig. 5D). This effect might be explained by the high dose of TGFB1 used in these experiments relative to what these cells would generate on their own, which could lead to more dramatic changes in gene expression.

Next, we analyzed the barcoding data to test for state switching at the single cell level. Our data set included 19,740 lineages (containing a minimum of 3 cells per lineage) with 49% of lineages represented across all four conditions. To test whether TGFB1 induces more primed cells, we first evaluated lineages where the untreated sample had cells exclusively in the drug-susceptible state. Across each of these lineages, we quantified the percentage of primed cells in the other conditions and found that the matched set of lineages in the TGF B1-treated condition had a higher fraction of cells in the primed state (Fig. 5G). Reassuringly, the TGFBRi and the PI3Ki only had minor increases in the percentage of primed cells across these lineages. We next tested for the opposite direction, switching from the primed state into the drug-susceptible state. We analyzed lineages in which the entire lineage was in the primed state in the untreated sample. In the PI3Ki-treated sample, these lineages switched states leading to a lower percentage of primed cells (Fig. 5H). We noted that while 6 of the 8 lineages switched out of the primed state in the PI3Ki, 2 lineages did not respond to the PI3Ki. It is possible that non-responsive lineages would require a longer treatment with the PI3Ki to fully shift into the drug-sensitive gene expression state.

To extend our analysis to include all of the lineage data, we developed a stochastic model of state switching between the drug-susceptible and drug-primed state. Our model included different state switching parameters (k_on_, k_off_), different growth rates, and different death rates in each cell state (Supp. Fig. 5E). We used the model to simulate experimental data for different scenarios in which different parameters are changing. For instance, for TGFB1, the increase in primed cells could be explained by 3 possible parameter changes, 1) increase in K_on_ for the primed state, 2) increase in primed cell proliferation rate, or 3) increase in death rate among drug-susceptible cells. To constrain the proliferation rate parameter in our model, we performed live-cell imaging to measure the direct effects of 5 days of TGFB1 and the PI3Ki on the proliferation rates of primed and drug-susceptible cells. Across conditions, we found that the primed cells proliferate more slowly than the drug-susceptible cells (Supp. Fig. 5F). Additionally, we found that treatment with either TGFB1 or the PI3Ki decreased growth rate by similar amounts in each population (Supp. Fig. 5F).

Given these constraints on proliferation rates, we then ran 1 million simulations of the model varying each parameter and found that increasing the rate at which the cells switch into the primed state is the best fit for our data. Interestingly, the model also suggested that drug-susceptible cells in lineages with a high proportion of cells already in the primed state are more easily switched into the primed state (See Supp. Methods 1). We similarly considered possible parameter changes that could account for the decrease in primed cells upon PI3Ki treatment. Given experimentally measured constraints on proliferation rates, we found that an increased rate of cells switching from the primed state to the drug-susceptible state was the best fit for our data (See Supp. Methods 1). Taken together, this data shows that TGFB1 and the PI3Ki can induce state switching and that the observed changes in the number of primed cells are not due to other population dynamics.

## Discussion

Here we show that scMemorySeq is a powerful method for tracking gene expression memory in single cells. Our approach leverages the combination of scRNA-seq and cellular lineage barcoding to quantify memory of gene expression states in single cell data. We applied this method to melanoma cells to track lineages as they switch states between a drug-susceptible and a state primed for drug resistance. By analyzing the gene expression differences in lineages that switch states, we identified and tested TGF-β and PI3K as mediators of state switching at the single-cell level. Ultimately, we show that by manipulating state switching, we can reduce resistance to targeted therapy.

Broadly, it is intriguing that modulating signaling alone is sufficient to globally modify gene expression states in single cells and to affect their susceptibility to drugs. Here, PI3Ki is driving cells into a MAPK dependent transcriptional state, sensitizing cells to MAPK inhibitors. Importantly, a number of papers have reported the use of PI3Ki to reduce resistance in melanoma (Villanueva et al. 2010; Irvine et al. 2018; Mendoza, Er, and Blenis 2011; Nymark Aasen et al. 2019). Distinguishing our use of a PI3Ki, here, we find that the PI3Ki can reduce drug resistance even when only applied briefly before the addition of targeted therapy. This approach may lay out a generalizable strategy for reducing drug resistance in which perturbing signaling can globally tune gene expression to achieve susceptibility to drugs. In such a strategy, modulating signaling pathways would be used prior to the addition of the main targeted therapy to drive heterogeneous populations of cells into a drug susceptible state. This is in contrast to dosing with a combination of inhibitors at the same time, which is not always tolerated due to toxicity and side effects (Park et al. 2013; Rafsanjani Nejad et al. 2021; Jardim et al. 2020).

Conceptually, the idea of leveraging the plasticity of cells to revert them into a drug-susceptible state to delay drug resistance has previously been described in the context of drug holidays (Kavran et al. 2022). In contrast to our approach, in which we actively drive cells into a drug-susceptible state, a drug holiday is a break from the drug to alleviate the selective pressure thus giving the drug-resistant cells the opportunity to switch back to a drug-susceptible state at their intrinsic rate. Given the previous success of drug holidays (Das Thakur et al. 2013; Kavran et al. 2022), one strategy that we believe should be further explored is to use state switching drugs during what would be the holiday period. With this approach, the switch to a drug-susceptible state would be accelerated during the drug holiday, leading to potentially even less resistance. Future work is still needed to model these different scenarios and to experimentally test the efficacy of such dosing strategies.

Extending scMemorySeq beyond state switching in melanoma, there are a number of biological contexts in which specific drugs or ligands could be shifting gene expression states and population dynamics simultaneously. This is particularly relevant in cancer where there is a growing body of literature describing considerable heterogeneity at the single-cell level that shows variable degrees of memory (Tirosh et al. 2016; Barkley et al. 2021; Neftel et al. 2019). Here, we show that the scMemorySeq approach can provide a detailed systematic overview of these populations, and can also be used to test how closely related cells from the same lineage will react to different conditions. We believe that this approach is generalizable and could be used in other contexts to profile cell state transitions under different drugs, ligands, or environmental conditions. This is a growing area of interest as multiple recent studies showed that microenvironment and growth conditions of cancer cells can globally change both gene expression and sensitivity to drugs (Raghavan et al. 2021; Neal et al. 2018; Neftel et al. 2019).

In sum, we show how scMemorySeq uses cellular barcoding to reveal single-cell dynamics of drug resistance in melanoma. By tracking the memory of gene expression states we can identify stable cell populations as well as the factors that cause cells to change states. This approach can be widely applied to discover unknown dynamics in heterogeneous cell populations and to identify the key factors responsible for gene expression state changes in biological systems.

## Methods

### Antibodies

NGFR primary antibody (Biolegend, 345108), EGFR primary antibody (fisher scientific, MABF120MI), NT5E primary antibody (Biolegend, 344005), Alexa Fluor 488 Donkey Anti-Mouse secondary antibody (Jackson Labs, 715-545-151)

### Small molecule inhibitors and recombinant proteins

Vemurafenib (Selleckchem, S1267) was reconstituted at 4mM in DMSO for the stock solution. Dabrafenib (Cayman, 16989-10) was reconstituted at 1.25mM in DMSO for the stock solution. Trametinib (Cayman, 16292-50) was reconstituted at 12.5uM in DMSO for the stock solution. TGFB1 (R&D systems, 240-B-002) was reconstituted at 100ug/mL in a 4mM hydrochloric acid, 1mg/mL bovine serum albumin solution for the stock. PI3K inhibitor (GDC-0941, Cayman, 11600-10) was reconstituted at 10mM in DMSO for the stock solution. TGFBRi (LY2109761, SML2051-5MG) was reconstituted at 40mM in DMSO for the stock solution. IL6 (R&D systems, 206-IL-010) was reconstituted at 100ug/mL in a 0.1% BSA PBS solution for the stock. EGF (R&D systems, 236-EG-200) was reconstituted at 200ug/mL in PBS for the stock solution. BDNF (R&D systems, 206-IL-010) was reconstituted at 100ug/mL in a 0.1% BSA PBS solution for the stock. EGF (R&D systems, 248-BDB-005) was reconstituted at 100ug/mL in water for the stock solution.

### Cell lines and tissue culture

We used the following cell lines: WM989 A6-G3, which are a twice single-cell bottlenecked clone of the melanoma line WM989 (provided by the Meenhard lab at the Wistar Institute), WM983B E9-D5, which are a twice single-cell bottlenecked clone of the melanoma line WM983B (provided by the Meenhard lab at the Wistar Institute), and HEK293FT cells which we used for lentiviral packaging. We authenticated the identity of all cell lines by STR profiling and confirmed that they are all negative for mycoplasma. STR profiling and mycoplasma testing was performed by the Penn Genomic Analysis Core. We cultured WM989 A6-G3 and WM983B E9-D5 in TU2% (78.4% MCDB 153, 19.6% Leibovitz’s L-15, 2% FBS, 1.68mM CaCl, 50 Units/mL penicillin, and 50µg/mL streptomycin). We cultured HEK293FT in DMEM 5% (95% DMEM high glucose with GlutaMAX, 5% FBS, 50 Units/mL penicillin, and 50µg/mL streptomycin). We grew all cells at 37°C and 5% CO_2_, and passaged them using 0.05% trypsin-EDTA.

### Barcode library

We used a high-complexity transcribed barcode library described in Emert et al. for our lineage barcodes (Emert et al. 2021). The plasmid uses LRG2.1T as a backbone, but we replaced the U6 promoter and sgRNA insert with GFP followed by a 100 nucleotide semi-random barcode, expressed by an EFS promoter. The barcode is semi-random as it is made up of WSN repeats (W = A or T, S = G or C, N = any) to maximize barcode complexity. A detailed protocol on the barcode production process can be found in Emert et al. which also links to this protocol: https://www.protocols.io/view/barcode-plasmid-library-cloning-4hggt3w. The sequence of the plasmid can be found here: https://benchling.com/s/seq-DAMUWPyU198hRSbpiecf?m=slm-GJ609ijArVWmkT8mk8zr

### Lentiviral packaging

We grew HEK293FT to about 90% confluence in a 10cm dish containing 10mL of media (see the “ cell lines and culture” section for details). To transfect cells with the lentiviral plasmid, we combined 500µl of OPTI-MEM with 80µl of 1mg/mL PEI in one tube. In a second tube, we combined 500µL of OPTI-MEM, 9 µg of the psPAX2 plasmid, 5.5µg of the VSVG plasmid, and 8µg of the barcode plasmid. We combined the contents of these two tubes and allowed the mixture to incubate at room temperature for 15 minutes. We then pipetted the solution dropwise into the plate of HEK293FT and incubated the cells at 37°C for 7 hours. Next, we removed the media, washed the plate once with DPBS, added 10mL of fresh media, and then incubated the cells at 37°C for ∼12 hours. We used fluorescence microscopy to confirm GFP expression in the cells and then applied a fresh 6mL of media for virus collection. We incubated the cells at 37°C for ∼12 hours and collected the media (this media contains the virus). We repeated the process of adding 6mL of media and collecting virus every 12 hours for a total of ∼72 hours. After the last collection, we filtered all the media containing the virus through a 0.2µm filter to ensure no HEK293FT cells were left with the virus media. Finally, we made 1mL aliquots of the media containing the virus and stored them at -80°C.

### Lentiviral transduction

When barcoding cells, we wanted to avoid multiple lineage barcodes per cell, and thus, we aimed to transduce ∼20% of cells. To transduce the cells, we made a mixture of polybrene (4µg/mL final concentration), virus (concentration determined through titration experiments to achieve 20% infection), and cells at 150,000 cells/mL. Next, we put 2mL of this mixture into each well of a 6-well plate and spun the plate at 600 RCF for 25 minutes. We then incubated the cells with the virus at 37°C for 8 hours. After the incubation, we removed the media containing the virus, washed each well with DBPS, and added 2mL of fresh media to each well. The next day, we transferred each well to its own 10cm dish. We then gave the cells 2-3 days to start expressing the barcodes. We confirmed the presence of barcodes by GFP expression in the cells (cells express GFP along with the barcode).

### Fluorescence-Activated Cell Sorting (FACS)

We dissociated cells into a single cell suspension using trypsin-EDTA and washed them once with 0.1% BSA. To stain for EGFR and NGFR, we first stained with the EGFR antibody (see antibodies section) diluted 1:200 in 0.1% BSA for 1 hour on ice. We then washed the cells twice with 0.1% BSA and stained with the anti-mouse A488 secondary antibody at a 1:500 dilution in 0.1% BSA for 30 minutes on ice. Next, to stain for NGFR, we washed the cells once with 0.1% BSA, and then resuspended in a 1:11 dilution of the NGFR antibody directly conjugated to APC in 0.5% BSA 2mM EDTA solution. We then incubated the cells on ice for 10 minutes. Finally, we washed the cells once with 0.5% BSA 2mM EDTA, resuspended in 1% BSA with DAPI, and kept the cells on ice until sorting.

To stain for NT5E, we resuspend the cells in a solution of NT5E antibody diluted 1:200 in 0.1% BSA and let the cells incubate for 30 minutes. We then washed the cells twice with 0.1% BSA, resuspended the cells in 1% BSA with DAPI, and kept on ice until sorting.

For flow sorting, we followed the staining protocols above and then sorted the cells on a Beckman Coulter Moflo Astrios with a 100µm nozzle. We used forward and side scatter to separate cells from debri and select singlets. We selected DAPI negative cells to remove dead cells. To sort primed cells with EGFR and NGFR, we selected the top 0.2% of EGFR and NGFR expressing cells. To sort primed cells with NT5E, we selected the top 2% of NT5E expressing cells.

### Single-cell RNA sequencing

We used the 10x Genomics 3’ sequencing kits for all our scRNA-seq experiments. For the first scRNA-seq experiment, introduced in Fig. 1C, we sorted 1,000 barcoded WM989 cells per well in a 96-well plate (one well of mixed cells and one of EGFR/NGFR-high cells) and allowed them to expand through 4-5 doublings. We then trypsinized one mixed well and one primed well, and prepared as described in the Chromium Single Cell 3’ Reagent Kit V3 user guide. When loading cells on the microfluidic chip, we split both the primed and the mixed cells across 2 wells. After GEM generation, we continued to follow the Chromium Single Cell 3’ Reagent Kit V3 user guide to generate libraries.

For the second scRNA-seq experiment (data shown in Supp. Fig. 2A, B), we sorted 1,000 WM989 primed cells (based on NT5E expression) and 1,000 mixed WM989 cells into one well of a 96-well plate, and 2,000 WM989B cells into another well. We then allowed the cells to undergo 4 divisions, trypsinized them, and prepared the samples all the way through library generation as described in the Chromium Single Cell 3’ Reagent Kit V3.1 (Dual Index) user guide.

For the third scRNA-seq experiment, shown in Fig. 5A, we sorted 2,000 barcoded WM989 cells into a single well of a 96-well plate. We waited for these cells to expand through 7 to 8 divisions, and then randomly split these cells across 4 separate plates. We waited one day for the cells to adhere to the plate, and then started treatments (one plate untreated, one plate 5ng/mL TGFB1, one plate 4µM LY2109761 (TGFBRi), and one plate 2µM GDC-0941 (PI3K inhibitor)). We incubated the cells in their respective treatments for 5 days. We carried the above steps with two samples in parallel as replicates. After 5 days, we trypsinized the cells and processed them all the way through library generation as described in the chromium Single cell 3’ Reagent kit V3.1 (Dual Index) user guide.

We sequenced all our single-cell libraries using a NextSeq 500 with the High Output Kit v2.5 (75 cycles, Illumina, 20024906). For samples sequenced with the Single Cell 3’ Reagent Kit V3 (single index), we used 8 reads for the index, 28 reads for read 1, and 49 reads for read 2. For samples sequenced with the Single Cell 3’ Reagent Kit V3.1 (dual index), we used 10 cycles for each index, 28 cycles for read 1, and 43 cycles for read 2.

### Lineage barcode recovery from scRNA-seq

To recover the lineage barcodes, we used an aliquot of the excess full length cDNA generated in the 10x library protocol. Specifically, we selectively amplified reads containing the lineage barcode using primers that flank the 10x cell barcode and the end of the lineage barcode in our library (Supp. Table 2) (Goyal et al. 2021). To perform the PCR, we combined 100ng of full length cDNA per reaction, 0.5µM of each primer, and PCR master mix (NEB, M0543S). We used 12 cycles to amplify the cDNA using the following protocol: an initial 30 second denature step at 98°C, then 98°C for 10 second followed by 65°C for 2 minutes repeated 12 times, and a 5 minute final extension step at 65°C. We then extract the amplified barcodes, which are ∼1.3kb, using SPRI beads (Becman Coulter, B23317) size selection (we use a 0.6X bead concentration). To sequence the barcode library, we used a NextSeq 500 with a Mid Output Kit v2.5 (150 cycles, Illumina, 20024904). We performed paired-end sequencing and used 28 cycles on read 1 to read the 10x barcode and UMI, 8 cycles on each index, and 123 cycles on read 2 to sequence the lineage barcode.

### gDNA barcode recovery

To sequence barcodes from gDNA, we trypsinized cells, pelleted them, and then extracted their gDNA using the QIAamp DNA Mini kit according to the manufacturer’s protocol (Qiagen, 56304). To amplify the barcodes, we performed PCR using primers with homology to each side of the barcode. The primers also contain the illumina adapter sequence, and index sequences (see Supp. Table 2 for primer sequences). To perform the PCR amplification, we used 500ng isolated gDNA, 0.5µM of each primer, and PCR Master Mix (NEB, M0543S) for each reaction. We used 24 cycles to amplify the barcodes using the following protocol: an initial 30 second denature step at 98°C, then 98°C for 10 second followed by 65°C for 40 second repeated 24 times, and a 5 minute final extension step at 65°C. After amplification, we used SPRI beads (Becman Coulter, B23317) to select for the amplified barcode product (expected length of ∼350bp). To isolate this fragment size, we performed a two sided selection where we first selected with a 0.6 X bead concentration and kept the supernatant (large gDNA fragments were on beads). We then select again using a 1.2X bead concentration keeping the material bound to the beads(small fragments such as the primers were in the supernatant). To sequence the barcode library, we used a NextSeq 500 with a Mid Output kit (150 cycles, Illumina, 20024904). We performed single -end sequencing and used 151 cycles on read 1 to read the lineage barcode and 8 cycles on each index.

### scRNA-seq analysis

We used the 10x Genomics Cell Ranger pipeline to generate FASTQ files (using the hg38 reference genome), to assemble the count matrix, and to aggregate replicate runs (without depth normalization). We also used the Cell Ranger feature barcode pipeline to integrate our lineage barcodes with the scRNA-seq data (more information in the “ Combining single RNA sequencing and barcode data section”).

Once we generated the aggregated count matrices with incorporated barcode information, we analyzed the data using Seurat V3 (Stuart et al. 2019). Using Seurat, we performed basic filtering of the data based on the number of unique genes detected per cell, both removing poorly sequenced cells (low number of genes), and data points likely to be doublets (high number of genes). We also filtered based on the percent of mitochondrial reads to eliminate low quality or dying cells. If we saw batch effects between replicates, we used the Seurat scRNA-seq integration pipeline to remove them. When there were no batch effects, we used SCtransform to normalize the data before running PCA, clustering, and dimensionality reduction with UMAP. When plotting gene expression information, we did not use the SCtransform data, but rather separately log normalized the data. To generate single-cell signature scores for a gene set, we used the UCell package (Andreatta and Carmona 2021). We selected the primed cell gene set by including all genes with a positive log_2_ fold change in our list of differentially expressed genes between primed and drug-susceptible cells (Supp. Table 1)

### Combining scRNA-seq and barcode data

To identify the lineage barcodes from the sequencing data, we used a custom python script (available through github here: https://github.com/SydShafferLab/BarcodeAnalysis) and the 10x Genomics Cell Ranger Feature Barcode pipeline. In this pipeline, we first identified lineage barcodes in the FASTQ files by searching for a known sequence at the beginning of all lineage barcodes. Once we identified all potential barcode sequences, we used the STARCODE (Zorita, Cuscó, and Filion 2015) to identify barcodes that were very similar to each other and replace them all with the most frequently detected sequence within the set of similar barcodes. We then put these modified barcode sequences back into the FASTQ file and generated a reference file containing all the edited barcode sequences. Next, we fed these edited FASTQ files and the reference file into the Cell Ranger pipeline, and used the Feature Barcode analysis function to link lineage barcodes with the cell barcodes. This provided us with the lineage and gene expression information for cells where a barcode was identified.

Our initial steps identified barcodes by combining similar barcodes, but when we looked at this output we found that we could more stringently call real lineages using additional filtering steps. The code used to accomplish this can be found in the “ Assign a lineage to each cell” section of the 10×1_r1_r2_Analysis_unorm_sctrans.Rmd script available on the Google Drive link in the Software and data availability section. In brief, we first eliminated lineages that appear across multiple samples, as such lineages are not possible. We then also removed lineages that are bigger than expected given the amount of time cells were given to proliferate. Finally, for cells that appeared to have more than one lineage barcode, we tested whether there are multiple cells with this same combination of barcodes and considered those cells to be in the same lineage.

### Validating primed cell markers

To test whether different proteins are markers of the primed cell state (NT5E, NGFR, EGFR), we stained live WM989 cells with an antibody for the marker of interest, and then sorted the stained cells. Specifically, we sorted a mixed population of cells in one well, and a population of cells high in our marker of interest in another well (for sorting detail refer to the “ Fluorescence-Activated Cell Sorting” section). We then allowed the cells to adhere to the plate for 24 hours and then started treatment with 1µM vemurafenib for 3 weeks. After 3 weeks in vemurafenib, we counted the number of cells and drug resistant colonies in each well to determine if the marker increased the number of cells that survive targeted therapy. The percentage of cells sorted in the primed condition was determined by sorting different percentages of high cells and treating them with targeted therapy to identify which percentages were resistant.

### ATAC-seq

We sorted 10,000 cell populations of EGFR/NGFR-High, EGFR-High, NGFR-High, and negative (for both markers) cells in triplicate as described in the FACS section of methods. Immediately after sorting, we performed OMNI-ATAC on each population of cells (Corces et al. 2017). We used the Illumina Tagment DNA Enzyme for tagmentation (Illumina 20034197) for tagmentation and performed two-sided bead purification before sequencing. We performed paired-end, single-index sequencing on pooled libraries using a 75-cycle NextSeq 500/550 High Output Kit v2.5 (20024906) allotting 38 cycles to both read 1 and read 2 and 8 cycles to the sample indices.

### ATAC-seq alignment and analysis

We adapted the paired-end analysis pipeline from (Sanford et al. 2020) for alignment, processing, and peak calling. Briefly, we aligned reads to hg38 using bowtie2 v2.3.4.1, filtered out low quality read alignments using samtools v1.1, removed duplicated reads with picard 1.96, and generated alignment files with inferred Tn5 insertions. To call peaks, we used MACS2 2.1.1.20160309. We then identified consensus peaks using the “ findConsensusPeakRegions” function in the consensusSeekeR package in R as peaks seen in at least 3 replicates out of 12 total (*consensusSeekeR: Bioconductor Package - Detection of Consensus Regions inside a Group of Experiments Using Genomic Positions and Genomic Ranges* n.d.). We then counted reads within these consensus regions for each sample and created a DESeq2 object which we used to perform PCA on consensus peaks (Love, Huber, and Anders 2014). We then plotted a row scaled heatmap with “ ward.D2” clustering of the top 20,000 most variable peaks.

### Mouse model tumor generation

The tissue was collected by the Weeraratna lab (Torre et al. 2021). Briefly, these melanoma tumors were generated by subcutaneously injecting 1*10^6 WM989-A6-G3-Cas9-5a3 cells into 8-week old NOD/SCID mice. The mice were fed AIN-76A chow, and the facilities were maintained between 21-23°C, a humidity of 30-35%, and lights had a 12h on/off cycle with lights on from 6:00 to 18:00. The tumor was collected when it measured ∼1,500 mm^3^. Tumor blocks were embedded in OCT, flash frozen, and stored at -80°C.

### Tissue RNA FISH

To analyze *NT5E* and *SOX10* expression in mouse tumors, we used HCRv3.0 with probes targeting *NT5E* and *SOX10* (Acheampong et al. 2022; Choi et al. 2018). The probes and fluorescently labeled hairpins were purchased from Molecular Instruments (*NT5E* lot #: PRK825, *SOX10* lot #: PRK826). To perform HCR in tissue, we made slight modifications to published protocols (Choi et al. 2018; Acheampong et al. 2022). First, we used cryostat sectioning to generate 6µm sections of fresh frozen tumor. We placed these sections on charged slides and fixed them with 4% formaldehyde for 10 minutes. We then washed the slides twice with 1X PBS and stored them in ethanol.

To start HCR, we placed the slide in a slide staining tray and washed the slides twice with 5X SSC (sodium chloride sodium citrate). After removing the 5X SSC, we added 200µl of hybridization buffer (30% formamide, 5X SSC, 9mM citric acid (pH 6.0), 0.1% Tween 20, 50ug/mL heparin, 1X Denhardt’s solution, 10% dextran sulfate) which was pre-heated to 37°C onto the tissue. We then incubated the slide for 10 minutes at 37°C. All incubation steps in this protocol were done with the slide staining tray closed and with water at the bottom of the tray to prevent the sample from drying out. During this incubation period, we added 0.8 pmol of each probe pool (in this case *NT5E* and *SOX10*) to 200µl of probe hybridization buffer pre-heated at 37°C, and kept the solution at 37°C. After 10 minutes, we removed the hybridization buffer from the tissue and added 300µl of hybridization buffer containing the probe pools. We placed a cover slip over the sample and incubated it for 12 -16 hours at 37°C. After the incubation, we prepared the hairpins by putting 0.6pmol each hairpin into its own tube (to keep hairpin 1 and hairpin 2 separate), and then performed snap cooling by heating them to 95°C for 90 seconds and then slowly cooled down to 25°C over 30 minutes in a thermocycler. While the hairpins snap cooled, we added 300µl of wash buffer (30% formamide, 5X SSC, 9mM citric acid (pH 6.0), 0.1% Tween 20, 50ug/mL heparin) to the slide to remove the cover slip. We then performed multiple wash steps with decreasing amounts of wash buffer in the solution. We first added 300µl of 75% wash buffer, 25% 5X SSCT (5X SSC with 0.1% Tween 20), removed that and added 300µl of 50% wash buffer, 50% 5X SSCT, removed that and added 25% wash buffer, 75% 5X SSCT, and finally removed that and added 300µl of 100% 5X SSCT. For each step of the wash, we left the slides in solution for 15 minutes at 37°C. After the last wash, we removed the 5X SSCT and added 200µl of room temperature amplification buffer (5X SSC, 0.1% Tween 20, 10% dextran sulfate) to the slide and incubated it at room temperature for 30 minutes. We then removed the amplification buffer and added the prepared hairpins mixed in 100µl of amplification buffer to the slide, and added a cover slip. We incubated the slide in the staining tray at room temperature for 12-16 hours. After incubating with the hairpins, we removed the cover slip and washed the slide off using successive 5X SSC washes. We put 300µl of 5X SSC on the sample for 5 minutes, removed it, then added 5X SCC for 15 minutes, removed it, added 5X SSC for 15 minutes again, removed it, and finally added 5X SSC with DAPI for 5 minutes. After the washes, we mounted the slide using TrueVIEW (Vector labs, SP-8500-15), added a coverslip, and sealed with nail polish.

### Flow cytometry

We dissociated cells from the plate using trypsin-EDTA into a single-cell suspension and washed once with 0.1% BSA. We then resuspended the cells in a 1:200 dilution of anti-NT5E antibody conjugated with APC and incubated them for 30min on ice. Next, we washed the cells once with 0.1% BSA, once with 1% BSA, and then resuspended them in 1% BSA for analysis by flow cytometry. We used an Accuri C6 for our flow cytometry, and quantified 10,000 events per sample. To analyze the data we used the R package flowCore (Hahne et al. 2009). In our analysis we used forward and side scatter to identify cells, and used the FL4 channel (640nm excitation laser and 675/25 filter) to quantify cell surface levels of NT5E. To determine what percent of cells were primed, we set an intensity threshold where 2% of untreated cells would land above the threshold. We considered any cells above this threshold as primed.

### Drug resistant colony experiments

We plated cells in 6-well plates with 10,000 cells per well. After plating, we gave cells 24 hours without treatment to adhere to the plate. We then initiated pretreatments and changed the media on the no pretreatment controls. During the pretreatment period, we treated the cells with doses of the drug that had extremely low toxicity and minimal effect on the proliferation rate of the cells (assay for determining the doses described in the “ pretreatment growth effects” section). We incubated cells in their respective pretreatment for 5 days. We then aspirated the media and replaced it with media containing 250nM dabrafenib and 2.5nM trametinib. We maintained treatment with 250nM dabrafenib and 2.5nM trametinib for 4 weeks, changing the media every 3 -4 days. After 4 weeks, we fixed the cells by aspirating off the media, washing the wells with DPBS, and treating them with 4% formaldehyde for 10 minutes. We then aspirated off the formaldehyde, and washed twice with DBPS. Finally, we added 2mL of DPBS to each well, and stained the cells with DAPI. We then imaged the wells using a 10X objective on a fluorescence microscope (Nikon, Eclipse Ti2).

### Cell and colony counting

To determine how many cells were in wells after drug treatment, we used a custom pipeline called DeepTile (https://github.com/arjunrajlaboratory/DeepTile/tree/071e3e9fb27f50ce024fd5ece25e3a4b0071f771) to feed tiled images into DeepCell to generate nuclear masks (Bannon et al. 2021; Greenwald et al. 2022). To simplify the interface with DeepTile and DeepCell, as well as remove nuclei incorrectly called outside the well, we used a custom tool DeepCellHelper (https://github.com/SydShafferLab/DeepCellHelper). We then determined the number of cells per well by counting the number of masks per image. To identify colonies, we used a custom graphic user interface ColonySelector (https://github.com/SydShafferLab/ColonySelector) to circle individual colonies in each well and save a file containing which nucleus belongs to which colony. Using the output of the colony selecting software, we counted the number of colonies there were in each well.

### IncuCyte imaging and analysis

For time-lapse experiments on the IncuCyte S3 (Sartorius), we used a clonal population of WM989 cells tagged with H2B-GFP for nuclear tracking. We took 4X images with a 300ms exposure for GFP every 12 hours to track cell growth over time. We used the IncuCyte software to generate nuclear masks and exported csv tables containing the number of nuclei in each well at each time point. We analyzed this data in R.

### Pretreatment growth effects

To determine if our treatments were leading to state specific changes in proliferation rates, we used a clonal WM989 H2B-GFP tagged cell line. To isolate drug-susceptible and primed cells, we sorted on NT5E and separated drug-susceptible and primed cells into separate wells. After 24 hours to adhere to the plate, we added TGFB1, PI3Ki, or nothing to the media. We then imaged the cells according to the description in the “ Incucyte imaging and analysis” section.

## Supporting information

Supplemental Video 1 (untreated)

Supplemental Video 2 (PI3Ki pretreated)

## Software and data availability

All data and code used for this paper can be found here: https://drive.google.com/drive/folders/1-C78090Z43w5kGb1ZW8pXgysjha35jlU?usp=sharing

## Author Contributions

G.H. and S.M.S. conceptualized the project and designed the study. G.H. performed all experiments and analysis with the following exceptions. R.A.R.H. helped with the barcode processing pipeline and provided helpful discussion. D.S. performed the ATAC sequencing experiment and its analysis, and helped with the barcode processing pipeline. M.S. and A.S performed the modeling analysis. B.E. provided technical guidance and troubleshooting for the barcoding library. A.T.W, M.E.F, and G.M.A generated the mouse PDX tissue. S.N. helped maintain cell lines and performed computational analyses. G.H. and S.M.S. wrote the paper.

## Acknowledgements

We thank all members of the Shaffer lab for feedback on experiments and the manuscript, P. Gonzalez-Camara and T. Ridky for thoughtful discussions and ideas, W. Niu for help on developing image processing pipelines, L. Bugaj for feedback on the manuscript, and M. Herlyn for providing cell lines. S.M.S. acknowledges support from the NIH Director’s Early Independence Award DP5OD028144 and the Wistar/Penn Skin Cancer SPORE (P50 CA174523). A.S. acknowledges support from R01GM124446. A.T.W. acknowledges support from R01CA207935 and P01CA114046.

