## Supplemental Figures for "Disrupting cellular memory to overcome drug resistance"

### Supplemental Figure 1

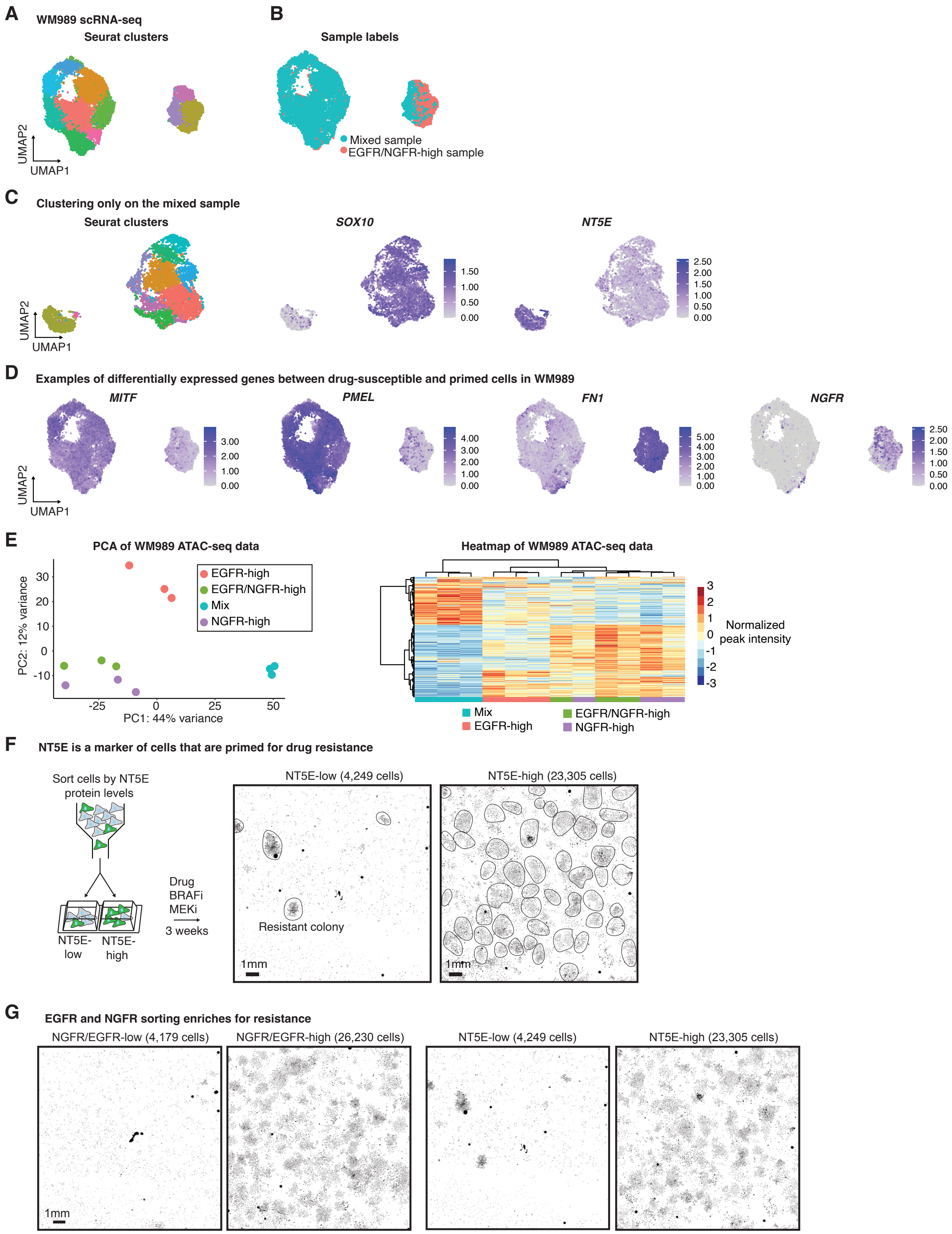

**Supplementary Figure 1: Additional characterization of primed cells by scRNA-seq, ATAC-seq, and resistance phenotype. A.** Seurat clusters of drug-naive WM989 cells displayed in UMAP space. **B**. UMAP plot showing sample labels from experiment in Fig. 1C. Each cell is labeled either from the mixed sample or from the sample that was enriched for EGFR/NGFR-high cells. The sample that was enriched for EGFR/NGFR-high cells are significantly enriched in the right cluster containing the primed cells. **C.** UMAP dimensionality reduction plots including only the mixed sample of cells (no enrichment for primed cells). Without including the sample that is enriched for primed cells, the drug-susceptible and primed cells (identified by *SOX10* and *NT5E* respectively) still separate into distinct clusters. **D.** UMAP plots showing normalized gene expression of the indicated gene for each cell. We find that melanocyte markers *MITF* and *PMEL* are high in the drug-susceptible cells, while *NGFR* and *FN1*, markers associated with drug resistance, are high in primed cells. Notably, *NGFR*-high cells appear in a different location of the right hand cluster than *AXL* and *EGFR* in Fig. 1D. **E.** We performed ATAC-seq on a population of sorted mixed, EGFR-high, NGFR-high, and EGFR/NGFR-high WM989 cells. Each condition had three replicates. The Scatter plot shows PCA analysis of differential peaks across the samples. The heatmap shows the top 2,000 differential peaks across samples. The order of both rows and columns of the heatmap were determined by Ward's minimum variance clustering. These plots show that although the major difference is between mixed and primed cells, there are consistent differences between EGFR and NGFR-high cells. **F.** We sorted cells based on NT5E staining and then treated them with BRAFi. Images show the fixed NT5E-low and NT5E-high cells stained with DAPI after 3 weeks in the targeted therapy. Each black dot is the nucleus of a drug-resistant cell, and drug-resistant colonies are circled in black. **G.** We sorted the top 0.2% of EGFR/NGFR-high expressing cells and the top 2% of NT5E-high cells, and then treated them with BRAFi. Images show the plates after 3 weeks of treatment with BRAFi, and clusters of cells correspond to resistant colonies. Sorting for NT5E enriches for resistance similarly to EGFR/NGFR as done previously (Shaffer et al. 2017).

### Supplemental Figure 2

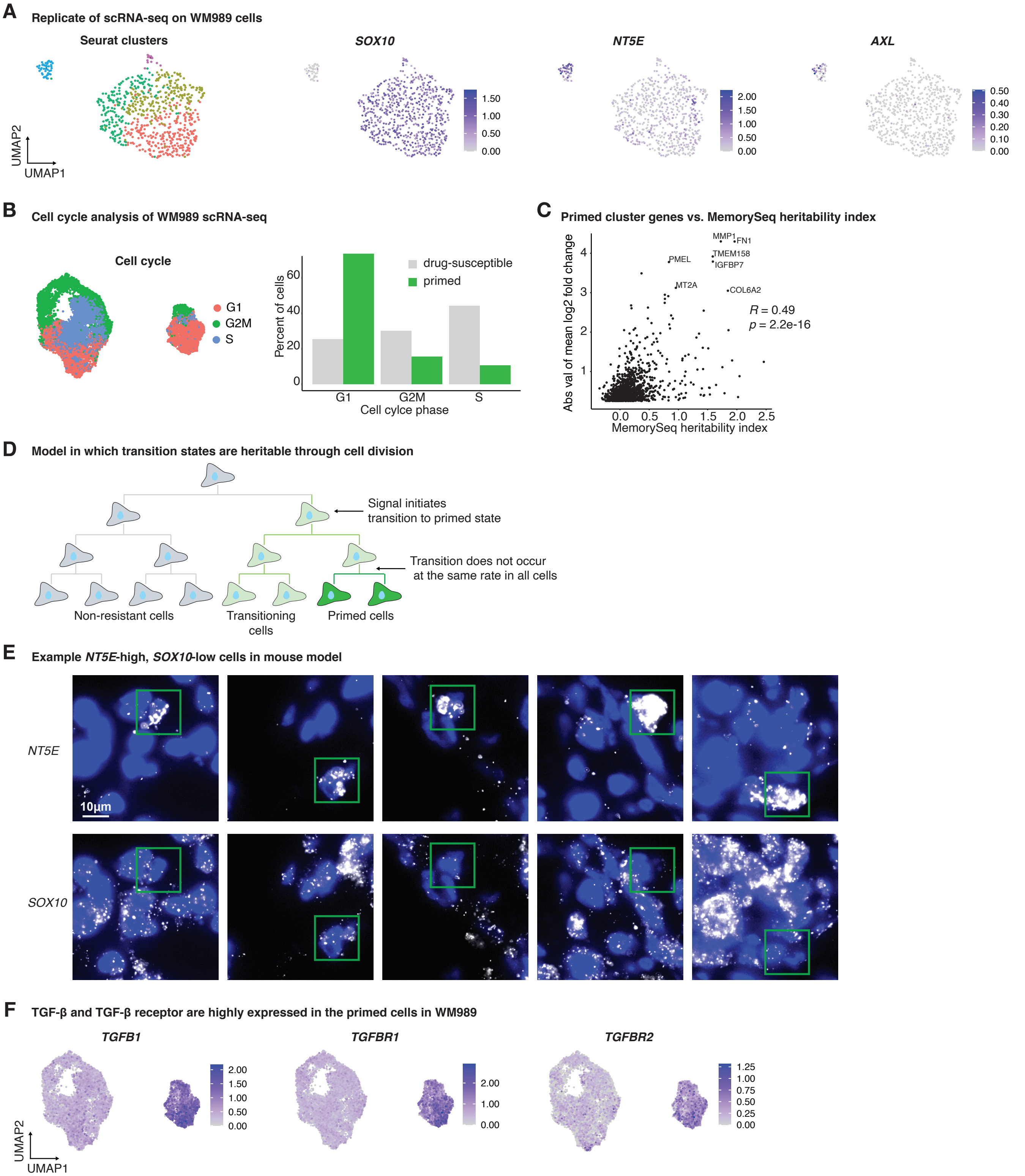

**Supplementary Figure 2: Analysis of primed cell signature. A.** We replicated the scRNA-seq on WM989 cells enriching for primed cells using NT5E as a marker. UMAP plots showing Seurat clustering and gene expression for this replicate. As in previous scRNA-seq of WM989, we can clearly distinguish the drug-susceptible and primed cells and they show differential expression of the same genes. **B.** UMAP plot showing the estimated cell cycle stage of each cell. Bar graph showing the quantification of this data highlighting the comparison in cell cycle between the cells in the drug-susceptible and primed clusters. The high proportion of primed cells in G1 suggests they are cycling slower than drug-susceptible cells. **C.** Scatter plot of the quantification of memory for each gene based on previously published bulk MemorySeq (Shaffer et al. 2020) (x-axis) and the differential expression between primed and drug-susceptible cells in the scRNA-seq (y-axis). From the MemorySeq data, we plotted the residual of the fit between the coefficient of variation and mean expression for each gene (described in the paper at the heritability index in which the higher the residual, the more is memory attributed to that gene), and on the y-axis, we plotted the fold change expression between the primed and drug-susceptible states. These two metrics correlate well showing that scRNA-seq comparison between the primed and drug-susceptible clusters identifies the same memory states. **D.** Schematic of a lineage tree showing how state switching can occur within a lineage from the drug-susceptible to the primed state. Notably, the transition from the drug-susceptible state to the primed state does not occur at the exact same rate in all the branches of the tree. This is important because it allows us to take a measurement at a single point in time at the end of the tree and capture cells undergoing the transition to the primed state. This, along with the assumption that the intermediate state has memory, is what makes it possible to capture the gene expression of transitioning cells in lineages that are crossing. **E.** Additional images of HCR RNA FISH of *NT5E* and *SOX10* in a mouse PDX drug naive tumor shown in Fig. 3A. Images show rare cells with high levels of *NT5E* and low *SOX10*, indicating that primed cells exist in vivo. The green boxes highlight specific cells in the experiment. DAPI is shown in blue and the RNA FISH HCR is displayed in white. Images acquired at 60X magnification. The scale bar represents 10µm. **F.** UMAP plots showing normalized gene expression of *TGFB1*, *TGFBR1* and *TGFBR2*. These three genes are all up-regulated in primed cells suggesting there is likely increased TGF-β signaling in primed cells through autocrine signaling, and potentially some paracrine signaling as well.

### Supplemental Figure 3

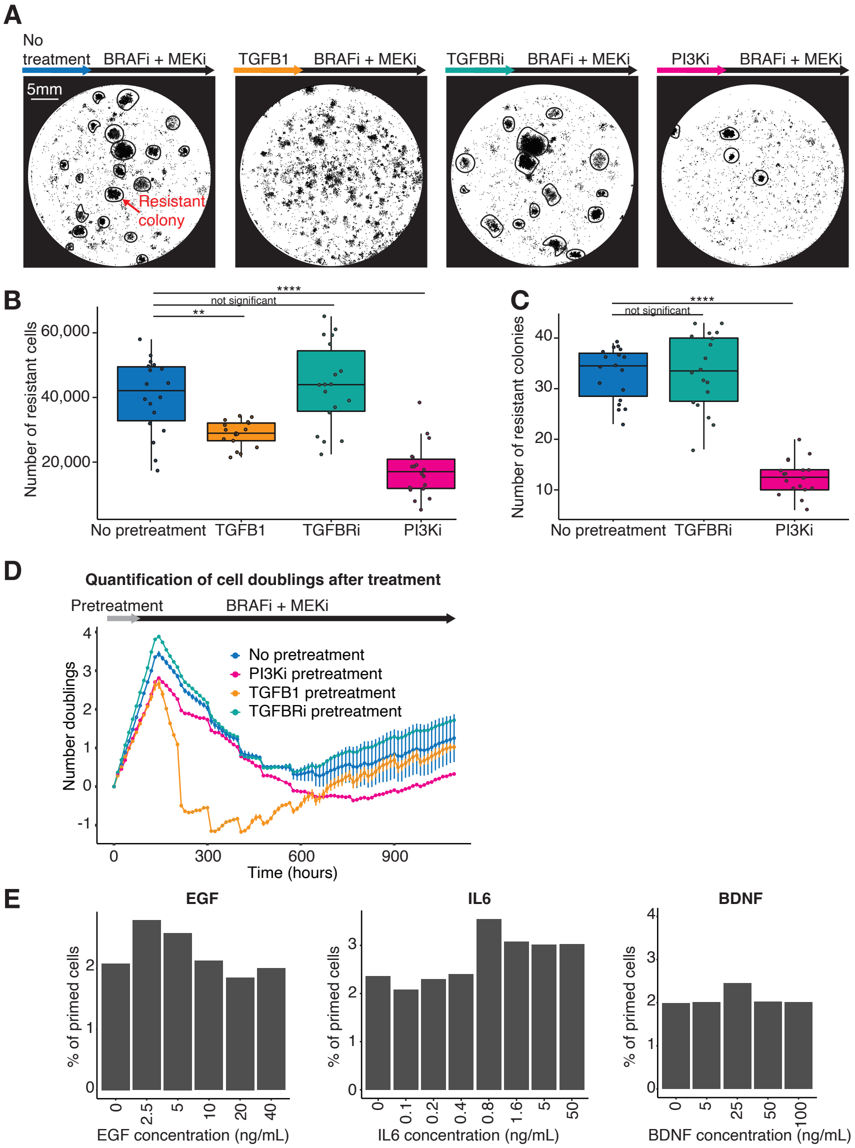

**Supplementary Figure 3: Modulating the number of primed cells affects resistant fate. A.** Example images of fixed cells stained with DAPI for each condition after 5 days of pretreatment and 4 weeks of selection with BRAFi and MEKi. Drug-resistant colonies are circled in black where readily identifiable. Scale bar in the first image represents 5mm and applies for all images in the panel. **B.** Box plot quantifying the number of drug-resistant cells from scans like those shown in panel C across 3 biological replicates, each with 6 technical replicates. P values were calculated using a wilcoxon test (not significant: p > 0.05, *: p <= 0.05, **: p <= 0.01, ***: p <= 0.001, ****: p <= 0.0001) **C.** Box plot quantifying the number of drug colonies (for each condition where distinct colonies were present) from scans like those shown in panel C across 3 biological replicates, each with 6 technical replicates. P values were calculated using a wilcoxon test (not significant: p > 0.05, *: p <= 0.05, **: p <= 0.01, ***: p <= 0.001, ****: p <= 0.0001). **D.** Quantification of live-cell imaging data. Cells were treated with their respective pretreatment for 5 days, and then treated with dabrafenib and trametinib for 40 days. Plots show the doubling of nuclei in each condition measured every 12 hours (relative to the initial number of nuclei). Control and TGFB1 conditions have 2 replicates each, the TGFBRi and the PI3Ki conditions have one replicate each. Error bars are the mean absolute deviation. **E.** Barplots showing the effect of treating with different growth factors and cytokines (EGF, IL6, and BDNF) on the number of primed cells in the population based on NT5E measurements by flow cytometry. These experiments show that the increase in primed cells caused by TGFB1 treatment does not occur with these growth factors or cytokines.

### Supplemental Figure 4

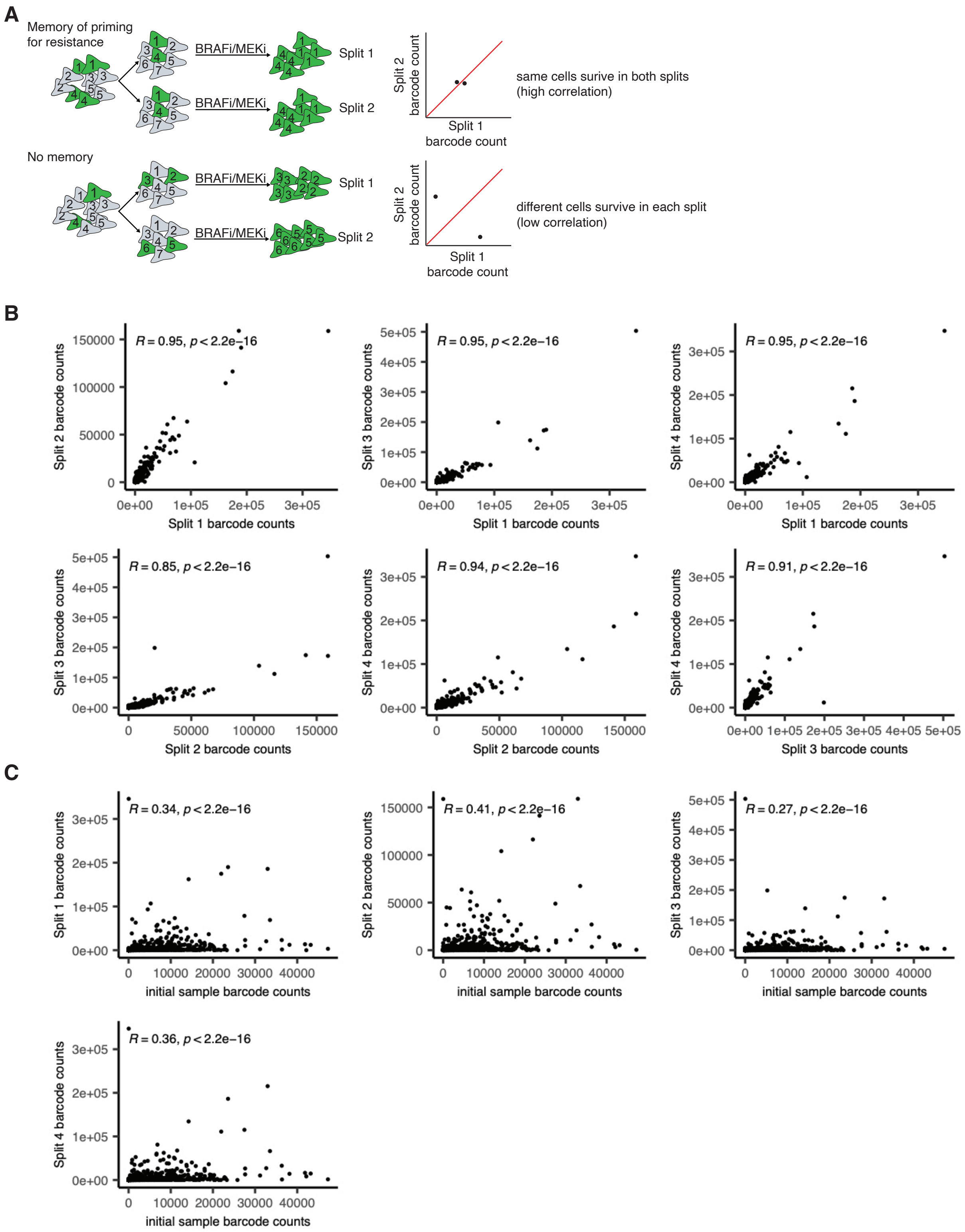

**Supplementary Figure 4: Evidence for cellular memory. A.** Schematic showing the experimental design we used to assess whether there is memory underlying resistance to BRAFi/MEKi over 7-8 doublings (number of cells shown in diagram is less than 7-8 doublings for simplicity). In this assay, we transduced cells with the lentiviral barcoding library (the barcodes are represented by the number in the cell) and then allowed the cells to divide through 7-8 doublings, so there are multiple cells with the same barcode. We then split the cells across multiple plates (two in this example) such that each plate has cells with each barcode. We then treated each plate with BRAFi/MEKi for 4 weeks and recovered the barcodes from gDNA. From the barcode sequences, we can determine whether the same cells survived in each of the plates. The top schematic shows a scenario in which resistance has memory and the bottom schematic shows a scenario in which resistance does not have memory. In the top schematic, the same cells survive in each plate leading to the same barcodes recovered. This outcome leads to highly correlated barcode frequencies in both plates. When memory is not maintained, as shown in the bottom schematic, cells with different barcodes survive in each plate, and therefore barcode abundance in both plates do not correlate. **B.** Results of a memory testing experiment described in A where barcoded cells were split into 4 plates after 7-8 doublings and treated with BRAFi/MEKi for 4 weeks. Scatter plots show pairwise comparison of barcode frequencies in each split. The high correlation in these plots indicated memory was conserved through these divisions. **C.** Data from the same experiment in B, but a sampling of barcodes was taken before cells were treated with BRAFi/MEKi. These scatter plots show the abundance of barcodes before treatment with BRAFi/MEKi on the x-axis, and the barcode abundances in each split on the y-axes. This data shows that barcode frequencies before adding BRAFi/MEKi does not correlate with barcode frequencies after treatment. This demonstrates that BRAFi/MEKi selects for cell state, and is not just selecting for the most abundant barcodes.

### Supplemental Figure 5

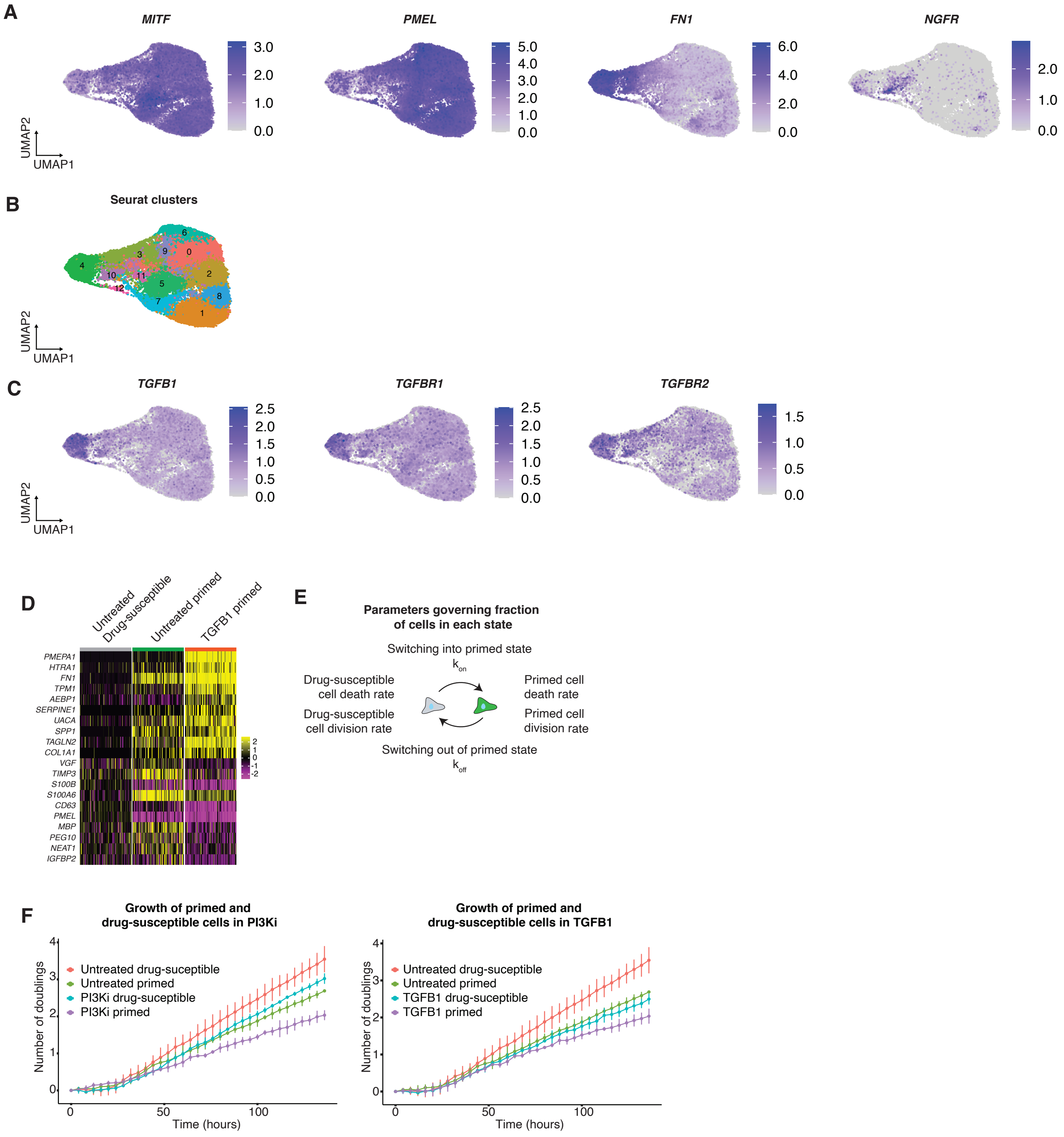

**Supplementary Figure 5: Effects of TGFB1 and PI3Ki on gene expression and proliferation rates. A.** UMAP plots showing normalized gene expression of *MITF*, *PMEL*, *FN1*, and *NGFR* for each cell. The gene expression patterns are very similar to those of scRNA-seq of WM989 cells in Supp. Fig. 1D. *MITF* and *PMEL* are generally anticorrelated with *FN1* and *NGFR*. **B.** Seurat clusters of all cells across all treatments from Fig. 4A. Most primed untreated cells were in cluster 10 and most fully primed cells that were induced by TGFB1 were in cluster 4, thus both clusters 10 and 4 were designated as containing primed cells. **C.** UMAP plots showing normalized gene expression of *TGFB1*, *TGFBR1* and *TGFBR2*. These TGF-β signaling associated genes are again associated with the primed state. Notably, TGFB1 treated primed cells have high levels of expression of *TGFB1*, *TGFBR1* and *TGFBR2*. **D.** Heatmap of gene expression for example genes in a representative sample of cells from the untreated drug-susceptible, untreated primed, and TGFB1 induced primed cells. The heatmap depicts the log10 normalized and scaled gene expression. We find for most primed cell genes that the TGFB1 induced primed cells have higher expression than the untreated primed cells. Notably, there are a few genes upregulated in untreated primed cells that are not upregulated in the TGFB1 treated primed cells including *VGF* and *S100A6*. **E.** To measure the effects of TGFB1 and PI3Ki on proliferation of each cell population, we sorted drug-susceptible and primed cells into separate wells and then treated them with TGFB1 or the PI3Ki (untreated control included). To track proliferation, we performed time-lapse microscopy over 5 days and counted nuclei every 4 hours. Plots show the doubling of nuclei in each condition. Error bars are the mean absolute deviation across 6 replicate wells. We find that both drug-susceptible cells and primed cells slow their proliferation in the presence of TGFB1 or the PI3Ki, but continue to divide.
