## Supplemental Method 1 for "Disrupting cellular memory to overcome drug resistance"

#### 1 Introduction

As discussed previously in this paper, we are interested in studying the dynamics of melanoma cells that switch back and forth between two transient transcriptional states: a "drug-susceptible" state in which they die off when exposed to the drug and a "primed" state in which they survive exposure to the drug. In this section, we discuss the results of an experiment in which populations of these cells were exposed to different treatments, to study their effects on the fraction of cells in the primed state. We will then use stochastic simulations to model different possible mechanisms of these treatments, to see which model scenarios yield results that most closely match the real data.

The experimental procedure was as follows. A population of cells was grown out for approximately 7-8 generations. The population was then split into two groups. One group was exposed to a treatment (either TGFB1, TGFBRi, or PI3Ki), while the other group was left as an untreated control. Both the treated and untreated groups were then grown out for approximately 2 more generations, and then finally sequenced to reveal the transcriptomic state of each cell. Throughout the experiment, the lineages of the cells were tracked. This allows us to compare the fraction of primed cells in the treated group and the untreated group for each lineage. When analyzing these results, we apply a threshold so that only lineages with at least 5 cells total are considered, and this same threshold was applied to the simulation results of our stochastic model.

Table 1: Fraction of Cells in Primed State – Treated vs. Untreated Statistics

| | **TGFB1** | | **TGFBRi** | | **PI3Ki** | |
|---|---|---|---|---|---|---|
| | **Untreated** | **Treated** | **Untreated** | **Treated** | **Untreated** | **Treated** |
| **Mean** | 0.0342 | 0.2900 | 0.0328 | 0.0353 | 0.0328 | 0.0079 |
| **Std. Deviation** | 0.1130 | 0.2615 | 0.1076 | 0.1081 | 0.1092 | 0.0451 |
| **CV** | 3.3032 | 0.9019 | 3.2794 | 3.0611 | 3.3330 | 5.7077 |
| **Fano Factor** | 0.3732 | 0.2359 | 0.3529 | 0.3308 | 0.3639 | 0.2576 |

The results of this experiment are shown in Table 1. Here we consider the fractions of primed cells for each lineage in the treated and untreated groups, and report the mean, standard deviation, coefficient of variation, and Fano factor for these fractions across lineages. Note that the untreated groups in all three experimental cases show very similar statistics – a sign that the control groups are working as expected. Comparing the treated and untreated mean primed fractions in each case allows us to see the effects of the treatments. In the case of TGFB1, the treatment increased the fraction of cells in the primed state. In the case of TGFBRi, the fraction of primed cells in the treated and untreated groups were very similar. In the case of PI3Ki, the treatment decreased the fraction of cells in the primed state.

Based on this experiment, we can clearly see the effects of each treatment. However, the mechanisms that are giving rise to these effects are still unknown. Take, for example, the case of the TGFB1 treatment. We can see that the treatment is increasing the fraction of cells in the primed state, but how exactly is this happening? Is the treatment increasing the rate at which cells switch into the primed state? Or is it decreasing the rate at which cells switch out of the primed state? Or could it be having some effect on the proliferation rates? To study this, we use a stochastic model of the experimental cell population to run computer simulations of different possible scenarios to see which simulated scenarios most closely match the real data from the experiment. We apply this analysis to the TGFB1 and PI3Ki cases. Please note that this should be understood as a hypothesis-generating analysis, not a hypothesis-testing analysis. That is, the conclusions we draw from the model are not definitive confirmations of the true underlying mechanisms of each treatment, but rather clues that could help us design future experiments. The experimental data and Python code used for this analysis are available upon request from the authors.

#### 2 Model Formulation

Let us consider a population of cells that proliferate and switch back and forth between the drug-susceptible state and the primed state. We use the integer-valued stochastic processes $s(t)$ and $p(t)$ to represent the number of cells in the drug-susceptible state and the number of cells in the primed state, respectively. Drug-susceptible cells proliferate with rate $r_1$ and primed cells proliferate with rate $r_2$. Drug-susceptible cells switch to the primed state with rate $k_{on}$ (we also call this "switching on") and primed cells switch back to the drug-susceptible state with rate $k_{off}$ ("switching off"). We refer to the average fraction of primed cells in the untreated state as $f_{exp}$, and this number is slightly different for the different experimental cases.

Table 2 shows the stochastic formulation of this model. Each row is an event that can occur in the cell population, requiring an update of $s(t)$ and $p(t)$, which keep track of the number of cells in each state. Each event occurs with the appropriate rate and propensity. We can use this setup to simulate the model using the stochastic simulation algorithm [1].

Table 2: Stochastic Model

| **Event** | **Var. Updates** | **Rate** | **Propensity** |
|---|---|---|---|
| Drug-susceptible proliferation | $s \to s+1$ | $r_1$ | $r_1 \cdot s$ |
| Primed proliferation | $p \to p+1$ | $r_2$ | $r_2 \cdot p$ |
| Drug-susceptible to primed | $s \to s-1$<br>$p \to p+1$ | $k_{on}$ | $k_{on} \cdot s$ |
| Primed to drug-susceptible | $s \to s+1$<br>$p \to p-1$ | $k_{off}$ | $k_{off} \cdot p$ |

Before fitting this model to data, we make some simplifying assumptions in order to reduce the number of independent parameters we need to fit. First, we set the drug-susceptible state proliferation rate $r_1 = 1$, so that time in our model is normalized to the average generation time of cells in the drug-susceptible state. So, our time unit is average drug-susceptible state generations, rather than hours, days, etc.

Next, we make a simplifying assumption in order to write $k_{on}$ in terms of $k_{off}$. Consider a scenario in which a large population of cells exists in a steady state, with no proliferation, and an average fraction of $\hat{f}$ in the primed state at any given time. Under these theoretical conditions, $\hat{f}$ can be written in terms of the state-switching rates as:

$$\hat{f} = \frac{k_{on}}{k_{on} + k_{off}} \tag{1}$$

The intuition here is that if we know one of the switch-rates and the average primed fraction in the population, we can figure out what the other switch rate must be in order to maintain that equilibrium. We make the simplifying assumption that the average primed fraction across lineages in our experimental data is approximately equal to this underlying theoretical steady state primed fraction: $f_{exp} \approx \hat{f}$. Making this assumption allows us to write the switch-on rate in terms of the switch-off rate and the average primed fraction:

$$k_{on} = \frac{k_{off}}{\frac{1}{f_{exp}} - 1} \tag{2}$$

A consequence of defining $k_{on}$ this way is that different experimental data sets can have different fractions in the primed state, meaning different values for $k_{on}$. That is why the $k_{on}$ values for the TGFB1 and PI3Ki treatment scenarios are slightly different in the following sections.
After normalizing the model time units to average drug-susceptible generation time and making our simplifying assumption to write equation (2), we are left with only two independent parameters to fit to the data: $r_2$ and $k_{off}$.

### 3 Estimating Untreated Parameters

The next step in our analysis is to estimate parameters for the untreated cells. To do this, we use data from another experiment in which cells were grown out for approximately 4 generations, and then sequenced to reveal their transcriptional state. Again, the lineages of the cells were tracked, so we can look at the numbers of drug-susceptible and primed cells from the same original lineage.

We use a modified minimum least-squares method to fit our untreated parameters to this data set. First, we consider a deterministic version of the stochastic model proposed in the previous section:

$$\frac{ds}{dt} = r_1 s - k_{on} s + k_{off} p \tag{3}$$

$$\frac{dp}{dt} = r_2 p + k_{on} s - k_{off} p \tag{4}$$

We then make the same simplifying assumptions described in the previous section, setting $r_1 = 1$ and $k_{on} = \frac{k_{off}}{\frac{1}{f_{exp}} - 1}$. In the case of this experimental data set, the average fraction of primed cells across lineages was 0.125, so $f_{exp} = 0.125$. Plugging in this value gives us $k_{on} = \frac{k_{off}}{7}$. So, the system of equations is:

$$\frac{ds}{dt} = s - \frac{k_{off}}{7} s + k_{off} p \tag{5}$$

$$\frac{dp}{dt} = r_2 p + \frac{k_{off}}{7} s - k_{off} p \tag{6}$$

In our model, we assume that a cell population is grown out from a single original cell, which could be either drug-susceptible or primed. So, this system of ODEs has two possible initial conditions: $(s(0) = 1, p(0) = 0)$ or $(s(0) = 0, p(0) = 1)$. We use the function notation $s_0(r_2, k_{off}), p_0(r_2, k_{off})$ to denote the solutions to the ODEs at $t = 4$, with the initial condition of one drug-susceptible cell and zero primed cells. We use the function notation $s_1(r_2, k_{off}), p_1(r_2, k_{off})$ to denote the solutions to the ODEs at $t = 4$, with the initial condition of zero drug-susceptible cells and one primed cell.

We refer to the experimental drug-susceptible and primed cell counts for each lineage as $\hat{s}$ and $\hat{p}$, respectively. We define a loss function as follows. For each experimental data point, we calculate the squared errors for the drug-susceptible and primed counts, and add them together, assuming the initial condition $(s(0) = 1, p(0) = 0)$. We then do the same calculation, assuming the initial condition $(s(0) = 0, p(0) = 1)$. After calculating both of these measures, we take the minimum of the two and add that to our total loss. We do this for each experimental data point. The mathematical formulation of this loss function is:

$$l(r_2, k_{off}) = \sum_{i=1}^{N} \min\left\{(s_0(r_2, k_{off}) - \hat{s}_i)^2 + (p_0(r_2, k_{off}) - \hat{p}_i)^2, (s_1(r_2, k_{off}) - \hat{s}_i)^2 + (p_1(r_2, k_{off}) - \hat{p}_i)^2\right\} \tag{7}$$

Then, we simply apply a numerical optimization in Python to find the values of $r_2$ and $koff$ that minimize the loss function.

$$\min_{r_2, k_{off}} \quad l(r_2, k_{off}) \tag{8}$$

Table 3 shows estimates for these parameter values that we have obtained through this method. We use a bootstrapping technique, running the analysis 40 times on random samples of the original data set, in order to generate a distribution of results. We report the 2.5 percentile value, median, and 97.5 percentile value of the results, so that the range between the lower bound and upper bound includes 95% of observed parameter fit values. The data and Python code used for the fitting are available upon request. Please note that these parameter values should be taken as rough estimates, not as exact, certain values.

Table 3: Estimated Parameter Values

| **Event description** | **Parameter** | **Fit Value** | | |
|---|---|---|---|---|
| | | **95% Lower Bound** | **Median** | **95% Upper Bound** |
| Primed cell proliferation | $r_2$ | 0.3714 | 0.4786 | 0.5605 |
| Primed to drug-susceptible switch | $k_{off}$ | 0.1247 | 0.1666 | 0.2158 |

### 4 Experiment Simulation

As discussed previously, we have data from an experiment in which cells were grown out for a period of time and then divided into two groups, with some given a treatment (TGFB1, TGFBRi, or PI3Ki), while others were left untreated. Since lineages were tracked throughout this experiment, we can compare the fraction of cells in the primed state for both treated and untreated cells within the same lineage. For example, Figure 1 is a plot of the fraction of treated cells in the primed state against the fraction of untreated cells in the primed state, for the TGFB1 experiment. Each point represents these fractions for a single lineage.

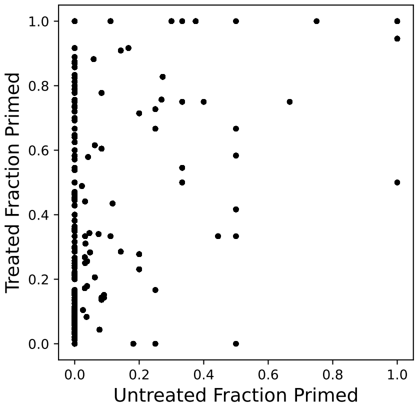


Figure 1: Fraction of primed treated cells vs. fraction of primed untreated cells per lineage, for the TGFB1 treatment case.

Our goal is to use the stochastic model described previously to generate simulated datasets like this for different possible mechanisms of the treatments, and then see how closely the simulated results for different possible scenarios fit the real data.

We simulate the experiment as follows. We begin with a single cell, which is randomly chosen to be in the primed state with probability $f_{exp}$ or to be in the drug-susceptible state with probability $1 - f_{exp}$. The cell is grown out into a colony from $t = 0$ until $t = 7$ (approximately 7 drug-susceptible state generations in our model). During this time, the cells proliferate and switch back and forth between the drug-susceptible and primed states. For this part of the simulation, we use the fitted untreated parameters listed in Table 3. At the end of this portion of the simulation, we are left with a simulated population of cells, which may be drug-susceptible, primed, or a mix. At this point, we divide the simulated cell population in half, making the simplifying assumption that they are divided exactly in half, which both divisions having exactly the same number of primed cells and exactly the same number of drug-susceptible cells. When an odd number interferes with this, we round the quotient to make this division feasible.

From this point, we use the two groups of simulated cells to simulate the second part of the experiment, in which one group is exposed to the treatment while the other is left untreated. In terms of our simulation, we continue to use the same parameters for the untreated group. For the treated group, we continue the simulation, but change one of the parameters to model a possible mechanism of the treatment. The process of selecting these new, treated parameter values is described in the next section.

### 5 Estimating Treated Parameters

As explained in the previous section, we model the effect of the treatment by changing one of the parameters, while keeping every other parameter the same as in the untreated case. For different scenarios, we change different parameters. Our ultimate goal is to compare the log-likelihood scores of different scenarios to see which ones can better explain the data. However, before we can compare across different scenarios, we must first select the treated parameter values for each case. We use a truncated, rough approximation of the maximum log-likelihood method to choose these parameter values.

Let us quickly review the concept of log-likelihood. Consider a set of experimental observations $X_1, X_2, ..., X_N$. We can attempt to model the system that yielded these observations, using a set of input parameters $\theta$ that yields a probability distribution $f_X(X, \theta)$. For a given parameter set $\theta$, we can calculate the log-likelihood of observing the data points:

$$\sum_{i=1}^{N} log(f_X(X_i, \theta)) \tag{9}$$

Here *log* is the natural logarithm. Then, to fit our model to the data, our task is to search through the space of possible parameter values to find the values of $\theta$ that maximize this measure of log-likelihood.

We use this approach to estimate the new treated parameter value for each treatment scenario. Since we do not have an exact analytical solution for the joint probability distribution of the prime fractions of treated and untreated cells in our model, we estimate this distribution empirically, by running many simulations and binning the results. This involves running the simulated experiment many times for each new parameter value being tested, so the process is quite computationally expensive and has a considerable runtime. Due to this computational constraint, we estimate the empirical probability distributions for each parameter set using only 1000 runs of the experiment simulation, a smaller than ideal number. This also leads to another computational problem – when empirically estimating the distributions, it is possible to have a bin with 0 observations in the model and at least 1 observation in the experimental data. This causes a calculation error, since we cannot take the natural logarithm of 0. So, when binning the results of simulations, we assume that each bin starts at 1, not 0, and count observations from there. To ensure that this computational shortcut is not skewing the results, we will drop it in the following section when we validate the log-likelihood scores for each scenario by using 1000000 runs of the model to generate the distributions.

Using this log-likelihood parameter estimation method, we test 400 potential new parameter values for each scenario, chosen from a uniform distribution, from the previous untreated parameter value to some upper or lower bound. We save the value that yields the highest log-likelihood score, and use that as our new treated parameter value for that scenario. The new parameter values for each scenario are shown in the following two tables.

Table 4: TGFB1 - Treated Parameter Estimation

| **Scenario** | **Changed Parameter** | **Untreated Value** | **Treated Value** |
|---|---|---|---|
| Drug-Susceptible Proliferation Decrease | $r_1$ | 1 | 0.0166 |
| Primed Proliferation Increase | $r_2$ | 0.4786 | 2.9059 |
| Switch-On Rate Increase | $k_{on}$ | $\frac{k_{off}}{\frac{1}{f_{exp}}-1} = 0.0059$ | 0.0855 |
| Dependent Switch-On Rate | $k_{on}$ | $\frac{k_{off}}{\frac{1}{f_{exp}}-1} = 0.0059$ | $0.0013 \cdot p$ |
| Switch-Off Rate Decrease | $k_{off}$ | 0.1666 | 0.0651 |

Table 5: PI3Ki - Treated Parameter Estimation

| **Scenario** | **Changed Parameter** | **Untreated Value** | **Treated Value** |
|---|---|---|---|
| Drug-Susceptible Proliferation Increase | $r_1$ | 1 | 2.2954 |
| Primed Proliferation Decrease | $r_2$ | 0.4786 | 0.0473 |
| Switch-On Rate Decrease | $k_{on}$ | $\frac{k_{off}}{\frac{1}{f_{exp}}-1} = 0.0056$ | 0.0012 |
| Switch-Off Rate Increase | $k_{off}$ | 0.1666 | 3.3485 |

### 6 TGFB1 Treatment

We have observed that when the cell populations are treated with TGFB1, that yields an increased fraction of cells in the primed state compared to an untreated control. However, the underlying mechanism behind this is not known. We consider five possible scenarios that could yield an increase in the fraction of cells in the primed state:

1. A decrease in the drug-susceptible proliferation rate $r_1$
2. An increase in the primed proliferation rate $r_2$
3. An increase in the switch-on rate $k_{on}$
4. A decrease in the switch-off rate $k_{off}$
5. A scenario in which the switch-on rate $k_{on}$ depends on the number of primed cells $p$

For each of these possible scenarios, we ran a simulation of the experiment 1000000 times, and binned the results in order to generate an empirical probability distribution. We used this probability distribution of simulated results to calculate a log-likelihood score for each scenario. Note that in terms of log-likelihood, a higher number means a better fit, and a more negative number means a worse fit. The results are shown in Table 6, listed in order of best-fit to worst-fit.

Figure 2 shows a visual representation of these results. For each plot of treated prime cell fractions to untreated prime cell fractions, the back dots show real data points from the experimental dataset. The blue shading shows the distribution of simulation results for each scenario. The shading is not exactly proportional to the distribution frequency, and has been adjusted to saturate at a level below the maximum to

Table 6: TGFB1 - Simulation Results

| **Scenario** | **Changed Parameter** | **Log-Likelihood Score** |
|---|---|---|
| Primed Proliferation Increase | $r_2$ | $-50563.78$ |
| Dependent Switch-On Rate | $k_{on}$ | $-54531.503$ |
| Drug-Susceptible Proliferation Decrease | $r_1$ | $-84862.688$ |
| Switch-On Rate Increase | $k_{on}$ | $-88256.291$ |
| Switch-Off Rate Decrease | $k_{off}$ | $-112885.818$ |

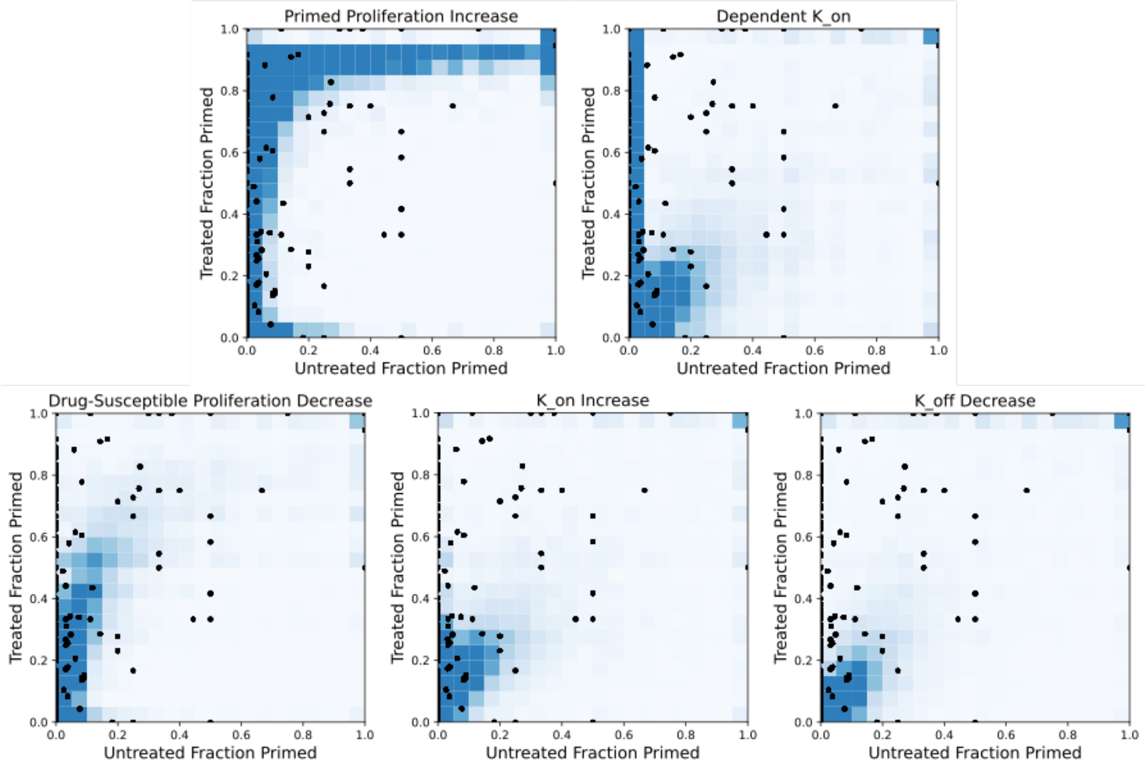


Figure 2: Fraction of primed treated cells vs. fraction of primed untreated cells per lineage, for the TGFB1 treatment case. Real data points, shown in black dots, are plotted over the distribution of simulation results, shown with blue shading.

highlight simulation outliers (in order to better to compare to the real data point outliers, which are visible).

Please note that many of the real experimental data points in these plots lie along the vertical axis (this may be difficult to see). We believe that the difference in the log-likelihood scores between scenarios is largely the result of their relative ability or inability to account for these data points. The Primed Proliferation Increase and Dependent Switch-On Rate scenarios are able to account for these data points along the vertical axis (as can be seen by their dark blue shaded bars along the vertical axis), while the Drug-Susceptible Proliferation Decrease, Switch-On Rate Increase, and Switch-Off Rate Decrease scenarios do not account for them as well.

### 7 PI3Ki Treatment

In the PI3Ki treatment data, we observed a decrease in the fraction of cells in the primed state in the treated sample compared to the untreated control. The underlying mechanism behind this is not known. In this section, we consider four possible scenarios:

1. An increase in the drug-susceptible proliferation rate $r_1$
2. A decrease in the primed proliferation rate $r_2$
3. A decrease in the switch-on rate $k_{on}$
4. An increase in the switch-off rate $k_{off}$

We applied the same analytical workflow here as in the TGFB1 analysis, running a simulation of the experiment 1000000 times and binning the results in order to generate an empirical probability distribution. Again, we used this simulated probability distribution to calculate a log-likelihood score for each scenario. These results are shown in Table 7, in order of best-fit to worst-fit.

Table 7: PI3Ki - Simulation Results

| **Scenario** | **Changed Parameter** | **Log-Likelihood Score** |
|---|---|---|
| Drug-Susceptible Proliferation Increase | $r_1$ | $-10544.337$ |
| Switch-Off Rate Increase | $k_{off}$ | $-10991.598$ |
| Primed Proliferation Decrease | $r_2$ | $-11017.362$ |
| Switch-On Rate Decrease | $k_{on}$ | $-11774.535$ |

Figure 3 shows a visual representation of these results, plotting the treated prime cell fractions against the untreated prime cell fractions. The black dots show real data points from the experimental dataset, and the blue shading shows the distribution of simulation results for each scenario. As with Figure 2, the shading in Figure 3 is not exactly proportional to the distribution frequency, and has been adjusted to saturate at a level below the maximum to highlight simulation outliers.

Although it is slightly difficult to see visually, we note that many of the real experimental data points lie along the horizontal axis, and the relative goodness or badness of fit for the scenarios seems to be determined by their ability or inability to account for these data points.

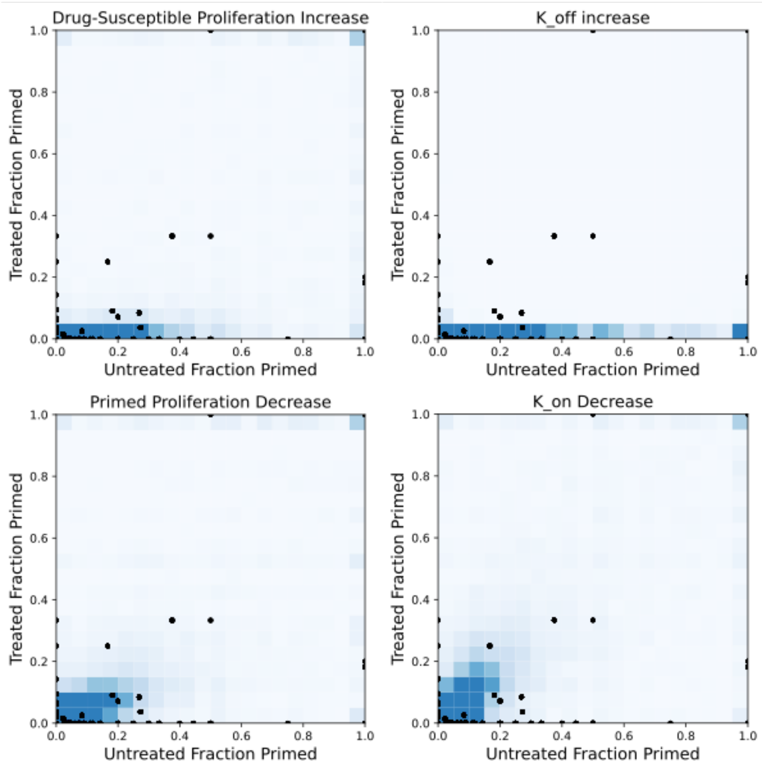


Figure 3: Fraction of primed treated cells vs. fraction of primed untreated cells per lineage, for the PI3Ki treatment case. Real data, shown in black dots, are plotted over the distribution of simulation results, shown with blue shading.
