## Supplemental Table 1 for "Disrupting cellular memory to overcome drug resistance"

SupplementaryTable1

| | p_val | avg_log2FC | pct.1 | pct.2 | p_val_adj |
|---|---|---|---|---|---|
| **ERRFI1** | 0 | 1.31179658468425 | 0.92 | 0.209 | 0 |
| **PIK3CD-AS2** | 0 | -0.312019703919717 | 0.55 | 0.836 | 0 |
| **SRM** | 0 | -0.792641970816508 | 0.988 | 0.999 | 0 |
| **RCC2** | 0 | -0.560559101252851 | 0.621 | 0.922 | 0 |
| **CAMK2N1** | 0 | 2.67623442401557 | 0.988 | 0.487 | 0 |
| **CDA** | 0 | 0.610752263442262 | 0.626 | 0.022 | 0 |
| **PINK1** | 0 | 0.362381803729335 | 0.846 | 0.542 | 0 |
| **RPL11** | 0 | -0.595207271099607 | 1 | 1 | 0 |
| **TRIM63** | 0 | -0.612156384342904 | 0.214 | 0.793 | 0 |
| **ZNF593** | 0 | -0.422432426192072 | 0.962 | 0.987 | 0 |
| **SH3BGRL3** | 0 | 1.39670304611431 | 1 | 0.998 | 0 |
| **IFI6** | 0 | 1.21979942451517 | 0.943 | 0.914 | 0 |
| **EPB41** | 0 | -0.386454537942468 | 0.693 | 0.914 | 0 |
| **SDC3** | 0 | -0.33178961955499 | 0.55 | 0.828 | 0 |
| **ZCCHC17** | 0 | -0.448007160743117 | 0.923 | 0.977 | 0 |
| **SERINC2** | 0 | 0.462131366298573 | 0.669 | 0.373 | 0 |
| **SPOCD1** | 0 | 0.446747090519719 | 0.747 | 0.075 | 0 |
| **FAM167B** | 0 | -0.377176882923078 | 0.548 | 0.842 | 0 |
| **MARCKSL1** | 0 | -0.704366264579438 | 0.941 | 0.989 | 0 |
| **PHC2** | 0 | 0.533499473218683 | 0.973 | 0.905 | 0 |
| **EVA1B** | 0 | 0.461816917542747 | 0.963 | 0.871 | 0 |
| **MRPS15** | 0 | -0.399157973482105 | 0.989 | 0.996 | 0 |
| **CTPS1** | 0 | -0.382416884398501 | 0.698 | 0.919 | 0 |
| **YBX1** | 0 | -0.827181279987811 | 1 | 1 | 0 |
| **PRDX1** | 0 | -1.97187436926202 | 1 | 1 | 0 |
| **AKR1A1** | 0 | -1.47018952418274 | 0.963 | 0.988 | 0 |
| **PIK3R3** | 0 | -1.03905976771153 | 0.428 | 0.961 | 0 |
| **UQCRH** | 0 | -0.354348464024635 | 1 | 1 | 0 |
| **CDKN2C** | 0 | 0.336612521955521 | 0.763 | 0.451 | 0 |

| | | | | | |
|---|---|---|---|---|---|
| **MRPL37** | 0 | -0.434445302959112 | 0.934 | 0.98 | 0 |
| **JUN** | 0 | 0.834083872302609 | 0.794 | 0.096 | 0 |
| **PGM1** | 0 | -0.452311194804683 | 0.883 | 0.981 | 0 |
| **JAK1** | 0 | 0.352442981977661 | 0.981 | 0.94 | 0 |
| **SERBP1** | 0 | -0.849753865135865 | 0.998 | 0.999 | 0 |
| **GADD45A** | 0 | 0.421523948665455 | 0.867 | 0.568 | 0 |
| **LHX8** | 0 | 0.35368163458518 | 0.708 | 0.249 | 0 |
| **AK5** | 0 | 0.373718474442621 | 0.725 | 0.206 | 0 |
| **IFI44** | 0 | 0.300319470719465 | 0.716 | 0.374 | 0 |
| **CTBS** | 0 | 0.340097174072423 | 0.917 | 0.758 | 0 |
| **ODF2L** | 0 | 0.37691374950761 | 0.951 | 0.82 | 0 |
| **GBP2** | 0 | 0.415028362941473 | 0.683 | 0.093 | 0 |
| **RPL5** | 0 | -0.640722123742707 | 1 | 1 | 0 |
| **CSF1** | 0 | 0.587224677447721 | 0.77 | 0.076 | 0 |
| **RHOC** | 0 | 0.450367572061208 | 0.995 | 0.99 | 0 |
| **SLC16A1** | 0 | -0.411698533358653 | 0.662 | 0.902 | 0 |
| **OLFML3** | 0 | 0.634984132585093 | 0.729 | 0.038 | 0 |
| **ATP1A1** | 0 | -1.08920144665014 | 0.966 | 0.998 | 0 |
| **IGSF3** | 0 | -0.495375605493414 | 0.298 | 0.848 | 0 |
| **TTF2** | 0 | -0.378307483651269 | 0.655 | 0.912 | 0 |
| **ITGA10** | 0 | 0.494301310543169 | 0.731 | 0.376 | 0 |
| **AC245297.3** | 0 | 0.282133913057468 | 0.835 | 0.58 | 0 |
| **PSMB4** | 0 | -0.472884888745101 | 0.971 | 0.993 | 0 |
| **S100A10** | 0 | 0.79470592011939 | 1 | 1 | 0 |
| **S100A11** | 0 | 0.995624239802114 | 1 | 0.998 | 0 |
| **S100A6** | 0 | 2.84270582033399 | 1 | 1 | 0 |
| **S100A4** | 0 | 1.10849494161901 | 0.896 | 0.39 | 0 |
| **S100A3** | 0 | 0.31537419573954 | 0.667 | 0.13 | 0 |
| **S100A2** | 0 | 0.343949785850995 | 0.678 | 0.275 | 0 |
| **S100A16** | 0 | 0.781402060888287 | 0.931 | 0.666 | 0 |
| **S100A13** | 0 | 2.04926772943321 | 1 | 0.997 | 0 |
| **S100A1** | 0 | -0.870003210231624 | 0.878 | 0.976 | 0 |

| | | | | | |
|---|---|---|---|---|---|
| **SLC27A3** | 0 | -0.607342989197405 | 0.414 | 0.778 | 0 |
| **RAB13** | 0 | 0.682280170381855 | 1 | 0.997 | 0 |
| **CCT3** | 0 | -0.485673137725231 | 0.998 | 0.999 | 0 |
| **BCAN** | 0 | -0.474393464285923 | 0.092 | 0.757 | 0 |
| **IFI16** | 0 | -0.692838590967787 | 0.99 | 0.999 | 0 |
| **TAGLN2** | 0 | 1.15505983287661 | 0.998 | 0.984 | 0 |
| **IGSF8** | 0 | -0.646144680082008 | 0.629 | 0.926 | 0 |
| **OLFML2B** | 0 | 0.360528674001761 | 0.647 | 0.019 | 0 |
| **DDR2** | 0 | 0.385469212810246 | 0.963 | 0.852 | 0 |
| **UCK2** | 0 | -0.32680653620371 | 0.663 | 0.883 | 0 |
| **MPC2** | 0 | 1.01379404491346 | 0.998 | 0.989 | 0 |
| **DCAF6** | 0 | 0.372255655102425 | 0.92 | 0.772 | 0 |
| **ATP1B1** | 0 | 0.253053450161324 | 0.508 | 0.133 | 0 |
| **PRRX1** | 0 | 0.921353915496248 | 0.98 | 0.741 | 0 |
| **C1orf21** | 0 | -0.342998212104791 | 0.77 | 0.93 | 0 |
| **IVNS1ABP** | 0 | -0.588221182508472 | 0.771 | 0.957 | 0 |
| **HMCN1** | 0 | -0.365871442505354 | 0.207 | 0.764 | 0 |
| **RGS2** | 0 | 1.07275578124353 | 0.83 | 0.166 | 0 |
| **IGFN1** | 0 | 1.3277367954186 | 0.781 | 0.037 | 0 |
| **SNRPE** | 0 | -0.49776292063698 | 0.992 | 0.999 | 0 |
| **DSTYK** | 0 | -0.890875301962491 | 0.708 | 0.976 | 0 |
| **IL24** | 0 | 0.408989230390595 | 0.25 | 0.016 | 0 |
| **FCMR** | 0 | 0.420074059174692 | 0.854 | 0.649 | 0 |
| **LAMB3** | 0 | 0.379225890802956 | 0.699 | 0.142 | 0 |
| **G0S2** | 0 | 1.55919727049204 | 0.479 | 0.032 | 0 |
| **PTPN14** | 0 | 0.59794856818358 | 0.93 | 0.676 | 0 |
| **CAPN2** | 0 | 0.649463875571225 | 0.971 | 0.946 | 0 |
| **CNIH3** | 0 | 0.586520876407466 | 0.888 | 0.321 | 0 |
| **PARP1** | 0 | -0.875064101308022 | 0.979 | 0.998 | 0 |
| **IRF2BP2** | 0 | 0.440171530131315 | 0.963 | 0.864 | 0 |
| **SDCCAG8** | 0 | 0.519581786953917 | 0.932 | 0.736 | 0 |
| **RPS7** | 0 | -0.48552689416855 | 1 | 1 | 0 |

| | | | | | |
|---|---|---|---|---|---|
| **HPCAL1** | 0 | 2.3462766157576 | 1 | 0.953 | 0 |
| **ODC1** | 0 | -0.508891310056024 | 0.621 | 0.93 | 0 |
| **LAPTM4A** | 0 | 0.960825760343623 | 0.991 | 0.956 | 0 |
| **ATRAID** | 0 | 0.390946728911746 | 0.998 | 0.993 | 0 |
| **FOSL2** | 0 | 0.275229785951258 | 0.699 | 0.257 | 0 |
| **YPEL5** | 0 | 0.373205358652536 | 0.827 | 0.433 | 0 |
| **LTBP1** | 0 | 0.733738560489844 | 0.813 | 0.045 | 0 |
| **CRIM1** | 0 | 0.350576921827701 | 0.63 | 0.067 | 0 |
| **FEZ2** | 0 | 0.331906449728804 | 0.939 | 0.812 | 0 |
| **PRKD3** | 0 | -0.421297690321795 | 0.737 | 0.901 | 0 |
| **QPCT** | 0 | -1.01138109918708 | 0.82 | 0.972 | 0 |
| **CDC42EP3** | 0 | 1.16153730346661 | 0.968 | 0.778 | 0 |
| **EPAS1** | 0 | 0.339588425931606 | 0.718 | 0.246 | 0 |
| **RHOQ** | 0 | -0.636447789689757 | 0.93 | 0.985 | 0 |
| **SOCS5** | 0 | 0.578134723692855 | 0.885 | 0.456 | 0 |
| **CALM2** | 0 | 0.802307377375569 | 0.999 | 0.996 | 0 |
| **RTN4** | 0 | 0.924458566452386 | 1 | 0.997 | 0 |
| **CFAP36** | 0 | 0.375731614687466 | 0.976 | 0.921 | 0 |
| **FAM161A** | 0 | -0.327942164104055 | 0.627 | 0.861 | 0 |
| **ANTXR1** | 0 | 0.364580417038249 | 0.85 | 0.567 | 0 |
| **FAM136A** | 0 | -0.414051608673641 | 0.869 | 0.962 | 0 |
| **CCT7** | 0 | -0.563903498914445 | 0.972 | 0.991 | 0 |
| **MTHFD2** | 0 | -0.431853249131284 | 0.804 | 0.961 | 0 |
| **WDR54** | 0 | 0.408251721861416 | 0.946 | 0.835 | 0 |
| **SUCLG1** | 0 | -0.441574893381151 | 0.962 | 0.988 | 0 |
| **TMSB10** | 0 | 1.68565448382131 | 1 | 1 | 0 |
| **TGOLN2** | 0 | 0.421809354897923 | 0.99 | 0.961 | 0 |
| **RPIA** | 0 | -0.279732074204586 | 0.491 | 0.792 | 0 |
| **TMEM131** | 0 | 0.403342640863203 | 0.918 | 0.763 | 0 |
| **RPL31** | 0 | -0.668451847535434 | 0.996 | 0.999 | 0 |
| **MAP4K4** | 0 | 2.05894921337861 | 0.998 | 0.98 | 0 |
| **LINC01918** | 0 | -0.669733580584505 | 0.072 | 0.741 | 0 |

| | | | | | |
|---|---|---|---|---|---|
| **FHL2** | 0 | 0.597785019837268 | 0.941 | 0.842 | 0 |
| **NCK2** | 0 | -0.414319538617818 | 0.613 | 0.902 | 0 |
| **TMEM87B** | 0 | 0.493310366526092 | 0.877 | 0.464 | 0 |
| **SLC20A1** | 0 | 1.84267147022538 | 0.997 | 0.919 | 0 |
| **IL1B** | 0 | 0.798695951138105 | 0.345 | 0.008 | 0 |
| **STEAP3** | 0 | 0.492824784352982 | 0.805 | 0.212 | 0 |
| **BIN1** | 0 | 0.491208987908008 | 0.823 | 0.184 | 0 |
| **IMP4** | 0 | -0.580118450429534 | 0.945 | 0.99 | 0 |
| **MGAT5** | 0 | -0.280565213248437 | 0.568 | 0.843 | 0 |
| **LYPD6B** | 0 | 0.28758510635412 | 0.408 | 0.004 | 0 |
| **LYPD6** | 0 | 0.878613874664035 | 0.838 | 0.354 | 0 |
| **RND3** | 0 | 0.85136371674136 | 0.969 | 0.815 | 0 |
| **FMNL2** | 0 | 0.404873825887523 | 0.939 | 0.826 | 0 |
| **CYTIP** | 0 | 0.310142290468165 | 0.416 | 0.004 | 0 |
| **TANK** | 0 | 0.524116070525221 | 0.943 | 0.712 | 0 |
| **FAP** | 0 | 0.377432490535094 | 0.67 | 0.011 | 0 |
| **COBLL1** | 0 | -0.29725752724618 | 0.197 | 0.678 | 0 |
| **MAP3K20** | 0 | 0.418688396665618 | 0.944 | 0.846 | 0 |
| **ATP5MC3** | 0 | -0.539270614825461 | 0.999 | 1 | 0 |
| **LINC01116** | 0 | 0.402095502760245 | 0.931 | 0.819 | 0 |
| **UBE2E3** | 0 | 0.450361750169609 | 0.975 | 0.93 | 0 |
| **TFPI** | 0 | 0.319834328795061 | 0.648 | 0.12 | 0 |
| **C2orf88** | 0 | -0.358288415264979 | 0.05 | 0.681 | 0 |
| **HIBCH** | 0 | -0.549032137809406 | 0.71 | 0.918 | 0 |
| **STK17B** | 0 | 0.815789080347348 | 0.963 | 0.731 | 0 |
| **HSPD1** | 0 | -0.750433532456257 | 0.999 | 1 | 0 |
| **CFLAR** | 0 | 0.641115103783449 | 0.906 | 0.446 | 0 |
| **FZD7** | 0 | 0.328865172788201 | 0.723 | 0.353 | 0 |
| **NRP2** | 0 | 0.56232773318067 | 0.962 | 0.885 | 0 |
| **ZDBF2** | 0 | 0.316006537869024 | 0.793 | 0.507 | 0 |
| **FN1** | 0 | 4.29856337303469 | 1 | 0.759 | 0 |
| **TNS1** | 0 | -0.414550838664154 | 0.353 | 0.82 | 0 |

| | | | | | |
|---|---|---|---|---|---|
| **SCG2** | 0 | 1.39453651886537 | 0.803 | 0.15 | 0 |
| **WDFY1** | 0 | -0.516528256608561 | 0.754 | 0.947 | 0 |
| **IRS1** | 0 | 0.539993948291178 | 0.798 | 0.177 | 0 |
| **PID1** | 0 | 0.261987235788723 | 0.612 | 0.16 | 0 |
| **SP100** | 0 | 0.517956479102727 | 0.957 | 0.866 | 0 |
| **ITM2C** | 0 | 0.880262533145147 | 1 | 0.99 | 0 |
| **NCL** | 0 | -0.736828364616334 | 0.999 | 1 | 0 |
| **ARL4C** | 0 | 2.2296707246749 | 0.992 | 0.317 | 0 |
| **MLPH** | 0 | 0.62073003526789 | 0.98 | 0.885 | 0 |
| **RAMP1** | 0 | -0.796318231207661 | 0.516 | 0.916 | 0 |
| **HES6** | 0 | -1.00500291460032 | 0.427 | 0.95 | 0 |
| **SNED1** | 0 | 0.303407877895118 | 0.438 | 0.007 | 0 |
| **HDLBP** | 0 | 0.46462449379888 | 0.999 | 0.999 | 0 |
| **FARP2** | 0 | -0.361097375094147 | 0.492 | 0.818 | 0 |
| **CHL1** | 0 | -0.282764461868865 | 0.037 | 0.588 | 0 |
| **HRH1** | 0 | 0.255291727957135 | 0.627 | 0.112 | 0 |
| **MRPS25** | 0 | -0.500565474472058 | 0.855 | 0.964 | 0 |
| **SH3BP5** | 0 | -0.336608804819024 | 0.354 | 0.742 | 0 |
| **RPL15** | 0 | -0.397239340021613 | 1 | 1 | 0 |
| **TGFBR2** | 0 | 0.295477128546993 | 0.722 | 0.204 | 0 |
| **RPSA** | 0 | -0.472657245690298 | 0.999 | 1 | 0 |
| **TMEM158** | 0 | 3.91830251554707 | 0.997 | 0.31 | 0 |
| **SMARCC1** | 0 | -0.375848359823722 | 0.908 | 0.977 | 0 |
| **UCN2** | 0 | -0.678169596094036 | 0.324 | 0.703 | 0 |
| **RPL29** | 0 | -0.582726423163565 | 1 | 1 | 0 |
| **WNT5A** | 0 | 0.571453229042358 | 0.667 | 0.023 | 0 |
| **THOC7** | 0 | -0.515678382062924 | 0.999 | 1 | 0 |
| **PRICKLE2** | 0 | 0.258710209890759 | 0.723 | 0.378 | 0 |
| **MAGI1** | 0 | 0.648912681036722 | 0.966 | 0.823 | 0 |
| **TMF1** | 0 | 0.514420506039062 | 0.999 | 0.995 | 0 |
| **ARL6IP5** | 0 | 0.909243758020587 | 1 | 0.998 | 0 |
| **FRMD4B** | 0 | -1.50261328676272 | 0.601 | 0.957 | 0 |

| | | | | | |
|---|---|---|---|---|---|
| **MITF** | 0 | -1.47138698949666 | 0.785 | 0.995 | 0 |
| **FOXP1** | 0 | 0.537260664717074 | 0.991 | 0.902 | 0 |
| **CHMP2B** | 0 | -0.393702755145774 | 0.912 | 0.972 | 0 |
| **ST3GAL6** | 0 | -0.534893501390295 | 0.594 | 0.919 | 0 |
| **DCBLD2** | 0 | 0.974893622161822 | 0.976 | 0.902 | 0 |
| **RPL24** | 0 | -0.545982450265136 | 1 | 1 | 0 |
| **NFKBIZ** | 0 | -0.495488733242036 | 0.54 | 0.875 | 0 |
| **IFT57** | 0 | 0.333873003871385 | 0.956 | 0.854 | 0 |
| **MYH15** | 0 | 0.67492176150815 | 0.736 | 0.015 | 0 |
| **PHLDB2** | 0 | 0.620158585896076 | 0.872 | 0.253 | 0 |
| **LSAMP** | 0 | -0.277098935717593 | 0.237 | 0.692 | 0 |
| **IGSF11** | 0 | -0.270987633282438 | 0.04 | 0.618 | 0 |
| **PARP14** | 0 | 0.412381612246215 | 0.917 | 0.753 | 0 |
| **MYLK** | 0 | 0.284717697338302 | 0.668 | 0.196 | 0 |
| **ITGB5** | 0 | 0.487331414296795 | 0.914 | 0.558 | 0 |
| **HEG1** | 0 | 0.520090778901776 | 0.855 | 0.497 | 0 |
| **SLC12A8** | 0 | 0.633441882663233 | 0.774 | 0.039 | 0 |
| **CHCHD6** | 0 | -1.63678870894874 | 0.938 | 0.996 | 0 |
| **ABTB1** | 0 | 0.330435978695252 | 0.757 | 0.262 | 0 |
| **MRPL3** | 0 | -0.467033670987121 | 0.976 | 0.996 | 0 |
| **BFSP2** | 0 | -0.26515950544887 | 0.034 | 0.487 | 0 |
| **ANAPC13** | 0 | 0.32202701132005 | 0.945 | 0.835 | 0 |
| **PLOD2** | 0 | 0.485172611731459 | 0.843 | 0.628 | 0 |
| **TIPARP** | 0 | 0.491141508224689 | 0.87 | 0.515 | 0 |
| **LXN** | 0 | -0.29247428883173 | 0.472 | 0.777 | 0 |
| **SKIL** | 0 | 0.280385440744248 | 0.682 | 0.208 | 0 |
| **PARL** | 0 | -0.418972129985748 | 0.828 | 0.957 | 0 |
| **ETV5** | 0 | -0.814812997398271 | 0.933 | 0.993 | 0 |
| **EIF4A2** | 0 | 0.555248053325968 | 0.996 | 0.989 | 0 |
| **LPP** | 0 | 0.354399157107832 | 0.875 | 0.59 | 0 |
| **IL1RAP** | 0 | -0.449901078853237 | 0.517 | 0.861 | 0 |
| **CCDC50** | 0 | -0.467886714655646 | 0.964 | 0.988 | 0 |

| | | | | | |
|---|---|---|---|---|---|
| **OPA1** | 0 | -0.335056185349081 | 0.764 | 0.904 | 0 |
| **CEP19** | 0 | 0.320336760224692 | 0.764 | 0.258 | 0 |
| **DLG1** | 0 | 0.355473209181684 | 0.877 | 0.633 | 0 |
| **MXD4** | 0 | 0.374233174521159 | 0.854 | 0.535 | 0 |
| **RGS12** | 0 | -0.393979320179663 | 0.344 | 0.775 | 0 |
| **LRPAP1** | 0 | -0.71275698035664 | 0.98 | 0.995 | 0 |
| **AL590235.2** | 0 | -0.268971003182621 | 0.05 | 0.61 | 0 |
| **LYAR** | 0 | -0.465232609867915 | 0.856 | 0.959 | 0 |
| **QDPR** | 0 | -0.559844822230312 | 0.84 | 0.971 | 0 |
| **SOD3** | 0 | 0.480302126927336 | 0.817 | 0.469 | 0 |
| **RBPJ** | 0 | 0.818515602180281 | 0.966 | 0.877 | 0 |
| **FAM114A1** | 0 | 0.516725541607953 | 0.96 | 0.825 | 0 |
| **SMIM14** | 0 | 0.394719835987857 | 0.799 | 0.278 | 0 |
| **CHIC2** | 0 | 0.251023909326557 | 0.732 | 0.348 | 0 |
| **PDGFRA** | 0 | 0.500221706120982 | 0.67 | 0.039 | 0 |
| **IGFBP7** | 0 | 3.78888601716872 | 0.995 | 0.285 | 0 |
| **UGT2B7** | 0 | 0.744600664125897 | 0.934 | 0.517 | 0 |
| **RUFY3** | 0 | -0.323496650544464 | 0.462 | 0.798 | 0 |
| **EREG** | 0 | 0.82394591821004 | 0.46 | 0.017 | 0 |
| **ANTXR2** | 0 | 0.36565192476399 | 0.685 | 0.09 | 0 |
| **PTPN13** | 0 | 0.698234163567505 | 0.919 | 0.537 | 0 |
| **SPP1** | 0 | 2.54380630685575 | 0.949 | 0.182 | 0 |
| **PKD2** | 0 | 0.319101694078276 | 0.791 | 0.399 | 0 |
| **SNCA** | 0 | -0.511735183237716 | 0.837 | 0.97 | 0 |
| **TSPAN5** | 0 | 0.434358485235405 | 0.808 | 0.27 | 0 |
| **H2AFZ** | 0 | -1.46360507896572 | 1 | 1 | 0 |
| **LEF1** | 0 | -0.504700794410253 | 0.587 | 0.889 | 0 |
| **ZNF330** | 0 | -0.367925658424499 | 0.716 | 0.916 | 0 |
| **INPP4B** | 0 | -0.470010317054273 | 0.715 | 0.938 | 0 |
| **RPS3A** | 0 | -0.486810448376037 | 1 | 1 | 0 |
| **TRIM2** | 0 | -0.75327792185255 | 0.425 | 0.929 | 0 |
| **TDO2** | 0 | 0.258812624387124 | 0.455 | 0.101 | 0 |

| | | | | | |
|---|---|---|---|---|---|
| **FNIP2** | 0 | -0.438390504102621 | 0.496 | 0.861 | 0 |
| **HPGD** | 0 | -0.565533497692458 | 0.076 | 0.758 | 0 |
| **VEGFC** | 0 | 0.745272257566158 | 0.84 | 0.093 | 0 |
| **SLC25A4** | 0 | -0.757774271012737 | 0.766 | 0.977 | 0 |
| **AHRR** | 0 | 0.344714066765824 | 0.819 | 0.303 | 0 |
| **ADCY2** | 0 | -0.298888874324408 | 0.096 | 0.647 | 0 |
| **CCT5** | 0 | -0.717441689148925 | 0.985 | 0.998 | 0 |
| **DAP** | 0 | 0.418537918594928 | 0.953 | 0.862 | 0 |
| **ANKH** | 0 | 0.490518246059999 | 0.864 | 0.451 | 0 |
| **MYO10** | 0 | -0.633032517812579 | 0.997 | 0.999 | 0 |
| **BASP1** | 0 | 1.60229488451094 | 0.979 | 0.151 | 0 |
| **SLC45A2** | 0 | -0.596184103527801 | 0.475 | 0.884 | 0 |
| **OSMR** | 0 | 0.401375093374024 | 0.837 | 0.354 | 0 |
| **ITGA1** | 0 | 0.253452075496261 | 0.54 | 0.082 | 0 |
| **ITGA2** | 0 | 0.896671492583073 | 0.715 | 0.216 | 0 |
| **GPX8** | 0 | 0.282370707835042 | 0.838 | 0.601 | 0 |
| **IL6ST** | 0 | 1.06455043444391 | 0.95 | 0.555 | 0 |
| **MAP1B** | 0 | 0.526897681601139 | 0.646 | 0.058 | 0 |
| **FOXD1** | 0 | 0.273845223752679 | 0.703 | 0.317 | 0 |
| **F2R** | 0 | 0.883388818435658 | 0.898 | 0.153 | 0 |
| **TBCA** | 0 | -0.394864033220221 | 0.994 | 0.998 | 0 |
| **ZFYVE16** | 0 | -0.439736334967403 | 0.772 | 0.91 | 0 |
| **RASA1** | 0 | 0.31320852556266 | 0.763 | 0.386 | 0 |
| **POLR3G** | 0 | -0.51429655415686 | 0.737 | 0.935 | 0 |
| **NR2F1** | 0 | 0.386742832952467 | 0.633 | 0.066 | 0 |
| **GLRX** | 0 | 0.555797072166135 | 0.932 | 0.769 | 0 |
| **CAST** | 0 | 0.519571270152789 | 0.997 | 0.986 | 0 |
| **PAM** | 0 | 0.360135200450314 | 0.805 | 0.404 | 0 |
| **CAMK4** | 0 | -0.337160014780311 | 0.542 | 0.825 | 0 |
| **SEMA6A** | 0 | -0.571575975284202 | 0.26 | 0.859 | 0 |
| **HINT1** | 0 | -0.566799952066451 | 0.999 | 0.999 | 0 |
| **P4HA2** | 0 | 0.322246804586374 | 0.834 | 0.549 | 0 |

| | | | | | |
|---|---|---|---|---|---|
| **IRF1** | 0 | 0.480958612366876 | 0.778 | 0.262 | 0 |
| **SPOCK1** | 0 | 0.907844549162507 | 0.726 | 0.019 | 0 |
| **CYSTM1** | 0 | -0.511948400826616 | 0.779 | 0.964 | 0 |
| **GNPDA1** | 0 | -0.315270387329254 | 0.756 | 0.91 | 0 |
| **FGF1** | 0 | 0.387399322597625 | 0.694 | 0.096 | 0 |
| **NR3C1** | 0 | 0.363334154747855 | 0.907 | 0.702 | 0 |
| **STK32A** | 0 | -0.397176624013136 | 0.064 | 0.722 | 0 |
| **RPS14** | 0 | -0.529401992617228 | 1 | 1 | 0 |
| **SMIM3** | 0 | 0.31378685381463 | 0.685 | 0.174 | 0 |
| **SPARC** | 0 | -0.579719706787046 | 0.941 | 0.99 | 0 |
| **ATOX1** | 0 | -1.96384672156024 | 0.998 | 1 | 0 |
| **G3BP1** | 0 | -0.80330315387674 | 0.924 | 0.989 | 0 |
| **PRELID1** | 0 | -0.47578915619122 | 0.998 | 0.999 | 0 |
| **NHP2** | 0 | -0.515670504198304 | 0.971 | 0.993 | 0 |
| **HNRNPAB** | 0 | -0.694066930834147 | 0.966 | 0.994 | 0 |
| **RNF130** | 0 | -0.300192569957215 | 0.726 | 0.892 | 0 |
| **TFAP2A** | 0 | -0.504791451112477 | 0.715 | 0.902 | 0 |
| **PHACTR1** | 0 | -0.681903052539988 | 0.35 | 0.899 | 0 |
| **TBC1D7** | 0 | -0.590994481872982 | 0.514 | 0.911 | 0 |
| **DTNBP1** | 0 | -0.28164561056051 | 0.663 | 0.858 | 0 |
| **ID4** | 0 | -0.254504276409688 | 0.103 | 0.547 | 0 |
| **HLA-A** | 0 | 1.39003630717932 | 0.997 | 0.972 | 0 |
| **PPP1R18** | 0 | 0.443594294508057 | 0.949 | 0.818 | 0 |
| **IER3** | 0 | 2.09243879971642 | 0.999 | 0.955 | 0 |
| **HLA-C** | 0 | 0.965509756023919 | 0.995 | 0.985 | 0 |
| **HLA-B** | 0 | 1.20456412037235 | 0.972 | 0.66 | 0 |
| **HMGA1** | 0 | 1.28340996500107 | 0.995 | 0.988 | 0 |
| **SMIM29** | 0 | 0.516947590458248 | 0.941 | 0.757 | 0 |
| **CDKN1A** | 0 | 0.370761683579327 | 0.668 | 0.056 | 0 |
| **DNPH1** | 0 | -0.359637002094079 | 0.744 | 0.911 | 0 |
| **YIPF3** | 0 | 0.343823186351805 | 0.951 | 0.831 | 0 |
| **VEGFA** | 0 | 1.64197843099717 | 0.963 | 0.368 | 0 |

| | | | | | |
|---|---|---|---|---|---|
| **PTP4A1** | 0 | -0.37254739208871 | 0.7 | 0.898 | 0 |
| **ME1** | 0 | 1.74170924204918 | 0.981 | 0.603 | 0 |
| **PRSS35** | 0 | 1.24509501135168 | 0.789 | 0.056 | 0 |
| **NT5E** | 0 | 1.2131009049105 | 0.971 | 0.596 | 0 |
| **RRAGD** | 0 | -0.520852094744843 | 0.387 | 0.893 | 0 |
| **SCML4** | 0 | -0.351992818863105 | 0.044 | 0.616 | 0 |
| **LAMA4** | 0 | -0.269568022679888 | 0.171 | 0.67 | 0 |
| **FABP7** | 0 | -0.774906025244765 | 0.355 | 0.89 | 0 |
| **HDDC2** | 0 | -0.446766317623653 | 0.951 | 0.988 | 0 |
| **ENPP1** | 0 | 1.25617589321994 | 0.852 | 0.57 | 0 |
| **STX7** | 0 | -0.32592025001134 | 0.687 | 0.889 | 0 |
| **SGK1** | 0 | -0.965976430209428 | 0.743 | 0.977 | 0 |
| **NHSL1** | 0 | -0.260116873449073 | 0.293 | 0.693 | 0 |
| **CITED2** | 0 | 0.436837930624661 | 0.724 | 0.284 | 0 |
| **RAB32** | 0 | -0.735938693831585 | 0.894 | 0.985 | 0 |
| **AKAP12** | 0 | -0.930391701473612 | 0.549 | 0.97 | 0 |
| **RGS17** | 0 | 0.296551612633757 | 0.727 | 0.366 | 0 |
| **SNX9** | 0 | 0.654464485306171 | 0.988 | 0.943 | 0 |
| **SYNJ2** | 0 | 0.539608956833017 | 0.859 | 0.409 | 0 |
| **PDGFA** | 0 | 0.541854591272691 | 0.853 | 0.266 | 0 |
| **PRKAR1B** | 0 | -0.476204915213994 | 0.813 | 0.94 | 0 |
| **TTYH3** | 0 | -0.525710214633351 | 0.628 | 0.89 | 0 |
| **RADIL** | 0 | -0.30753690573114 | 0.08 | 0.656 | 0 |
| **ACTB** | 0 | 0.528991622562863 | 1 | 1 | 0 |
| **FSCN1** | 0 | 0.34997475133359 | 0.641 | 0.024 | 0 |
| **CYTH3** | 0 | -0.27149006902988 | 0.425 | 0.769 | 0 |
| **ARL4A** | 0 | 0.448062000923259 | 0.913 | 0.741 | 0 |
| **ETV1** | 0 | 0.877632533689212 | 0.966 | 0.575 | 0 |
| **BZW2** | 0 | -0.595239248185595 | 0.857 | 0.975 | 0 |
| **AHR** | 0 | 0.259802127353557 | 0.68 | 0.354 | 0 |
| **FAM126A** | 0 | 0.673869768547561 | 0.915 | 0.676 | 0 |
| **IGF2BP3** | 0 | 0.52313588459273 | 0.916 | 0.819 | 0 |

| | | | | | |
|---|---|---|---|---|---|
| **CYCS** | 0 | -0.628984965964172 | 0.993 | 0.999 | 0 |
| **SNX10** | 0 | -0.548551832923041 | 0.221 | 0.83 | 0 |
| **GGCT** | 0 | -0.740136491230687 | 0.744 | 0.972 | 0 |
| **STK17A** | 0 | 0.476753816102292 | 0.951 | 0.823 | 0 |
| **TBRG4** | 0 | -0.453641996907915 | 0.872 | 0.965 | 0 |
| **EGFR** | 0 | 0.515664652139782 | 0.782 | 0.145 | 0 |
| **CHCHD2** | 0 | -0.546653312381579 | 0.999 | 1 | 0 |
| **STX1A** | 0 | 0.550688064883972 | 0.828 | 0.147 | 0 |
| **CLIP2** | 0 | 0.306294002864882 | 0.777 | 0.319 | 0 |
| **RHBDD2** | 0 | 0.325125489875011 | 0.84 | 0.539 | 0 |
| **STEAP1** | 0 | 0.494048880050135 | 0.697 | 0.342 | 0 |
| **CDK6** | 0 | -0.528121390058675 | 0.591 | 0.927 | 0 |
| **SAMD9L** | 0 | 0.362894351755398 | 0.69 | 0.261 | 0 |
| **TFPI2** | 0 | 1.86191511490462 | 0.878 | 0.328 | 0 |
| **GNG11** | 0 | 2.09697522802254 | 0.999 | 0.935 | 0 |
| **BET1** | 0 | 0.340065841291227 | 0.92 | 0.764 | 0 |
| **SGCE** | 0 | 0.385424463631985 | 0.944 | 0.805 | 0 |
| **SLC25A13** | 0 | -0.814189494923299 | 0.785 | 0.983 | 0 |
| **BAIAP2L1** | 0 | -0.50403074283134 | 0.727 | 0.951 | 0 |
| **TSC22D4** | 0 | 0.601504712174929 | 0.958 | 0.852 | 0 |
| **PCOLCE** | 0 | 1.83476989243361 | 0.942 | 0.441 | 0 |
| **TRIM56** | 0 | 0.53961618831995 | 0.972 | 0.877 | 0 |
| **SERPINE1** | 0 | 0.590637302753445 | 0.57 | 0.04 | 0 |
| **AP1S1** | 0 | 0.602895925189499 | 1 | 0.996 | 0 |
| **VGF** | 0 | 0.881243630015587 | 0.99 | 0.864 | 0 |
| **ARMC10** | 0 | -0.342949176956742 | 0.933 | 0.979 | 0 |
| **KMT2E** | 0 | 0.778852333526851 | 0.998 | 0.969 | 0 |
| **CCDC71L** | 0 | 0.493425453605851 | 0.882 | 0.543 | 0 |
| **LAMB1** | 0 | 0.589298714384126 | 0.968 | 0.839 | 0 |
| **PNPLA8** | 0 | 0.313660074177915 | 0.924 | 0.777 | 0 |
| **IMMP2L** | 0 | -0.298102337246651 | 0.355 | 0.746 | 0 |
| **CAV2** | 0 | 0.557098273339286 | 0.916 | 0.579 | 0 |

| | | | | | |
|---|---|---|---|---|---|
| **CAV1** | 0 | 2.74177106931384 | 0.998 | 0.91 | 0 |
| **CAPZA2** | 0 | 0.336228690249656 | 0.991 | 0.977 | 0 |
| **TSPAN33** | 0 | -0.289444811528126 | 0.306 | 0.742 | 0 |
| **CPA4** | 0 | 0.302656619033979 | 0.311 | 0.006 | 0 |
| **AC016831.7** | 0 | 0.539242145313187 | 0.84 | 0.347 | 0 |
| **LINC00513** | 0 | 0.293032617530175 | 0.675 | 0.288 | 0 |
| **AC016831.5** | 0 | 0.668136413554165 | 0.921 | 0.629 | 0 |
| **AC058791.1** | 0 | 0.55312296793979 | 0.831 | 0.447 | 0 |
| **MKLN1** | 0 | 0.486611281379982 | 0.95 | 0.848 | 0 |
| **AKR1B1** | 0 | 2.60472491099998 | 1 | 0.993 | 0 |
| **CALD1** | 0 | 0.859567686250607 | 1 | 1 | 0 |
| **AC093673.1** | 0 | 0.268884064268092 | 0.68 | 0.145 | 0 |
| **ZYX** | 0 | 0.564407278074598 | 0.931 | 0.653 | 0 |
| **INSIG1** | 0 | 0.883840995550952 | 0.947 | 0.854 | 0 |
| **DNAJB6** | 0 | 0.429572567917886 | 0.987 | 0.97 | 0 |
| **IL3RA** | 0 | 0.279128185154691 | 0.511 | 0.04 | 0 |
| **GYG2** | 0 | -0.260678851518853 | 0.128 | 0.673 | 0 |
| **TBL1X** | 0 | 0.401774245451946 | 0.807 | 0.331 | 0 |
| **GPR143** | 0 | -0.451848665169008 | 0.764 | 0.956 | 0 |
| **TMSB4X** | 0 | 2.62208378554497 | 0.999 | 0.992 | 0 |
| **GPM6B** | 0 | -1.77449211036024 | 0.645 | 0.997 | 0 |
| **ASB9** | 0 | -0.29114224699329 | 0.527 | 0.819 | 0 |
| **PIR** | 0 | -1.33692231919184 | 0.47 | 0.984 | 0 |
| **AP1S2** | 0 | -0.53168531400757 | 0.979 | 0.996 | 0 |
| **SH3KBP1** | 0 | 0.538010428264927 | 0.989 | 0.97 | 0 |
| **SMS** | 0 | -1.05475431641457 | 0.957 | 0.996 | 0 |
| **SAT1** | 0 | -1.87278712147706 | 0.917 | 0.997 | 0 |
| **MIR222HG** | 0 | 0.379878231731774 | 0.705 | 0.18 | 0 |
| **TIMP1** | 0 | 1.74152683168107 | 0.999 | 0.987 | 0 |
| **PLP2** | 0 | 0.763208465324901 | 0.995 | 0.971 | 0 |
| **HSD17B10** | 0 | -0.388956021666486 | 0.884 | 0.963 | 0 |
| **MSN** | 0 | -0.550818058638818 | 0.981 | 0.996 | 0 |

| | | | | | |
|---|---|---|---|---|---|
| **GJB1** | 0 | -0.325068712792425 | 0.028 | 0.626 | 0 |
| **CITED1** | 0 | -0.497155283753857 | 0.14 | 0.69 | 0 |
| **SRPX2** | 0 | 0.269243921548281 | 0.611 | 0.015 | 0 |
| **TCEAL9** | 0 | 0.579617561651914 | 0.974 | 0.865 | 0 |
| **BEX3** | 0 | 0.511508012789656 | 0.997 | 0.994 | 0 |
| **TCEAL3** | 0 | 0.346971287894457 | 0.884 | 0.618 | 0 |
| **PLP1** | 0 | -0.919891222726677 | 0.37 | 0.97 | 0 |
| **ACSL4** | 0 | 0.647532840006857 | 0.948 | 0.771 | 0 |
| **SLC25A5** | 0 | -0.78787358926908 | 0.995 | 0.999 | 0 |
| **SEPT6** | 0 | -0.383411304412101 | 0.32 | 0.785 | 0 |
| **RTL8C** | 0 | 0.3199170869675 | 0.913 | 0.772 | 0 |
| **CETN2** | 0 | 0.335074303861816 | 0.902 | 0.735 | 0 |
| **BGN** | 0 | 1.43004284715093 | 0.767 | 0.047 | 0 |
| **SLC6A8** | 0 | -0.472019790904531 | 0.845 | 0.964 | 0 |
| **L1CAM** | 0 | 0.89271031958444 | 0.994 | 0.957 | 0 |
| **FLNA** | 0 | 1.0598216903058 | 0.997 | 0.98 | 0 |
| **LAGE3** | 0 | -0.384966750332074 | 0.984 | 0.994 | 0 |
| **DKC1** | 0 | -0.525133508445343 | 0.954 | 0.989 | 0 |
| **AC100810.1** | 0 | 0.421236624591072 | 0.964 | 0.88 | 0 |
| **GATA4** | 0 | -0.263559238579496 | 0.27 | 0.689 | 0 |
| **CTSB** | 0 | 1.02275796790928 | 0.999 | 0.984 | 0 |
| **LONRF1** | 0 | -0.504017230635564 | 0.448 | 0.886 | 0 |
| **ASAH1** | 0 | -0.889438452486181 | 0.932 | 0.995 | 0 |
| **PSD3** | 0 | 0.498509708028268 | 0.848 | 0.323 | 0 |
| **LZTS1** | 0 | -0.583105524755738 | 0.671 | 0.947 | 0 |
| **PDLIM2** | 0 | 0.451301716086221 | 0.987 | 0.968 | 0 |
| **BIN3** | 0 | -0.474730310524303 | 0.68 | 0.912 | 0 |
| **LOXL2** | 0 | 0.544050888240304 | 0.779 | 0.148 | 0 |
| **SLC25A37** | 0 | 0.607155081438731 | 0.985 | 0.913 | 0 |
| **STC1** | 0 | 1.96083387010078 | 0.932 | 0.09 | 0 |
| **BNIP3L** | 0 | 0.702428177102345 | 0.98 | 0.859 | 0 |
| **DPYSL2** | 0 | 0.720744845215905 | 0.886 | 0.106 | 0 |

| | | | | | |
|---|---|---|---|---|---|
| **NRG1** | 0 | 0.280439363292428 | 0.441 | 0.039 | 0 |
| **EIF4EBP1** | 0 | -0.522453269292678 | 0.966 | 0.994 | 0 |
| **FGFR1** | 0 | 0.52668564553924 | 0.874 | 0.437 | 0 |
| **SFRP1** | 0 | 1.58654763875472 | 0.947 | 0.44 | 0 |
| **PLAT** | 0 | -0.421087845470451 | 0.602 | 0.865 | 0 |
| **PCMTD1** | 0 | 0.376257917135437 | 0.875 | 0.6 | 0 |
| **RGS20** | 0 | -1.04419500855045 | 0.566 | 0.97 | 0 |
| **CA8** | 0 | -0.450890487799448 | 0.317 | 0.763 | 0 |
| **CHD7** | 0 | -0.593618092507964 | 0.567 | 0.929 | 0 |
| **CYP7B1** | 0 | -0.371573517015398 | 0.093 | 0.704 | 0 |
| **SGK3** | 0 | -0.406699238828289 | 0.332 | 0.802 | 0 |
| **RPL7** | 0 | -0.589727942272147 | 1 | 1 | 0 |
| **TPD52** | 0 | -0.560002029050513 | 0.754 | 0.961 | 0 |
| **PAG1** | 0 | 0.554607731787564 | 0.921 | 0.671 | 0 |
| **FABP5** | 0 | -1.5826072597609 | 0.928 | 1 | 0 |
| **PMP2** | 0 | -0.329383310915909 | 0.059 | 0.586 | 0 |
| **ATP6V0D2** | 0 | -0.619812017146191 | 0.319 | 0.737 | 0 |
| **TMEM64** | 0 | -0.296477150199594 | 0.493 | 0.774 | 0 |
| **GEM** | 0 | 0.292896142497444 | 0.57 | 0.189 | 0 |
| **ESRP1** | 0 | -0.41055581632471 | 0.065 | 0.748 | 0 |
| **RNF19A** | 0 | -0.475475955619149 | 0.789 | 0.933 | 0 |
| **CTHRC1** | 0 | 0.747964406333798 | 0.987 | 0.955 | 0 |
| **TRPS1** | 0 | 0.349835704773608 | 0.816 | 0.471 | 0 |
| **EXT1** | 0 | 0.616599898286763 | 0.976 | 0.862 | 0 |
| **NOV** | 0 | 0.692285171369325 | 0.888 | 0.414 | 0 |
| **SNTB1** | 0 | 0.710904038863371 | 0.902 | 0.393 | 0 |
| **TMEM65** | 0 | 0.363646854741464 | 0.833 | 0.515 | 0 |
| **MTSS1** | 0 | 0.836446479210952 | 0.864 | 0.08 | 0 |
| **MYC** | 0 | -0.980654000709121 | 0.672 | 0.936 | 0 |
| **EFR3A** | 0 | 0.289440182449814 | 0.881 | 0.695 | 0 |
| **WISP1** | 0 | 0.451028569413233 | 0.471 | 0.005 | 0 |
| **NDRG1** | 0 | 1.06160483215693 | 0.929 | 0.212 | 0 |

| | | | | | |
|---|---|---|---|---|---|
| **TSNARE1** | 0 | 0.609476135065985 | 0.787 | 0.175 | 0 |
| **PLEC** | 0 | 0.584458394634721 | 0.985 | 0.91 | 0 |
| **EXOSC4** | 0 | -0.35115119300517 | 0.897 | 0.963 | 0 |
| **CYC1** | 0 | -0.552655393747527 | 0.995 | 0.999 | 0 |
| **BOP 1.00** | 0 | -0.509964442211086 | 0.821 | 0.964 | 0 |
| **VPS28** | 0 | 0.57578855178476 | 0.993 | 0.984 | 0 |
| **RPL8** | 0 | -0.564657306114576 | 1 | 1 | 0 |
| **KANK1** | 0 | -0.332423744816786 | 0.613 | 0.862 | 0 |
| **MLANA** | 0 | -3.49055492791187 | 0.572 | 0.999 | 0 |
| **LURAP1L** | 0 | 0.277375217539619 | 0.59 | 0.104 | 0 |
| **CDKN2B** | 0 | 0.389973184032513 | 0.702 | 0.349 | 0 |
| **TLN1** | 0 | 0.983597178466606 | 0.995 | 0.957 | 0 |
| **CREB3** | 0 | 0.290819235508137 | 0.864 | 0.681 | 0 |
| **EBLN3P** | 0 | 0.287923248299761 | 0.891 | 0.705 | 0 |
| **ALDH1B1** | 0 | -0.31583801674377 | 0.354 | 0.762 | 0 |
| **ANXA1** | 0 | 1.28241657781364 | 0.915 | 0.149 | 0 |
| **PSAT1** | 0 | -0.288682447661626 | 0.459 | 0.801 | 0 |
| **GOLM1** | 0 | -0.411754302202833 | 0.597 | 0.905 | 0 |
| **CTSL** | 0 | 0.938142507193101 | 0.96 | 0.821 | 0 |
| **NFIL3** | 0 | 0.307935864982191 | 0.734 | 0.353 | 0 |
| **CARD19** | 0 | 0.481091221846007 | 0.957 | 0.797 | 0 |
| **COL15A1** | 0 | 0.659117258945527 | 0.781 | 0.289 | 0 |
| **TGFBR1** | 0 | 1.1058464453633 | 0.96 | 0.693 | 0 |
| **BAAT** | 0 | -0.535693080149815 | 0.066 | 0.771 | 0 |
| **SLC44A1** | 0 | -0.494621215014152 | 0.926 | 0.982 | 0 |
| **TXN** | 0 | -0.56516120058369 | 0.998 | 0.999 | 0 |
| **TNC** | 0 | 0.41082657495728 | 0.729 | 0.178 | 0 |
| **GSN** | 0 | -0.572787692622675 | 0.942 | 0.986 | 0 |
| **STOM** | 0 | -0.348764683533909 | 0.741 | 0.897 | 0 |
| **PSMB7** | 0 | -0.466863609180151 | 0.977 | 0.991 | 0 |
| **RPL35** | 0 | -0.621416170400777 | 0.999 | 1 | 0 |
| **ANGPTL2** | 0 | -0.320922559554123 | 0.351 | 0.752 | 0 |

| | | | | | |
|---|---|---|---|---|---|
| **FAM129B** | 0 | 0.354168098432046 | 0.845 | 0.552 | 0 |
| **SET** | 0 | -0.585556919679924 | 0.998 | 0.999 | 0 |
| **IER5L** | 0 | 0.758574756709192 | 0.846 | 0.398 | 0 |
| **RPL7A** | 0 | -0.53970792549854 | 1 | 1 | 0 |
| **EGFL7** | 0 | -0.266492737505208 | 0.327 | 0.729 | 0 |
| **FAM69B** | 0 | -0.457826767969116 | 0.14 | 0.804 | 0 |
| **RABL6** | 0 | -0.394759205806089 | 0.93 | 0.977 | 0 |
| **IFITM3** | 0 | 0.781032269894808 | 0.941 | 0.696 | 0 |
| **CD151** | 0 | 0.506221447299851 | 0.992 | 0.971 | 0 |
| **CD81** | 0 | 0.727741483942988 | 0.999 | 0.997 | 0 |
| **PHLDA2** | 0 | 1.17470327047048 | 0.991 | 0.921 | 0 |
| **CAVIN3** | 0 | 1.15999169111987 | 0.983 | 0.862 | 0 |
| **ADM** | 0 | 1.43380178510752 | 0.943 | 0.505 | 0 |
| **MICAL2** | 0 | 0.257089107154805 | 0.626 | 0.079 | 0 |
| **PDE3B** | 0 | -0.520052670718282 | 0.167 | 0.827 | 0 |
| **SOX6** | 0 | -0.728014725741856 | 0.209 | 0.929 | 0 |
| **NUCB2** | 0 | 0.762194718853448 | 0.978 | 0.894 | 0 |
| **BDNF** | 0 | 0.408251958132605 | 0.643 | 0.012 | 0 |
| **RCN1** | 0 | 0.888273007733825 | 0.995 | 0.968 | 0 |
| **CD44** | 0 | 1.91623556359347 | 1 | 0.986 | 0 |
| **C11orf96** | 0 | -0.436829079728634 | 0.14 | 0.6 | 0 |
| **CD82** | 0 | 0.834025897434929 | 0.947 | 0.566 | 0 |
| **PTPRJ** | 0 | -0.26114981623381 | 0.178 | 0.677 | 0 |
| **FTH1** | 0 | 0.995839642751749 | 1 | 1 | 0 |
| **ASRGL1** | 0 | -0.795607355020852 | 0.241 | 0.898 | 0 |
| **AHNAK** | 0 | 0.687634833195447 | 0.994 | 0.89 | 0 |
| **CAPN1** | 0 | 0.297581606496244 | 0.921 | 0.783 | 0 |
| **MALAT1** | 0 | 2.10675169838412 | 1 | 0.998 | 0 |
| **LTBP3** | 0 | 0.302057145725942 | 0.849 | 0.572 | 0 |
| **FAM89B** | 0 | 0.532455094897021 | 0.935 | 0.711 | 0 |
| **CCDC85B** | 0 | 1.05969689922727 | 1 | 0.997 | 0 |
| **C11orf68** | 0 | 0.352303408189869 | 0.847 | 0.578 | 0 |

| | | | | | |
|---|---|---|---|---|---|
| **DRAP1** | 0 | 0.507637891198755 | 0.992 | 0.986 | 0 |
| **RIN1** | 0 | 0.287120132396531 | 0.701 | 0.34 | 0 |
| **MRPL11** | 0 | -0.432702316724857 | 0.95 | 0.986 | 0 |
| **C11orf24** | 0 | -0.377224697758065 | 0.906 | 0.972 | 0 |
| **MYEOV** | 0 | 0.559507504440731 | 0.716 | 0.168 | 0 |
| **CCND1** | 0 | 2.94703174934513 | 0.999 | 0.994 | 0 |
| **NUMA1** | 0 | 0.436176572355349 | 0.995 | 0.98 | 0 |
| **STARD10** | 0 | 0.526873363044706 | 0.985 | 0.957 | 0 |
| **PGM2L1** | 0 | 0.401092070640993 | 0.695 | 0.286 | 0 |
| **GAB2** | 0 | -0.368888794379437 | 0.248 | 0.763 | 0 |
| **NARS2** | 0 | -0.441473168174721 | 0.696 | 0.919 | 0 |
| **RAB38** | 0 | -0.988989989013983 | 0.708 | 0.982 | 0 |
| **CTSC** | 0 | -0.652125676181343 | 0.931 | 0.988 | 0 |
| **TYR** | 0 | -0.785409310561435 | 0.127 | 0.92 | 0 |
| **AMOTL1** | 0 | 0.308692112830165 | 0.787 | 0.428 | 0 |
| **MMP1** | 0 | 4.30118461991786 | 0.962 | 0.258 | 0 |
| **CASP4** | 0 | 0.434242991936555 | 0.92 | 0.663 | 0 |
| **ATM** | 0 | -0.368982744864782 | 0.875 | 0.954 | 0 |
| **RDX** | 0 | -0.563294207964119 | 0.955 | 0.992 | 0 |
| **NNMT** | 0 | 1.21605327515512 | 0.772 | 0.061 | 0 |
| **REXO2** | 0 | 0.416565342005641 | 0.991 | 0.983 | 0 |
| **CADM1** | 0 | 0.862629936479143 | 0.93 | 0.721 | 0 |
| **BCL9L** | 0 | 0.282187514228161 | 0.745 | 0.343 | 0 |
| **MCAM** | 0 | -0.723449675974769 | 0.514 | 0.931 | 0 |
| **EI24** | 0 | -0.412037424196651 | 0.943 | 0.987 | 0 |
| **ST3GAL4** | 0 | -0.587313292367499 | 0.936 | 0.979 | 0 |
| **NTM** | 0 | 0.326615897603477 | 0.614 | 0.039 | 0 |
| **KIN** | 0 | -0.309347413973544 | 0.647 | 0.864 | 0 |
| **ATP5F1C** | 0 | -0.402348190035298 | 0.954 | 0.981 | 0 |
| **FRMD4A** | 0 | 0.90248099025396 | 0.974 | 0.912 | 0 |
| **VIM** | 0 | 0.8776732318872 | 1 | 1 | 0 |
| **BAMBI** | 0 | -0.510153793589776 | 0.655 | 0.898 | 0 |

| | | | | | |
|---|---|---|---|---|---|
| **SVIL** | 0 | 0.524667381633132 | 0.868 | 0.407 | 0 |
| **ITGB1** | 0 | 1.08675480600398 | 0.991 | 0.921 | 0 |
| **NRP1** | 0 | 0.585021912482474 | 0.711 | 0.013 | 0 |
| **ARHGAP22** | 0 | 0.760703352667485 | 0.869 | 0.12 | 0 |
| **DKK 1.00** | 0 | 0.408998921120521 | 0.286 | 0.031 | 0 |
| **TFAM** | 0 | -0.403859367839365 | 0.938 | 0.984 | 0 |
| **ARID5B** | 0 | 0.585016654331132 | 0.908 | 0.552 | 0 |
| **SRGN** | 0 | 2.0481073516153 | 0.925 | 0.058 | 0 |
| **DDIT4** | 0 | 1.25681580488414 | 0.983 | 0.838 | 0 |
| **VCL** | 0 | 0.652516269893088 | 0.922 | 0.66 | 0 |
| **RPS24** | 0 | -0.882307104212541 | 1 | 1 | 0 |
| **ANXA11** | 0 | 0.708831039990519 | 0.989 | 0.957 | 0 |
| **TSPAN14** | 0 | -0.552350624919674 | 0.757 | 0.953 | 0 |
| **MYOF** | 0 | 0.661201156070884 | 0.802 | 0.138 | 0 |
| **NPM3** | 0 | -0.513771553368187 | 0.852 | 0.972 | 0 |
| **NOLC1** | 0 | -0.79968465215992 | 0.916 | 0.99 | 0 |
| **TRIM8** | 0 | 0.665516427660118 | 0.922 | 0.545 | 0 |
| **ARL 3.00** | 0 | 0.500438018228072 | 0.983 | 0.941 | 0 |
| **BORCS7** | 0 | 0.311525376987482 | 0.902 | 0.722 | 0 |
| **SFR1** | 0 | 0.2516374112809 | 0.701 | 0.261 | 0 |
| **GSTO1** | 0 | -0.763672471617966 | 0.999 | 0.999 | 0 |
| **MXI1** | 0 | -0.568527318956888 | 0.771 | 0.974 | 0 |
| **RGS10** | 0 | -0.814588270726257 | 0.985 | 0.999 | 0 |
| **PTPRE** | 0 | 0.746632050199021 | 0.967 | 0.676 | 0 |
| **FKBP4** | 0 | -0.4760694228302 | 0.896 | 0.979 | 0 |
| **CD9** | 0 | 0.598371819049085 | 0.984 | 0.966 | 0 |
| **TNFRSF1A** | 0 | 0.456180076436201 | 0.936 | 0.745 | 0 |
| **PTMS** | 0 | 0.920311878315208 | 0.999 | 0.998 | 0 |
| **C12orf57** | 0 | 0.745595365386781 | 0.999 | 0.986 | 0 |
| **C1S** | 0 | 0.5858706003272 | 0.725 | 0.241 | 0 |
| **C1R** | 0 | 0.558391686407813 | 0.77 | 0.194 | 0 |
| **LINC00937** | 0 | -0.444476134362428 | 0.172 | 0.812 | 0 |

| | | | | | |
|---|---|---|---|---|---|
| **CLEC2B** | 0 | 0.865786656351486 | 0.945 | 0.471 | 0 |
| **YBX3** | 0 | -0.520075352970679 | 0.988 | 0.998 | 0 |
| **GPRC5A** | 0 | -0.279390539629814 | 0.213 | 0.666 | 0 |
| **LDHB** | 0 | -0.485615788625944 | 0.999 | 1 | 0 |
| **SOX5** | 0 | -0.296506829865701 | 0.412 | 0.786 | 0 |
| **RASSF8** | 0 | 1.35332077679135 | 0.993 | 0.951 | 0 |
| **BHLHE41** | 0 | -0.542711560326205 | 0.686 | 0.918 | 0 |
| **PPFIBP1** | 0 | 0.528770497449753 | 0.995 | 0.987 | 0 |
| **BICD1** | 0 | 0.373947790941486 | 0.857 | 0.549 | 0 |
| **KIF21A** | 0 | -0.271986303898356 | 0.463 | 0.79 | 0 |
| **ZCRB1** | 0 | 0.379146357468163 | 0.988 | 0.957 | 0 |
| **SLC38A2** | 0 | 0.727529569631224 | 0.961 | 0.782 | 0 |
| **TUBA1A** | 0 | 0.471483899098513 | 0.873 | 0.625 | 0 |
| **SMARCD1** | 0 | -0.377358571207811 | 0.869 | 0.957 | 0 |
| **LIMA1** | 0 | 0.465514871646874 | 0.98 | 0.951 | 0 |
| **SMAGP** | 0 | 0.855228255778047 | 0.925 | 0.249 | 0 |
| **IGFBP6** | 0 | 0.621732319026283 | 0.833 | 0.415 | 0 |
| **ITGA5** | 0 | 0.51340156393071 | 0.844 | 0.317 | 0 |
| **GDF11** | 0 | -0.326845416781387 | 0.669 | 0.872 | 0 |
| **PMEL** | 0 | -3.77679224070717 | 0.747 | 0.997 | 0 |
| **CDK2** | 0 | -1.02484142949113 | 0.539 | 0.954 | 0 |
| **ERBB3** | 0 | -0.798603392485385 | 0.284 | 0.931 | 0 |
| **PA2G4** | 0 | -0.618275328509365 | 0.987 | 0.997 | 0 |
| **MYL6B** | 0 | 0.651848311967494 | 0.992 | 0.976 | 0 |
| **MYL6** | 0 | 0.937574563307761 | 0.999 | 0.996 | 0 |
| **LRP1** | 0 | 0.67928175280879 | 0.947 | 0.513 | 0 |
| **SHMT2** | 0 | -0.40774657511754 | 0.902 | 0.978 | 0 |
| **HMGA2** | 0 | 1.17695808640648 | 0.986 | 0.846 | 0 |
| **TMBIM4** | 0 | 0.432165884333457 | 0.958 | 0.847 | 0 |
| **CPM** | 0 | -0.325892292877472 | 0.447 | 0.824 | 0 |
| **PHLDA1** | 0 | 1.60287318345866 | 0.999 | 0.984 | 0 |
| **E2F7** | 0 | 0.438951702406432 | 0.808 | 0.394 | 0 |

| | | | | | |
|---|---|---|---|---|---|
| **DUSP6** | 0 | 0.728030218566604 | 0.974 | 0.836 | 0 |
| **ATP2B1** | 0 | 1.02226901200324 | 0.988 | 0.897 | 0 |
| **ATP2B1-AS1** | 0 | 0.339302783024228 | 0.7 | 0.175 | 0 |
| **ELK3** | 0 | 0.377515975528078 | 0.864 | 0.589 | 0 |
| **GNPTAB** | 0 | -0.384434595372236 | 0.768 | 0.939 | 0 |
| **DRAM1** | 0 | 0.371094167056661 | 0.772 | 0.291 | 0 |
| **C12orf75** | 0 | 1.58726617684362 | 0.99 | 0.933 | 0 |
| **ISCU** | 0 | 0.377126378178288 | 0.969 | 0.926 | 0 |
| **SSH1** | 0 | 0.663253552491071 | 0.958 | 0.714 | 0 |
| **SH2B3** | 0 | 0.964660686009729 | 0.961 | 0.733 | 0 |
| **RPL6** | 0 | -0.434209161162044 | 1 | 1 | 0 |
| **PEBP1** | 0 | -0.743201784518077 | 0.998 | 1 | 0 |
| **CIT** | 0 | -0.500944730252004 | 0.529 | 0.868 | 0 |
| **MLXIP** | 0 | -0.43429278253112 | 0.734 | 0.941 | 0 |
| **SCARB1** | 0 | -0.802596980252142 | 0.622 | 0.966 | 0 |
| **UBC** | 0 | 1.31624707273053 | 0.999 | 0.986 | 0 |
| **RAN** | 0 | -0.582023540641143 | 0.999 | 1 | 0 |
| **AL161772.1** | 0 | -0.383266930027832 | 0.411 | 0.817 | 0 |
| **SAP18** | 0 | -0.44352820751899 | 0.997 | 0.998 | 0 |
| **MRPL57** | 0 | -0.379423294940495 | 0.994 | 0.996 | 0 |
| **HMGB1** | 0 | -0.557649142398162 | 0.999 | 0.999 | 0 |
| **ALOX5AP** | 0 | 0.292778075572331 | 0.343 | 0.002 | 0 |
| **HSPH1** | 0 | 0.67350566303169 | 0.994 | 0.979 | 0 |
| **LACC1** | 0 | 0.845237283135606 | 0.882 | 0.301 | 0 |
| **TSC22D1** | 0 | 0.63310880736809 | 0.997 | 0.983 | 0 |
| **TPT1** | 0 | 0.569027038453267 | 1 | 1 | 0 |
| **ZC3H13** | 0 | -0.423554999707876 | 0.972 | 0.99 | 0 |
| **ITM2B** | 0 | 0.715696879251183 | 0.987 | 0.94 | 0 |
| **TDRD3** | 0 | -1.29734185849184 | 0.491 | 0.988 | 0 |
| **LMO7** | 0 | 0.402863644960248 | 0.804 | 0.424 | 0 |
| **EDNRB** | 0 | -0.423508124563998 | 0.158 | 0.71 | 0 |
| **DCT** | 0 | -2.34445063913643 | 0.288 | 0.964 | 0 |

| | | | | | |
|---|---|---|---|---|---|
| **FARP1** | 0 | -0.488241420624386 | 0.689 | 0.933 | 0 |
| **IRS2** | 0 | -0.507555430337527 | 0.626 | 0.906 | 0 |
| **ATP11A** | 0 | -0.256610531647135 | 0.295 | 0.7 | 0 |
| **MCF2L** | 0 | -0.702148342805018 | 0.473 | 0.929 | 0 |
| **TMEM255B** | 0 | 0.824492309619459 | 0.93 | 0.364 | 0 |
| **CDC16** | 0 | -0.438918369495783 | 0.699 | 0.913 | 0 |
| **CHAMP1** | 0 | -0.424994804886704 | 0.678 | 0.905 | 0 |
| **LRP10** | 0 | 0.375083997903229 | 0.915 | 0.674 | 0 |
| **STXBP6** | 0 | -0.375465348067837 | 0.452 | 0.847 | 0 |
| **FRMD6** | 0 | -0.450537059111925 | 0.676 | 0.915 | 0 |
| **SAMD4A** | 0 | -0.283494993597696 | 0.398 | 0.749 | 0 |
| **LGALS3** | 0 | -1.09940561518638 | 0.999 | 1 | 0 |
| **HIF1A** | 0 | 0.815160363150209 | 0.983 | 0.906 | 0 |
| **SNAPC1** | 0 | 0.468589237757107 | 0.909 | 0.682 | 0 |
| **SYNE2** | 0 | -0.325350446692446 | 0.33 | 0.763 | 0 |
| **ZFP36L1** | 0 | 1.09665748322585 | 0.994 | 0.923 | 0 |
| **ACTN1** | 0 | 0.749903688342015 | 0.998 | 0.989 | 0 |
| **NUMB** | 0 | 0.414686048863617 | 0.862 | 0.508 | 0 |
| **IRF2BPL** | 0 | 0.777779476814341 | 0.967 | 0.701 | 0 |
| **CALM1** | 0 | -0.849385378130641 | 0.999 | 0.999 | 0 |
| **RPS6KA5** | 0 | -0.315155393993586 | 0.504 | 0.805 | 0 |
| **C14orf132** | 0 | 0.346677092438931 | 0.764 | 0.24 | 0 |
| **HSP90AA1** | 0 | -1.02861020181483 | 1 | 1 | 0 |
| **MOK** | 0 | -0.325140263240056 | 0.652 | 0.888 | 0 |
| **CRIP2** | 0 | 0.471730842057041 | 0.854 | 0.485 | 0 |
| **LINC02249** | 0 | -0.308428486614117 | 0.077 | 0.615 | 0 |
| **TRPM1** | 0 | -0.571400702645104 | 0.059 | 0.726 | 0 |
| **KLF13** | 0 | 0.318163195938891 | 0.836 | 0.503 | 0 |
| **SCG5** | 0 | 0.547788104366871 | 0.442 | 0.07 | 0 |
| **THBS1** | 0 | 0.417029080553399 | 0.565 | 0.124 | 0 |
| **ZNF106** | 0 | -0.92565306954366 | 0.953 | 0.995 | 0 |
| **SERF2** | 0 | 0.539640422500016 | 1 | 1 | 0 |

| | | | | | |
|---|---|---|---|---|---|
| **EIF3J** | 0 | -0.402549746209566 | 0.861 | 0.959 | 0 |
| **B2M** | 0 | 1.42735553546064 | 1 | 1 | 0 |
| **SORD** | 0 | -0.469501881639402 | 0.719 | 0.935 | 0 |
| **C15orf48** | 0 | 0.350839218786882 | 0.486 | 0.024 | 0 |
| **SLC24A5** | 0 | -0.378055336751525 | 0.295 | 0.78 | 0 |
| **DTWD1** | 0 | 0.479575354887924 | 0.913 | 0.595 | 0 |
| **CCPG1** | 0 | 0.756882431074985 | 0.894 | 0.586 | 0 |
| **MYO1E** | 0 | -0.583627990805556 | 0.725 | 0.931 | 0 |
| **ANXA2** | 0 | 0.6203127918329 | 0.996 | 0.992 | 0 |
| **TPM1** | 0 | 0.688547290409154 | 0.93 | 0.76 | 0 |
| **RPS27L** | 0 | 0.645563972824679 | 0.998 | 0.99 | 0 |
| **ITGA11** | 0 | 0.481262621644079 | 0.684 | 0.017 | 0 |
| **GLCE** | 0 | 0.322365913356566 | 0.814 | 0.521 | 0 |
| **UACA** | 0 | 1.00711978587817 | 0.979 | 0.84 | 0 |
| **PKM** | 0 | 0.939213049254632 | 1 | 1 | 0 |
| **NPTN** | 0 | 0.322393630046906 | 0.874 | 0.697 | 0 |
| **COX5A** | 0 | -0.488579868445854 | 0.998 | 0.999 | 0 |
| **CSPG4** | 0 | -0.403125339313169 | 0.476 | 0.856 | 0 |
| **ABHD17C** | 0 | 0.557948471163777 | 0.817 | 0.293 | 0 |
| **CEMIP** | 0 | 1.0158420959637 | 0.741 | 0.045 | 0 |
| **TLNRD1** | 0 | -0.410618218087261 | 0.536 | 0.857 | 0 |
| **AKAP13** | 0 | 0.330607970864987 | 0.903 | 0.688 | 0 |
| **RLBP1** | 0 | -0.578955546493588 | 0.042 | 0.754 | 0 |
| **MCTP2** | 0 | 0.719776735201474 | 0.871 | 0.291 | 0 |
| **NR2F2** | 0 | 0.81930163583545 | 0.994 | 0.948 | 0 |
| **POLR3K** | 0 | -0.421768153652329 | 0.928 | 0.976 | 0 |
| **RHBDF1** | 0 | 0.343717329833987 | 0.806 | 0.428 | 0 |
| **MPG** | 0 | 0.704067531384663 | 0.994 | 0.987 | 0 |
| **CLCN7** | 0 | -0.311190686683964 | 0.772 | 0.905 | 0 |
| **MRPS34** | 0 | -0.412137333281867 | 0.996 | 0.998 | 0 |
| **RPS2** | 0 | -0.595911941165494 | 1 | 1 | 0 |
| **HCFC1R1** | 0 | 0.944608282578205 | 0.979 | 0.918 | 0 |

| | | | | | |
|---|---|---|---|---|---|
| **TRAP1** | 0 | -0.424804941811801 | 0.86 | 0.959 | 0 |
| **SOCS1** | 0 | 0.332012511563152 | 0.563 | 0.057 | 0 |
| **RSL1D1** | 0 | -0.431461570133315 | 0.986 | 0.997 | 0 |
| **SNX29** | 0 | 0.373304331766918 | 0.798 | 0.293 | 0 |
| **C16orf45** | 0 | 0.250448686723976 | 0.671 | 0.159 | 0 |
| **RPS15A** | 0 | -0.557813133934821 | 1 | 1 | 0 |
| **METTL9** | 0 | 1.00003980172701 | 0.999 | 0.998 | 0 |
| **SPNS1** | 0 | -0.313750672120308 | 0.773 | 0.92 | 0 |
| **LAT** | 0 | 0.274811968430451 | 0.813 | 0.508 | 0 |
| **MAZ** | 0 | -0.35476342914395 | 0.923 | 0.975 | 0 |
| **MVP** | 0 | 0.299283027274984 | 0.675 | 0.112 | 0 |
| **SEZ6L2** | 0 | 0.489361306992274 | 0.934 | 0.699 | 0 |
| **YPEL3** | 0 | 0.625344263561405 | 0.911 | 0.517 | 0 |
| **DCTPP1** | 0 | -0.497532300886328 | 0.881 | 0.974 | 0 |
| **PYCARD** | 0 | -0.785073722658395 | 0.903 | 0.979 | 0 |
| **TGFB1I1** | 0 | 0.36661731651495 | 0.862 | 0.597 | 0 |
| **IRX3** | 0 | 0.358476849352024 | 0.823 | 0.436 | 0 |
| **MMP2** | 0 | 1.34259146881094 | 0.967 | 0.316 | 0 |
| **MT2A** | 0 | 3.12151582899319 | 1 | 0.995 | 0 |
| **MT1F** | 0 | 0.270984371300241 | 0.671 | 0.264 | 0 |
| **MT1X** | 0 | 0.981683493163009 | 0.957 | 0.851 | 0 |
| **NUP93** | 0 | -0.296311157544078 | 0.645 | 0.88 | 0 |
| **ADGRG1** | 0 | -1.42498335137562 | 0.483 | 0.989 | 0 |
| **GOT2** | 0 | -0.398302437928914 | 0.862 | 0.966 | 0 |
| **CKLF** | 0 | 0.444454446808059 | 0.875 | 0.563 | 0 |
| **NQO1** | 0 | 1.21292352226152 | 0.998 | 0.989 | 0 |
| **BCAR1** | 0 | 0.57044252158696 | 0.935 | 0.782 | 0 |
| **CENPN** | 0 | -0.38220615073434 | 0.614 | 0.891 | 0 |
| **GCSH** | 0 | -0.730535131007672 | 0.923 | 0.992 | 0 |
| **CDH13** | 0 | 0.298704240205531 | 0.656 | 0.044 | 0 |
| **TRAPPC2L** | 0 | -0.460274353812484 | 0.946 | 0.988 | 0 |
| **RFLNB** | 0 | -0.644114519107121 | 0.56 | 0.938 | 0 |

| | | | | | |
|---|---|---|---|---|---|
| **TSR1** | 0 | -0.466681445652561 | 0.886 | 0.972 | 0 |
| **SGSM2** | 0 | -0.269161895650451 | 0.584 | 0.827 | 0 |
| **CLUH** | 0 | -0.287697152463729 | 0.638 | 0.866 | 0 |
| **C1QBP** | 0 | -0.831203827486916 | 0.999 | 1 | 0 |
| **DHX33** | 0 | -0.907837738976825 | 0.824 | 0.987 | 0 |
| **NLRP1** | 0 | 0.263201347770746 | 0.601 | 0.089 | 0 |
| **EIF5A** | 0 | -0.611308466891276 | 0.991 | 0.999 | 0 |
| **CD68** | 0 | 0.39969414170179 | 0.928 | 0.78 | 0 |
| **MYH10** | 0 | -1.19411366964894 | 0.905 | 0.996 | 0 |
| **GAS7** | 0 | -0.625971733455655 | 0.788 | 0.972 | 0 |
| **PMP22** | 0 | -0.981177467106642 | 0.966 | 0.996 | 0 |
| **RAB34** | 0 | 0.411401983385729 | 0.988 | 0.965 | 0 |
| **TRAF4** | 0 | -0.288865362376456 | 0.401 | 0.769 | 0 |
| **MYO1D** | 0 | -0.354013154384989 | 0.19 | 0.749 | 0 |
| **TMEM98** | 0 | -0.812289806271534 | 0.694 | 0.975 | 0 |
| **CCL2** | 0 | 1.07811014623327 | 0.641 | 0.028 | 0 |
| **RPL23** | 0 | -0.544154691113269 | 0.999 | 1 | 0 |
| **CNP** | 0 | -0.400343587230603 | 0.742 | 0.925 | 0 |
| **CAVIN1** | 0 | 0.526613370178896 | 0.911 | 0.65 | 0 |
| **RPL27** | 0 | -0.664434155050109 | 0.999 | 1 | 0 |
| **ETV4** | 0 | -0.722691034450783 | 0.873 | 0.98 | 0 |
| **ITGB3** | 0 | 0.300716171757216 | 0.807 | 0.458 | 0 |
| **ATP5MC1** | 0 | -0.694307161254355 | 0.998 | 0.999 | 0 |
| **PHB** | 0 | -0.661566949933577 | 0.999 | 1 | 0 |
| **NGFR** | 0 | 0.35149102994921 | 0.445 | 0.025 | 0 |
| **ITGA3** | 0 | 1.52491283607675 | 0.991 | 0.448 | 0 |
| **COL1A1** | 0 | 1.27991556312884 | 0.811 | 0.051 | 0 |
| **NME1** | 0 | -0.744435604103262 | 0.996 | 0.999 | 0 |
| **SEPT4** | 0 | -0.374242451974176 | 0.289 | 0.725 | 0 |
| **VMP1** | 0 | 0.769650685263634 | 0.996 | 0.969 | 0 |
| **BCAS3** | 0 | -1.46405984632414 | 0.716 | 0.984 | 0 |
| **PECAM1** | 0 | 0.291325458404778 | 0.6 | 0.01 | 0 |

| | | | | | |
|---|---|---|---|---|---|
| **SMURF2** | 0 | 0.543645159546068 | 0.856 | 0.452 | 0 |
| **RGS9** | 0 | 0.262890092047279 | 0.654 | 0.209 | 0 |
| **PRKCA** | 0 | 0.589963902955065 | 0.774 | 0.135 | 0 |
| **ARSG** | 0 | 0.928313539031775 | 0.954 | 0.521 | 0 |
| **SLC16A6** | 0 | 0.996921414839682 | 0.806 | 0.118 | 0 |
| **WIPI1** | 0 | 1.02529789056831 | 0.991 | 0.939 | 0 |
| **LINC00511** | 0 | -0.39106545570053 | 0.597 | 0.861 | 0 |
| **SLC39A11** | 0 | -0.392623042090818 | 0.607 | 0.872 | 0 |
| **TTYH2** | 0 | -0.627699827434388 | 0.241 | 0.897 | 0 |
| **ATP5PD** | 0 | 0.416050816609864 | 0.996 | 0.995 | 0 |
| **MRPS7** | 0 | -0.421835374825299 | 0.988 | 0.994 | 0 |
| **H3F3B** | 0 | 0.461701661187223 | 0.999 | 1 | 0 |
| **UBALD2** | 0 | 0.650486450366069 | 0.96 | 0.859 | 0 |
| **SPHK1** | 0 | 0.450079677105597 | 0.883 | 0.6 | 0 |
| **MXRA7** | 0 | 0.604421781410201 | 0.983 | 0.878 | 0 |
| **SEPT9** | 0 | -0.793599338900243 | 0.963 | 0.998 | 0 |
| **TMC6** | 0 | -0.493675368565278 | 0.476 | 0.902 | 0 |
| **TBC1D16** | 0 | -0.926304220134909 | 0.848 | 0.991 | 0 |
| **NDUFAF8** | 0 | -0.566514505093914 | 0.999 | 1 | 0 |
| **TSPAN10** | 0 | -1.15673551404917 | 0.718 | 0.954 | 0 |
| **MRPL12** | 0 | -0.515796393649474 | 0.986 | 0.997 | 0 |
| **PYCR1** | 0 | -0.321220712569792 | 0.722 | 0.914 | 0 |
| **DUS1L** | 0 | -0.366033591147489 | 0.929 | 0.975 | 0 |
| **FASN** | 0 | -0.445281717067595 | 0.955 | 0.987 | 0 |
| **SLC16A3** | 0 | 1.40453567263153 | 0.995 | 0.957 | 0 |
| **METRNL** | 0 | 0.308279746698089 | 0.808 | 0.476 | 0 |
| **TGIF1** | 0 | 0.345841887788831 | 0.764 | 0.288 | 0 |
| **RAB31** | 0 | 0.483638426717477 | 0.838 | 0.269 | 0 |
| **VAPA** | 0 | 0.351983816614316 | 0.967 | 0.921 | 0 |
| **SNRPD1** | 0 | -0.636650728842296 | 0.989 | 0.997 | 0 |
| **CABLES1** | 0 | -0.536611632934396 | 0.535 | 0.912 | 0 |
| **NPC1** | 0 | 0.548030048266664 | 0.862 | 0.442 | 0 |

| | | | | | |
|---|---|---|---|---|---|
| **ANKRD29** | 0 | 0.329925091506091 | 0.601 | 0.014 | 0 |
| **OSBPL1A** | 0 | -0.468799648442055 | 0.761 | 0.959 | 0 |
| **RNF125** | 0 | -0.351789120038871 | 0.17 | 0.756 | 0 |
| **GALNT1** | 0 | -0.338158241331973 | 0.695 | 0.902 | 0 |
| **SLC39A6** | 0 | -0.36158918177797 | 0.829 | 0.952 | 0 |
| **RAB27B** | 0 | 0.903545418519134 | 0.916 | 0.126 | 0 |
| **NEDD4L** | 0 | -0.654908705491115 | 0.765 | 0.958 | 0 |
| **SEC11C** | 0 | -0.738512328327133 | 0.938 | 0.994 | 0 |
| **SERPINB8** | 0 | 0.28137049331996 | 0.661 | 0.147 | 0 |
| **ZADH2** | 0 | -0.344151303969442 | 0.567 | 0.838 | 0 |
| **MBP** | 0 | 1.36953280979298 | 0.996 | 0.987 | 0 |
| **PARD6G** | 0 | -0.301000016810301 | 0.531 | 0.831 | 0 |
| **FKBP1A** | 0 | 0.760388629802388 | 1 | 1 | 0 |
| **SIRPB1** | 0 | 0.670901339453705 | 0.621 | 0.008 | 0 |
| **SIRPA** | 0 | -1.08804584535651 | 0.493 | 0.971 | 0 |
| **SNRPB** | 0 | -0.59304635043349 | 0.992 | 0.997 | 0 |
| **NOP56** | 0 | -0.730892683602512 | 0.994 | 0.999 | 0 |
| **MRPS26** | 0 | -0.500799021079842 | 0.99 | 0.996 | 0 |
| **RNF24** | 0 | 0.497533690264355 | 0.918 | 0.608 | 0 |
| **PRNP** | 0 | 2.19433886094994 | 0.999 | 0.926 | 0 |
| **PLCB4** | 0 | -0.700066481275346 | 0.325 | 0.916 | 0 |
| **SNRPB2** | 0 | -0.476876546155594 | 0.995 | 0.998 | 0 |
| **DSTN** | 0 | 1.32741260947119 | 1 | 1 | 0 |
| **CST3** | 0 | 1.71749464380466 | 1 | 1 | 0 |
| **CST7** | 0 | 0.670637664670535 | 0.538 | 0.049 | 0 |
| **COMMD7** | 0 | 0.526446250384466 | 0.989 | 0.957 | 0 |
| **AHCY** | 0 | -0.585394514472113 | 0.988 | 0.998 | 0 |
| **PROCR** | 0 | 0.38523010018677 | 0.92 | 0.735 | 0 |
| **MMP24OS** | 0 | 0.713212150103923 | 0.99 | 0.963 | 0 |
| **TGIF2** | 0 | -0.277864622079438 | 0.253 | 0.721 | 0 |
| **TGM2** | 0 | 0.760079183866303 | 0.784 | 0.169 | 0 |
| **MYBL2** | 0 | -0.534836038581045 | 0.376 | 0.791 | 0 |

| | | | | | |
|---|---|---|---|---|---|
| **TOX2** | 0 | 1.00124325938775 | 0.947 | 0.3 | 0 |
| **PKIG** | 0 | 0.568232991931408 | 0.961 | 0.75 | 0 |
| **CTSA** | 0 | 0.717775320356764 | 0.998 | 0.993 | 0 |
| **NCOA3** | 0 | 0.574153996403454 | 0.875 | 0.495 | 0 |
| **B4GALT5** | 0 | -0.374911579111912 | 0.805 | 0.941 | 0 |
| **CEBPB** | 0 | 1.56389075631581 | 0.996 | 0.929 | 0 |
| **NFATC2** | 0 | -0.415238376310443 | 0.112 | 0.68 | 0 |
| **TFAP2C** | 0 | 0.413835487792167 | 0.801 | 0.321 | 0 |
| **AL035541.1** | 0 | -1.1747577957934 | 0.203 | 0.943 | 0 |
| **PMEPA1** | 0 | 0.385569850720154 | 0.622 | 0.127 | 0 |
| **GNAS** | 0 | 0.71641269801575 | 1 | 1 | 0 |
| **CTSZ** | 0 | 0.514679735417877 | 0.965 | 0.884 | 0 |
| **AL162457.2** | 0 | -0.347440483277615 | 0.202 | 0.701 | 0 |
| **PSMA7** | 0 | -0.866098278677613 | 1 | 1 | 0 |
| **SS18L1** | 0 | -0.368948789466826 | 0.481 | 0.817 | 0 |
| **SLCO4A1** | 0 | 0.533005167641933 | 0.855 | 0.634 | 0 |
| **STMN3** | 0 | 0.436323741042078 | 0.944 | 0.794 | 0 |
| **PLEKHJ1** | 0 | -0.339592324567365 | 0.864 | 0.955 | 0 |
| **OAZ1** | 0 | 0.42179538728965 | 1 | 1 | 0 |
| **LSM 7.00** | 0 | -0.740627240235563 | 0.984 | 0.999 | 0 |
| **TIMM13** | 0 | -0.620673737822061 | 0.983 | 0.995 | 0 |
| **GADD45B** | 0 | 0.282138534765921 | 0.717 | 0.27 | 0 |
| **MFSD12** | 0 | -1.07168795276808 | 0.984 | 0.998 | 0 |
| **PLIN3** | 0 | 0.457677632521684 | 0.928 | 0.789 | 0 |
| **TUBB4A** | 0 | -0.388399254167977 | 0.269 | 0.764 | 0 |
| **CD70** | 0 | 0.438840010152069 | 0.701 | 0.187 | 0 |
| **CD320** | 0 | -0.630165228358509 | 0.93 | 0.992 | 0 |
| **MRPL4** | 0 | -0.356288143420105 | 0.958 | 0.987 | 0 |
| **KANK2** | 0 | 0.425729601072813 | 0.943 | 0.795 | 0 |
| **TMEM205** | 0 | 0.623160030520588 | 0.94 | 0.665 | 0 |
| **PLPPR2** | 0 | 0.26264937101315 | 0.773 | 0.445 | 0 |
| **JUNB** | 0 | 1.05274867633208 | 0.952 | 0.692 | 0 |

| | | | | | |
|---|---|---|---|---|---|
| **IER2** | 0 | 0.671333428039607 | 0.948 | 0.724 | 0 |
| **AC020916.1** | 0 | 0.337988052421628 | 0.722 | 0.182 | 0 |
| **GIPC1** | 0 | -0.47245367812508 | 0.919 | 0.981 | 0 |
| **TPM4** | 0 | 1.24724107971814 | 0.998 | 0.976 | 0 |
| **KLF2** | 0 | 0.290728445861108 | 0.575 | 0.135 | 0 |
| **BST2** | 0 | 0.450373513181365 | 0.561 | 0.089 | 0 |
| **JUND** | 0 | 0.840523201232193 | 0.982 | 0.871 | 0 |
| **FKBP8** | 0 | 0.450810823177093 | 0.975 | 0.937 | 0 |
| **FXYD3** | 0 | -1.56122081607774 | 0.183 | 0.99 | 0 |
| **USF2** | 0 | -0.38835876700031 | 0.941 | 0.983 | 0 |
| **LINC01531** | 0 | -0.251188089273011 | 0.031 | 0.503 | 0 |
| **GAPDHS** | 0 | -0.750877707356595 | 0.078 | 0.848 | 0 |
| **TMEM147** | 0 | -0.460214164001933 | 0.996 | 0.999 | 0 |
| **TIMM50** | 0 | -1.22343011840509 | 0.946 | 0.998 | 0 |
| **DLL3** | 0 | -0.409685038307535 | 0.639 | 0.912 | 0 |
| **FBL** | 0 | -0.392357564009892 | 0.945 | 0.989 | 0 |
| **PLD3** | 0 | 0.432832559671757 | 0.933 | 0.683 | 0 |
| **MIA** | 0 | -0.560124116639399 | 0.257 | 0.717 | 0 |
| **AXL** | 0 | 0.448332408334407 | 0.63 | 0.019 | 0 |
| **TGFB1** | 0 | 0.975323639125765 | 0.967 | 0.57 | 0 |
| **RABAC1** | 0 | 0.594666931604872 | 0.994 | 0.956 | 0 |
| **POU2F2** | 0 | 0.328398919233201 | 0.632 | 0.146 | 0 |
| **CEACAM1** | 0 | -0.535413027639718 | 0.207 | 0.791 | 0 |
| **ETHE1** | 0 | 0.74092546418315 | 0.971 | 0.791 | 0 |
| **PLAUR** | 0 | 1.42512560596777 | 0.97 | 0.471 | 0 |
| **KCNN4** | 0 | 0.453001624678785 | 0.81 | 0.154 | 0 |
| **BCL3** | 0 | 0.290841972309609 | 0.682 | 0.16 | 0 |
| **TOMM40** | 0 | -0.483176913754716 | 0.931 | 0.986 | 0 |
| **FOSB** | 0 | 0.295142704497694 | 0.422 | 0.102 | 0 |
| **RTN2** | 0 | 0.261244942714785 | 0.784 | 0.448 | 0 |
| **MEIS3** | 0 | 0.312655527275702 | 0.7 | 0.179 | 0 |
| **EHD2** | 0 | 0.282336899171768 | 0.732 | 0.225 | 0 |

| | | | | | |
|---|---|---|---|---|---|
| **RPL18** | 0 | -0.52343831308582 | 1 | 1 | 0 |
| **PPP1R15A** | 0 | 0.481908068439781 | 0.966 | 0.844 | 0 |
| **NUCB1** | 0 | 0.362401822280829 | 0.943 | 0.816 | 0 |
| **FTL** | 0 | 1.13760523453484 | 1 | 1 | 0 |
| **RRAS** | 0 | 0.858927196820058 | 0.957 | 0.465 | 0 |
| **TBC1D17** | 0 | 0.256553984133541 | 0.785 | 0.447 | 0 |
| **CLEC11A** | 0 | 0.848019358760397 | 0.989 | 0.973 | 0 |
| **TNNT1** | 0 | 0.674791268068427 | 0.988 | 0.95 | 0 |
| **IL11** | 0 | 1.04824121662176 | 0.749 | 0.031 | 0 |
| **BID** | 0 | -0.557783751446036 | 0.746 | 0.961 | 0 |
| **RANBP1** | 0 | -0.76350377520494 | 0.994 | 0.999 | 0 |
| **YDJC** | 0 | -0.35029896141961 | 0.916 | 0.971 | 0 |
| **PRAME** | 0 | -0.501961744292304 | 0.995 | 0.999 | 0 |
| **CHCHD10** | 0 | -0.574482334044295 | 0.988 | 0.997 | 0 |
| **DDT** | 0 | -0.489678324133807 | 0.999 | 1 | 0 |
| **SNRPD3** | 0 | -0.687741970126758 | 0.984 | 0.995 | 0 |
| **HPS4** | 0 | -0.589779276494483 | 0.849 | 0.971 | 0 |
| **TIMP3** | 0 | 2.42221055476831 | 0.993 | 0.991 | 0 |
| **MYH9** | 0 | 0.791808769976586 | 0.992 | 0.953 | 0 |
| **RAC2** | 0 | 0.30328421014253 | 0.751 | 0.414 | 0 |
| **LGALS1** | 0 | 1.75634921126695 | 1 | 1 | 0 |
| **TRIOBP** | 0 | 0.571407249371485 | 0.918 | 0.645 | 0 |
| **H1F0** | 0 | 0.630059524201005 | 0.96 | 0.83 | 0 |
| **GCAT** | 0 | -0.330512418231429 | 0.612 | 0.871 | 0 |
| **POLR2F** | 0 | -0.794518735363186 | 0.992 | 0.998 | 0 |
| **SOX10** | 0 | -1.09690433392172 | 0.172 | 0.982 | 0 |
| **TOMM22** | 0 | -0.445999825044439 | 0.984 | 0.998 | 0 |
| **PRR5** | 0 | -0.389626981375159 | 0.466 | 0.845 | 0 |
| **KIAA0930** | 0 | -0.628741864842798 | 0.835 | 0.974 | 0 |
| **C21orf91** | 0 | -0.438901906520351 | 0.465 | 0.847 | 0 |
| **APP** | 0 | 0.5844440448132 | 0.999 | 0.988 | 0 |
| **MRPS6** | 0 | 1.04735990651489 | 0.997 | 0.985 | 0 |

| | | | | | |
|---|---|---|---|---|---|
| **SLC5A3** | 0 | 1.14640149080178 | 0.974 | 0.896 | 0 |
| **PSMG1** | 0 | -0.3731825876969 | 0.907 | 0.969 | 0 |
| **BACE2** | 0 | -1.14144126319429 | 0.953 | 0.999 | 0 |
| **RRP1B** | 0 | -0.441509417262924 | 0.943 | 0.984 | 0 |
| **RRP1** | 0 | -0.33822479840334 | 0.803 | 0.929 | 0 |
| **PTTG1IP** | 0 | -0.705414749355444 | 0.971 | 0.997 | 0 |
| **COL6A1** | 0 | 2.91244335885161 | 0.997 | 0.637 | 0 |
| **COL6A2** | 0 | 3.04946597739215 | 0.995 | 0.435 | 0 |
| **S100B** | 0 | -1.96006048882432 | 0.807 | 0.998 | 0 |
| **MT-ND2** | 0 | -0.425476691506304 | 0.999 | 1 | 0 |
| **MT-CO1** | 0 | -0.566589264424179 | 1 | 1 | 0 |
| **MT-CO2** | 0 | -0.70815112097724 | 1 | 1 | 0 |
| **MT-ATP6** | 0 | -1.11867410814782 | 1 | 1 | 0 |
| **MT-CO3** | 0 | -0.841542989124886 | 1 | 1 | 0 |
| **MT-ND3** | 0 | -0.590398443950616 | 0.999 | 1 | 0 |
| **MT-ND4L** | 0 | -0.698153279528392 | 0.969 | 0.994 | 0 |
| **MT-ND4** | 0 | -0.867114316204174 | 1 | 1 | 0 |
| **MT-ND5** | 0 | -0.600201768116222 | 1 | 0.999 | 0 |
| **MT-ND6** | 0 | -0.991256061201865 | 0.972 | 0.995 | 0 |
| **MT-CYB** | 0 | -0.994982507933106 | 1 | 1 | 0 |
| **AR** | 6.5658 | 0.284574209300082 | 0.798 | 0.523 | 2.2020682 |
| **ACSL3** | 5.9141 | -0.480891336693077 | 0.978 | 0.993 | 1.9835035 |
| **POLR3D** | 6.9891 | -0.288371581064789 | 0.727 | 0.895 | 2.3440330 |
| **SIGIRR** | 8.0323 | 0.29952515754993 | 0.813 | 0.545 | 2.6938928 |
| **MLF 2.00** | 8.0620 | 0.40091391142227 | 0.998 | 0.997 | 2.7038346 |
| **MYBBP1A** | 1.2813 | -0.285028175615243 | 0.638 | 0.865 | 4.2975415 |
| **TRAPPC1** | 1.6571 | 0.373413647716561 | 0.996 | 0.991 | 5.5575922 |
| **SLC1A3** | 3.1453 | 0.299657604150427 | 0.759 | 0.451 | 1.0548986 |
| **WDFY2** | 2.3094 | 0.272301102578576 | 0.806 | 0.563 | 7.7453850 |
| **ARHGAP18** | 4.8721 | 0.528657613515787 | 0.961 | 0.906 | 1.6340141 |
| **MMD** | 6.2747 | 0.296588292668777 | 0.769 | 0.483 | 2.1044247 |
| **IDS** | 6.2762 | 0.381384960638446 | 0.955 | 0.856 | 2.1049276 |

| | | | | | |
|---|---|---|---|---|---|
| **TAPBP** | 1.7220 | 0.34288990218759 | 0.953 | 0.883 | 5.7752463 |
| **MMP14** | 8.7375 | 0.487073767500305 | 0.928 | 0.763 | 2.9304067 |
| **TXNL4A** | 1.0600 | -0.385886064222206 | 0.987 | 0.994 | 3.5553218 |
| **PFDN2** | 2.1087 | -0.448613262690386 | 0.998 | 0.998 | 7.0724453 |
| **MAP3K11** | 2.8926 | -0.307391699619446 | 0.778 | 0.911 | 9.7013486 |
| **EMP2** | 1.1130 | 0.34714822572621 | 0.926 | 0.768 | 3.7329071 |
| **ANKRD13D** | 1.4567 | 0.280913080281133 | 0.881 | 0.671 | 4.8855329 |
| **HMGN1** | 3.9028 | -0.423262543529878 | 0.998 | 0.999 | 1.3089386 |
| **PFDN5** | 6.1235 | 0.346162441311716 | 1 | 0.999 | 2.0537038 |
| **SOS 1** | 1.0752 | 0.442047521796675 | 0.852 | 0.643 | 3.6062560 |
| **RERE** | 1.0771 | 0.391737548647643 | 0.968 | 0.896 | 3.6123847 |
| **SDF2L1** | 1.8649 | -0.41363230252598 | 0.922 | 0.976 | 6.2548300 |
| **CD55** | 3.2288 | -0.339744923269369 | 0.662 | 0.867 | 1.0828997 |
| **ATP5MF** | 4.2125 | -0.37846469927154 | 0.999 | 1 | 1.4128203 |
| **ZNF689** | 7.8307 | -0.349479571129867 | 0.42 | 0.715 | 2.6262699 |
| **SLC25A33** | 1.2638 | -0.299382631726798 | 0.727 | 0.891 | 4.2387859 |
| **KNOP1** | 1.3117 | -0.290691761083581 | 0.802 | 0.925 | 4.3994391 |
| **TBC1D14** | 1.3307 | -0.275717530137771 | 0.564 | 0.792 | 4.4631524 |
| **CDKN2A** | 3.9871 | 0.653254551516493 | 0.997 | 0.992 | 1.3371993 |
| **PSAP** | 1.7945 | -0.431956299042983 | 1 | 0.999 | 6.0184542 |
| **DYNLRB1** | 1.8667 | 0.328483964552702 | 1 | 0.999 | 6.2605415 |
| **KDELR3** | 3.1752 | 0.307834963190265 | 0.865 | 0.632 | 1.0649123 |
| **TCEAL8** | 3.6704 | 0.302723289756715 | 0.949 | 0.861 | 1.2309864 |
| **TSPO** | 3.8259 | -0.360841554967183 | 0.999 | 1 | 1.2831469 |
| **TWIST1** | 8.7275 | 0.379966178824998 | 0.95 | 0.838 | 2.9270534 |
| **CAPZB** | 1.6614 | 0.400639157624072 | 0.998 | 0.994 | 5.5721640 |
| **SERPINE2** | 1.7787 | 0.761700644593039 | 0.982 | 0.97 | 5.9655805 |
| **JPT1** | 3.0526 | -0.52081470180501 | 0.96 | 0.991 | 1.0237827 |
| **MRTO4** | 4.1170 | -0.385437882281242 | 0.923 | 0.975 | 1.3807870 |
| **PRSS23** | 4.3814 | 0.676185814729801 | 0.899 | 0.802 | 1.4694583 |
| **RAB7A** | 4.6100 | -0.358839962547129 | 0.992 | 0.998 | 1.5461243 |
| **OST4** | 9.6864 | 0.417162190151077 | 0.999 | 0.999 | 3.2486412 |

| | | | | | |
|---|---|---|---|---|---|
| **RRP7A** | 1.1647 | -0.303797522070865 | 0.772 | 0.92 | 3.9062458 |
| **ATP6AP1** | 6.9797 | -0.343519120339007 | 0.931 | 0.978 | 2.3408771 |
| **SEC22B** | 9.9988 | 0.355176991699303 | 0.885 | 0.747 | 3.3533976 |
| **GPATCH4** | 1.3947 | -0.391110152285714 | 0.941 | 0.984 | 4.6777425 |
| **NPM1** | 4.4450 | -0.568343058761471 | 1 | 1 | 1.4907882 |
| **WDR45** | 8.8540 | 0.269899733344504 | 0.87 | 0.656 | 2.9694729 |
| **SEMA3A** | 2.6101 | 0.290248698045313 | 0.601 | 0.287 | 8.7538882 |
| **LY96** | 1.6333 | 0.324252522243944 | 0.918 | 0.755 | 5.4778456 |
| **CHMP5** | 2.2799 | 0.373464902637127 | 0.957 | 0.914 | 7.6464061 |
| **OS9** | 2.6966 | 0.332114593640645 | 0.937 | 0.826 | 9.0439822 |
| **CD164** | 4.4728 | 0.365961476537007 | 0.977 | 0.942 | 1.5001131 |
| **G6PD** | 4.8736 | 0.386664039412908 | 0.962 | 0.924 | 1.6345172 |
| **TXN2** | 4.9654 | -0.328594222764135 | 0.985 | 0.996 | 1.6653097 |
| **NDUFAB1** | 4.0405 | -0.348658945703555 | 0.995 | 0.998 | 1.3551089 |
| **DLC1** | 5.9375 | 0.443187195484872 | 0.841 | 0.672 | 1.9913299 |
| **RPS3** | 1.0429 | -0.464518460593331 | 1 | 1 | 3.4977481 |
| **C8orf33** | 9.5737 | -0.353252024154037 | 0.905 | 0.964 | 3.2108507 |
| **SMARCD2** | 1.7798 | -0.285776703418949 | 0.785 | 0.914 | 5.9693667 |
| **CADPS** | 1.9898 | 0.490134749725201 | 0.832 | 0.615 | 6.6735945 |
| **SLFN5** | 4.4501 | 0.29323584702585 | 0.794 | 0.499 | 1.4924907 |
| **RPS15** | 4.2367 | -0.335121689334187 | 1 | 1 | 1.4209139 |
| **DAD1** | 1.6176 | 0.347037513363004 | 0.998 | 0.993 | 5.4253206 |
| **ITGB8** | 2.2943 | 0.263148096110447 | 0.674 | 0.347 | 7.6946430 |
| **COMMD4** | 3.9552 | -0.355984427105684 | 0.946 | 0.983 | 1.3265222 |
| **YWHAQ** | 4.6404 | 0.389041384356965 | 0.998 | 0.993 | 1.5563162 |
| **TNFRSF12A** | 4.9823 | 0.987711350983441 | 0.863 | 0.764 | 1.6709783 |
| **GRN** | 8.5101 | 0.445559164485491 | 0.991 | 0.984 | 2.8541337 |
| **EXOC3** | 1.5876 | 0.258091244082281 | 0.854 | 0.622 | 5.3246253 |
| **RAB27A** | 4.6563 | -0.408172627594677 | 0.904 | 0.956 | 1.5616608 |
| **GINS2** | 1.4221 | -0.395986650281011 | 0.509 | 0.807 | 4.7696509 |
| **CNPY2** | 3.3286 | -0.444745676265168 | 0.955 | 0.989 | 1.1163762 |
| **TYMS** | 6.0910 | -0.354810603410894 | 0.488 | 0.815 | 2.0428315 |

| | | | | | |
|---|---|---|---|---|---|
| **AFG3L2** | 8.7359 | -0.268046075560584 | 0.68 | 0.861 | 2.9298615 |
| **NDUFAF6** | 9.4132 | -0.264361503684669 | 0.663 | 0.869 | 3.1570042 |
| **SLC7A5** | 1.4952 | -0.312661494220567 | 0.338 | 0.66 | 5.0147099 |
| **SNU13** | 2.5190 | -0.349854355557932 | 0.996 | 0.999 | 8.4484610 |
| **RPL32** | 4.0555 | -0.402300564472852 | 1 | 1 | 1.3601483 |
| **WDR77** | 1.1944 | -0.29769627179884 | 0.766 | 0.915 | 4.0059518 |
| **C20orf27** | 3.3057 | -0.385838027857077 | 0.974 | 0.992 | 1.1086984 |
| **KDELR1** | 4.7578 | 0.354332387487328 | 0.993 | 0.982 | 1.5956833 |
| **SSR4** | 5.7820 | 0.435586218842847 | 1 | 1 | 1.9391988 |
| **RPS18** | 1.3170 | -0.423472895199914 | 1 | 1 | 4.4169749 |
| **CASC4** | 4.5872 | 0.260504564602518 | 0.864 | 0.651 | 1.5384628 |
| **ETS1** | 1.0020 | -0.304574760289893 | 0.529 | 0.796 | 3.3605379 |
| **GLOD4** | 1.1184 | -0.330033524553583 | 0.88 | 0.955 | 3.7511087 |
| **SLC22A18** | 4.1643 | 0.250059211317868 | 0.764 | 0.467 | 1.3966506 |
| **SNRPF** | 6.7827 | -0.399051130464785 | 0.966 | 0.991 | 2.2748014 |
| **MCL1** | 1.0766 | 0.349042375577662 | 0.88 | 0.718 | 3.6108942 |
| **GLA** | 1.6664 | -0.299700081463692 | 0.616 | 0.832 | 5.5889525 |
| **BUD23** | 2.3926 | -0.373547026282048 | 0.976 | 0.99 | 8.0245546 |
| **ARID4B** | 4.2483 | 0.357694341786881 | 0.977 | 0.946 | 1.4247991 |
| **PHGDH** | 1.2822 | -0.443284585950638 | 0.915 | 0.983 | 4.3003614 |
| **REV3L** | 1.4557 | -0.309408439489065 | 0.714 | 0.877 | 4.8824427 |
| **BBX** | 2.0934 | 0.401146624445216 | 0.997 | 0.992 | 7.0210253 |
| **PBDC1** | 3.4028 | -0.316934536569888 | 0.807 | 0.923 | 1.1412527 |
| **LACTB** | 5.7786 | 0.303299170762007 | 0.874 | 0.71 | 1.9380375 |
| **SMAD3** | 6.1201 | 0.25844208345081 | 0.676 | 0.383 | 2.0525780 |
| **KRCC1** | 3.4959 | 0.281349010067652 | 0.9 | 0.756 | 1.1724587 |
| **NDUFS8** | 5.0063 | 0.35082504132984 | 0.992 | 0.98 | 1.6790462 |
| **SNX6** | 7.1816 | 0.30332745245296 | 0.981 | 0.942 | 2.4085868 |
| **VDAC3** | 8.1806 | -0.356586186953297 | 0.94 | 0.977 | 2.7436407 |
| **TMEM183A** | 9.5623 | -0.288590500779882 | 0.858 | 0.949 | 3.2070059 |
| **KRT10** | 1.5454 | 0.375178176244447 | 0.988 | 0.972 | 5.1831646 |
| **COQ10B** | 3.6718 | 0.259520300090292 | 0.862 | 0.682 | 1.2314724 |

| | | | | | |
|---|---|---|---|---|---|
| **P4HB** | 4.2272 | 0.395867542464738 | 1 | 1 | 1.4177343 |
| **SNRNP25** | 7.5632 | -0.366086133976907 | 0.904 | 0.96 | 2.5365673 |
| **SORT1** | 1.3603 | -0.258372474314448 | 0.648 | 0.851 | 4.5625045 |
| **CCT2** | 6.1632 | -0.444424556583604 | 0.984 | 0.994 | 2.0670177 |
| **ATP5PB** | 1.4536 | -0.384979423465336 | 0.987 | 0.997 | 4.8752802 |
| **SERINC3** | 2.7722 | 0.325756956156496 | 0.983 | 0.942 | 9.2975322 |
| **FDFT1** | 8.1375 | -0.362965338551466 | 0.975 | 0.993 | 2.7291593 |
| **EIF1** | 2.0670 | 0.378555167550119 | 0.999 | 0.999 | 6.9324165 |
| **GRSF1** | 2.1058 | -0.321668783183265 | 0.916 | 0.974 | 7.0626493 |
| **MIDN** | 2.3699 | 0.388108276284348 | 0.973 | 0.928 | 7.9482391 |
| **FAM49B** | 6.3261 | -0.310614724423446 | 0.883 | 0.956 | 2.1216547 |
| **CYB5R3** | 1.0533 | 0.336379133452182 | 0.994 | 0.987 | 3.5327357 |
| **BRI3** | 3.0059 | 0.363871273787792 | 1 | 1 | 1.0081401 |
| **ACAA2** | 4.2993 | -0.285632698109825 | 0.865 | 0.945 | 1.4419192 |
| **GXYLT2** | 8.0040 | 0.437898690132812 | 0.954 | 0.884 | 2.6844055 |
| **POP1** | 1.3192 | -0.276802785302078 | 0.636 | 0.839 | 4.4246089 |
| **PLK2** | 3.3990 | -0.271495898432043 | 0.276 | 0.616 | 1.1399857 |
| **TP53TG1** | 3.8625 | 0.296903463350263 | 0.858 | 0.659 | 1.2954102 |
| **DUT** | 1.0200 | -0.474685895178601 | 0.983 | 0.993 | 3.4210034 |
| **OSTM1** | 6.3606 | -0.290563562929991 | 0.783 | 0.923 | 2.1332404 |
| **CD59** | 1.0198 | 0.404787492837636 | 0.989 | 0.977 | 3.4202802 |
| **THEM4** | 2.1193 | -0.301115603039037 | 0.866 | 0.945 | 7.1077623 |
| **ERLEC1** | 2.6657 | 0.307780306342876 | 0.964 | 0.918 | 8.9404481 |
| **CCT6A** | 3.0958 | -0.43213544952199 | 1 | 1 | 1.0383006 |
| **ACADVL** | 3.4988 | 0.361446860341121 | 0.96 | 0.895 | 1.1734528 |
| **HAGHL** | 4.7088 | -0.336364346417936 | 0.858 | 0.942 | 1.5792442 |
| **MAP9** | 5.2556 | 0.267657398175286 | 0.837 | 0.635 | 1.7626247 |
| **RNF213** | 6.0599 | 0.33667741148191 | 0.975 | 0.909 | 2.0323705 |
| **IDH3B** | 9.0911 | -0.324003052223121 | 0.924 | 0.969 | 3.0489898 |
| **NR1H2** | 2.5278 | 0.269882870946615 | 0.915 | 0.789 | 8.4777526 |
| **IQGAP1** | 3.2192 | 0.369654162590119 | 0.985 | 0.95 | 1.0796693 |
| **CBX3** | 6.6365 | -0.395782115805276 | 0.996 | 0.999 | 2.2257562 |

| | | | | | |
|---|---|---|---|---|---|
| **PON2** | 6.7294 | 0.414343743124232 | 0.986 | 0.972 | 2.2569206 |
| **PAICS** | 1.1305 | -0.377308472395988 | 0.944 | 0.986 | 3.7914763 |
| **FAM96B** | 2.5647 | 0.381666705714098 | 0.998 | 0.997 | 8.6015715 |
| **RPL12** | 2.5658 | -0.443679348468925 | 1 | 1 | 8.6052284 |
| **CTDSP2** | 2.7423 | 0.25736919518166 | 0.878 | 0.713 | 9.1971772 |
| **MOSPD3** | 3.7488 | 0.284014981607336 | 0.914 | 0.795 | 1.2572740 |
| **TMEM9** | 7.9060 | 0.261178633633425 | 0.933 | 0.818 | 2.6515255 |
| **METTL23** | 1.1138 | 0.326758637931862 | 0.98 | 0.949 | 3.7355768 |
| **H1FX** | 2.1370 | 0.581565446557612 | 0.984 | 0.964 | 7.1674045 |
| **FAM207A** | 7.5324 | -0.315037488325017 | 0.897 | 0.961 | 2.5262377 |
| **RPS8** | 7.7235 | -0.340249614630304 | 1 | 1 | 2.5903286 |
| **SARAF** | 9.5660 | 0.399443904855451 | 0.993 | 0.982 | 3.2082755 |
| **ZBTB38** | 1.5090 | 0.401920559072103 | 0.956 | 0.889 | 5.0610025 |
| **GALE** | 4.5952 | -0.344742944001414 | 0.894 | 0.956 | 1.5411703 |
| **RASSF3** | 8.8428 | -0.260934953366626 | 0.536 | 0.788 | 2.9657124 |
| **MALT1** | 1.0554 | 0.256242082070664 | 0.801 | 0.568 | 3.5399060 |
| **SAMD9** | 1.5475 | 0.34499647964511 | 0.819 | 0.601 | 5.1903021 |
| **ILK** | 2.1402 | 0.280628622350986 | 0.935 | 0.852 | 7.1778052 |
| **POLR1D** | 2.5873 | -0.321496607122826 | 0.986 | 0.994 | 8.6774294 |
| **UQCR10** | 4.2170 | -0.335037207932664 | 0.999 | 1 | 1.4143235 |
| **NIN** | 5.6244 | -0.291796772912302 | 0.74 | 0.895 | 1.8863404 |
| **RPL38** | 5.9171 | -0.439877664481183 | 0.999 | 1 | 1.9844906 |
| **UBE2R2** | 9.8862 | 0.309818626870414 | 0.962 | 0.918 | 3.3156456 |
| **UCHL5** | 1.0086 | -0.356337007098494 | 0.912 | 0.972 | 3.3826572 |
| **FMN1** | 1.2367 | -0.355994228767944 | 0.827 | 0.931 | 4.1477328 |
| **PHF20L1** | 2.3381 | 0.337421473770644 | 0.978 | 0.959 | 7.8415985 |
| **DMXL2** | 3.8059 | 0.408240697858457 | 0.856 | 0.685 | 1.2764255 |
| **ROCK2** | 4.1437 | 0.358820018411099 | 0.976 | 0.94 | 1.3897268 |
| **CSNK1D** | 6.9111 | 0.304349805556155 | 0.963 | 0.899 | 2.3178742 |
| **SRP14** | 7.9140 | 0.312255853328395 | 1 | 1 | 2.6542154 |
| **PPT1** | 6.9042 | -0.277522023880799 | 0.837 | 0.944 | 2.3155435 |
| **TIMM8B** | 1.9598 | -0.315862080510185 | 0.912 | 0.974 | 6.5728008 |

| | | | | | |
|---|---|---|---|---|---|
| **RPL22** | 3.9001 | -0.2817689221742 | 1 | 1 | 1.3080285 |
| **MZT2A** | 5.8302 | -0.319835499006522 | 0.991 | 0.996 | 1.9553560 |
| **SLC35B2** | 8.3497 | -0.270502408298761 | 0.734 | 0.875 | 2.8003362 |
| **SLIRP** | 7.1544 | -0.39768399078258 | 0.979 | 0.993 | 2.3994436 |
| **MRPS2** | 8.0971 | -0.26416964979274 | 0.801 | 0.917 | 2.7156211 |
| **HYI** | 8.5012 | 0.313615608375044 | 0.902 | 0.766 | 2.8511529 |
| **HOXB2** | 3.6494 | 0.275863639404097 | 0.896 | 0.765 | 1.2239664 |
| **ATP5F1B** | 4.1361 | -0.400307403150892 | 0.996 | 0.999 | 1.3871731 |
| **SHC1** | 4.2979 | 0.290758196756527 | 0.93 | 0.825 | 1.4414391 |
| **PARVB** | 4.4365 | -0.274389786760412 | 0.852 | 0.944 | 1.4879188 |
| **IFRD2** | 6.2698 | -0.285518358871278 | 0.861 | 0.95 | 2.1027711 |
| **EPS8** | 7.2238 | -0.347214421213892 | 0.93 | 0.984 | 2.4227363 |
| **PYGL** | 3.4788 | -0.276068426411418 | 0.626 | 0.842 | 1.1667436 |
| **MTHFD1** | 3.7023 | -0.277573066639308 | 0.707 | 0.887 | 1.2416933 |
| **AC016831.1** | 9.6638 | 0.289942588138934 | 0.744 | 0.468 | 3.2410485 |
| **MRPS9** | 1.7984 | -0.302219401744631 | 0.764 | 0.899 | 6.0317472 |
| **GOLGB1** | 1.9758 | 0.480016804715854 | 0.994 | 0.983 | 6.6264437 |
| **MGST3** | 2.0118 | 0.367975133754618 | 1 | 0.999 | 6.7474970 |
| **UBXN4** | 7.5330 | 0.352687491753073 | 0.991 | 0.969 | 2.5264323 |
| **SDHAF3** | 1.1730 | -0.299694648054691 | 0.864 | 0.949 | 3.9343022 |
| **WWTR1** | 1.3411 | 0.409267245179397 | 0.987 | 0.962 | 4.4979588 |
| **SNRPC** | 2.5031 | -0.308384846916476 | 0.97 | 0.989 | 8.3950115 |
| **NTHL1** | 2.5267 | -0.267180683453781 | 0.801 | 0.915 | 8.4740855 |
| **LEPROT** | 5.6287 | 0.309297920901594 | 0.963 | 0.917 | 1.8877817 |
| **RPA1** | 9.2121 | -0.317992109612297 | 0.868 | 0.954 | 3.0895774 |
| **LDHA** | 6.0984 | 0.63494546006863 | 1 | 0.998 | 2.0453088 |
| **AIMP2** | 8.4784 | -0.258104072536739 | 0.698 | 0.863 | 2.8435007 |
| **CYTL1** | 4.7909 | -0.417286606587288 | 0.875 | 0.961 | 1.6067957 |
| **TMBIM6** | 8.5827 | 0.43106718851791 | 0.999 | 0.998 | 2.8784936 |
| **FAM43A** | 1.5320 | 0.321950127057414 | 0.631 | 0.355 | 5.1381942 |
| **TPD52L1** | 2.6898 | -0.288335391468107 | 0.392 | 0.692 | 9.0213103 |
| **POLR2E** | 2.8529 | -0.295369601607533 | 0.983 | 0.993 | 9.5683755 |

| | | | | | |
|---|---|---|---|---|---|
| **PGRMC2** | 3.3903 | 0.286434922574624 | 0.911 | 0.773 | 1.1370489 |
| **TIMM17A** | 6.4152 | -0.314344842482548 | 0.915 | 0.971 | 2.1515510 |
| **ALDH9A1** | 1.2840 | -0.278049446916342 | 0.78 | 0.914 | 4.3063154 |
| **MDH1** | 2.9318 | -0.423932463517006 | 0.972 | 0.988 | 9.8329814 |
| **MT-ATP8** | 4.2099 | -0.275099909970447 | 0.828 | 0.939 | 1.4119408 |
| **PGAM1** | 7.0499 | -0.370502703763144 | 0.999 | 0.999 | 2.3644136 |
| **NME4** | 1.1991 | -0.373918159677294 | 0.987 | 0.994 | 4.0216581 |
| **SAMM50** | 1.9815 | -0.283373275847642 | 0.829 | 0.928 | 6.6456855 |
| **AAK1** | 1.9060 | 0.296236862344493 | 0.889 | 0.748 | 6.3925828 |
| **MYL12A** | 1.9589 | 0.342455837708044 | 0.976 | 0.954 | 6.5699053 |
| **HDHD5** | 2.5737 | -0.253290211235769 | 0.756 | 0.894 | 8.6319000 |
| **ORC6** | 3.1644 | -0.382043378732448 | 0.593 | 0.834 | 1.0612846 |
| **ATP6AP2** | 9.2867 | 0.367364912984659 | 0.955 | 0.918 | 3.1146006 |
| **GABARAPL2** | 1.6156 | 0.312667332102506 | 0.995 | 0.987 | 5.4186041 |
| **HMGB3** | 9.0323 | -0.267058533962389 | 0.577 | 0.867 | 3.0292669 |
| **STMN1** | 1.9693 | -0.497335847129558 | 0.972 | 0.995 | 6.6049455 |
| **EZH2** | 2.1307 | -0.305955376064931 | 0.583 | 0.828 | 7.1461661 |
| **RPL39** | 1.7311 | -0.402054817126004 | 0.995 | 0.998 | 5.8060342 |
| **M6PR** | 8.7732 | -0.323991089898218 | 0.957 | 0.984 | 2.9423705 |
| **HEBP2** | 1.0930 | 0.316867532937413 | 0.951 | 0.912 | 3.6660157 |
| **BAD** | 2.5401 | 0.270698136103193 | 0.941 | 0.866 | 8.5191718 |
| **WBP11** | 1.3441 | -0.32065074924969 | 0.955 | 0.99 | 4.5080756 |
| **CDIPT** | 2.5252 | 0.263734157071291 | 0.942 | 0.857 | 8.4693264 |
| **ENAH** | 4.4242 | -0.339041032030201 | 0.911 | 0.987 | 1.4838077 |
| **CCNI** | 5.4716 | 0.335651501256338 | 0.999 | 0.998 | 1.8350763 |
| **NDUFC2** | 7.0178 | -0.550000786964412 | 0.998 | 1 | 2.3536490 |
| **EEF1D** | 5.5465 | 0.37455596696607 | 1 | 1 | 1.8602088 |
| **CHML** | 9.0120 | -0.263892676019187 | 0.465 | 0.746 | 3.0224522 |
| **NPC2** | 1.1554 | 0.35902273067671 | 0.997 | 0.989 | 3.8752755 |
| **TRAM1** | 5.8083 | 0.311058943617251 | 0.98 | 0.952 | 1.9480121 |
| **RPL14** | 2.4746 | -0.300673006873915 | 1 | 1 | 8.2995484 |
| **RPS26** | 4.4727 | -0.375580899703002 | 0.999 | 1 | 1.5000766 |

| | | | | | |
|---|---|---|---|---|---|
| **SRPX** | 6.9705 | -0.308260403425363 | 0.842 | 0.943 | 2.3377809 |
| **LARP6** | 4.6451 | 0.29246162226085 | 0.915 | 0.824 | 1.5578750 |
| **PERP** | 2.9122 | 0.347362948600432 | 0.9 | 0.798 | 9.7671438 |
| **MME** | 3.9741 | 0.649392890050341 | 0.906 | 0.822 | 1.3328526 |
| **BSG** | 2.2860 | 0.326856697693743 | 1 | 0.998 | 7.6670061 |
| **CHCHD3** | 2.3236 | -0.312059992773992 | 0.973 | 0.99 | 7.7930157 |
| **PARK7** | 1.4089 | -0.299439987816557 | 0.999 | 1 | 4.7253507 |
| **PPM1G** | 2.5083 | -0.335405952815709 | 0.993 | 0.998 | 8.4123753 |
| **STAT1** | 3.6717 | 0.51209188122732 | 0.948 | 0.921 | 1.2314470 |
| **WDR61** | 6.9669 | -0.252871731462033 | 0.827 | 0.926 | 2.3365678 |
| **IFIT3** | 1.2784 | 0.455252437242684 | 0.669 | 0.377 | 4.2875439 |
| **PPP1R35** | 1.2812 | 0.28989659093364 | 0.954 | 0.904 | 4.2972192 |
| **AMDHD2** | 6.5955 | -0.250880514115959 | 0.753 | 0.879 | 2.2120108 |
| **COX7B** | 8.6376 | -0.320267858562649 | 0.998 | 0.998 | 2.8968814 |
| **PJA2** | 9.4365 | 0.280360291967452 | 0.924 | 0.801 | 3.1648278 |
| **NFIB** | 9.7054 | 0.266079652455786 | 0.792 | 0.567 | 3.2549993 |
| **SPECC1** | 1.2535 | -0.269379236762323 | 0.706 | 0.881 | 4.2040846 |
| **ST8SIA1** | 6.0473 | -0.250439735190812 | 0.68 | 0.888 | 2.0281469 |
| **EXOSC1** | 7.2874 | -0.280535367414772 | 0.892 | 0.956 | 2.4440802 |
| **PDLIM7** | 1.0009 | 0.267101745802823 | 0.813 | 0.632 | 3.3569257 |
| **PTP4A2** | 4.6647 | 0.309247908584361 | 0.997 | 0.995 | 1.5644739 |
| **IFIT1** | 6.9667 | 0.337389497135551 | 0.45 | 0.175 | 2.3365245 |
| **ABRACL** | 8.3528 | -0.265188268567994 | 0.841 | 0.933 | 2.8013727 |
| **NOP16** | 1.2020 | -0.292885229428566 | 0.826 | 0.932 | 4.0314540 |
| **ITGA6** | 5.8495 | -0.264525525673213 | 0.57 | 0.811 | 1.9618053 |
| **GLG1** | 6.4429 | 0.266030556619306 | 0.951 | 0.87 | 2.1608477 |
| **VDAC1** | 5.7477 | -0.361768110051184 | 0.99 | 0.996 | 1.9276732 |
| **ROCK1** | 1.4205 | 0.311624433797742 | 0.982 | 0.96 | 4.7642181 |
| **CBX1** | 1.5986 | -0.318845886081691 | 0.963 | 0.99 | 5.3616170 |
| **NUCKS1** | 2.3535 | -0.41863676736234 | 0.998 | 0.999 | 7.8934601 |
| **NOC2L** | 8.8410 | -0.282931330602238 | 0.9 | 0.962 | 2.9651130 |
| **VMA21** | 1.6815 | -0.285695444253802 | 0.929 | 0.979 | 5.6396934 |

| | | | | | |
|---|---|---|---|---|---|
| **UBE3A** | 4.0975 | -0.313934660296846 | 0.951 | 0.985 | 1.3742242 |
| **RHEB** | 6.0273 | -0.378200262588312 | 0.997 | 0.999 | 2.0214466 |
| **RCAN1** | 6.6784 | -0.262303386824551 | 0.707 | 0.878 | 2.2398057 |
| **ATAD3A** | 7.9951 | -0.262360684052613 | 0.788 | 0.911 | 2.6814158 |
| **SNRPA1** | 1.0518 | -0.345251939791996 | 0.914 | 0.968 | 3.5277790 |
| **BRK1** | 1.0531 | 0.296127098336657 | 0.994 | 0.99 | 3.5319292 |
| **HMG20B** | 1.4204 | -0.38672482911269 | 0.976 | 0.992 | 4.7639073 |
| **NDUFAF4** | 1.4780 | -0.265463378908536 | 0.812 | 0.914 | 4.9571957 |
| **RPL37** | 4.5330 | -0.387596223933272 | 1 | 1 | 1.5202837 |
| **ATP6V1C1** | 5.2860 | -0.301421130584192 | 0.943 | 0.978 | 1.7728254 |
| **TCOF1** | 2.7027 | -0.322404768086735 | 0.855 | 0.947 | 9.0644079 |
| **C4orf3** | 7.0299 | 0.354211786639303 | 0.979 | 0.954 | 2.3576924 |
| **RPS4X** | 9.3490 | -0.350368923874586 | 0.999 | 1 | 3.1354683 |
| **EEF1B2** | 1.9248 | -0.359089254127178 | 1 | 1 | 6.4555569 |
| **TNFRSF14** | 3.1992 | -0.3220623435325 | 0.946 | 0.981 | 1.0729655 |
| **NEAT1** | 8.1439 | 0.984625212769744 | 0.999 | 0.992 | 2.7313186 |
| **TPP1** | 4.8485 | 0.265833270143644 | 0.947 | 0.853 | 1.6261177 |
| **CCDC137** | 1.3583 | -0.305524770495691 | 0.905 | 0.964 | 4.5557365 |
| **PDCD5** | 1.3604 | -0.320639605655348 | 0.995 | 0.998 | 4.5626949 |
| **POMP** | 1.2213 | -0.33176411222713 | 0.999 | 0.999 | 4.0962525 |
| **EIF4A1** | 2.1324 | -0.439885226867507 | 0.996 | 0.999 | 7.1518671 |
| **AP2M1** | 7.7589 | 0.299648749877129 | 0.992 | 0.985 | 2.6021884 |
| **NAV1** | 1.8978 | -0.307981254851074 | 0.792 | 0.902 | 6.3649356 |
| **MCM3** | 2.5971 | -0.374267717741958 | 0.708 | 0.873 | 8.7101545 |
| **TKT** | 3.9360 | -0.346239492595235 | 0.994 | 0.999 | 1.3200754 |
| **ALYREF** | 6.1849 | -0.336234549585887 | 0.88 | 0.958 | 2.0743117 |
| **SEC31A** | 6.4620 | 0.296415313068228 | 0.967 | 0.937 | 2.1672343 |
| **SOAT1** | 3.8253 | -0.332950969393786 | 0.819 | 0.924 | 1.2829530 |
| **SKP1** | 4.0373 | 0.362613705167411 | 0.999 | 0.996 | 1.3540322 |
| **WDR43** | 5.2492 | -0.32259906687035 | 0.861 | 0.948 | 1.7605010 |
| **AGPAT2** | 6.0403 | -0.25490480433811 | 0.865 | 0.934 | 2.0257976 |
| **AURKB** | 9.3079 | -0.435531086187103 | 0.412 | 0.729 | 3.1217140 |

| | | | | | |
|---|---|---|---|---|---|
| **MCFD2** | 1.5066 | 0.291540859779428 | 0.988 | 0.976 | 5.0531256 |
| **COX20** | 2.0069 | 0.337119623785554 | 0.988 | 0.979 | 6.7308859 |
| **BIRC5** | 1.1624 | -0.454820604084203 | 0.599 | 0.882 | 3.8986567 |
| **PLOD3** | 1.5414 | -0.316611342215167 | 0.958 | 0.984 | 5.1695815 |
| **CENPM** | 3.0600 | -0.269230414375573 | 0.613 | 0.889 | 1.0262800 |
| **EIF4G1** | 1.9452 | -0.355652530857288 | 0.969 | 0.989 | 6.5238848 |
| **COMMD6** | 3.6517 | 0.291485751204386 | 0.995 | 0.993 | 1.2247356 |
| **SYNGR1** | 1.4879 | -0.302071078760682 | 0.875 | 0.948 | 4.9903637 |
| **N4BP2L2** | 1.6087 | 0.308234862201806 | 0.978 | 0.936 | 5.3952697 |
| **AIFM1** | 2.2724 | -0.261113240465727 | 0.791 | 0.9 | 7.6214015 |
| **RPL37A** | 1.8951 | -0.363309957576122 | 1 | 1 | 6.3558768 |
| **FBXO32** | 2.0960 | 0.602538180957024 | 0.978 | 0.952 | 7.0298275 |
| **ZNF703** | 4.6547 | -0.353078219540485 | 0.815 | 0.908 | 1.5611088 |
| **TUBGCP2** | 1.4292 | 0.274389448698636 | 0.957 | 0.922 | 4.7935670 |
| **MRPS12** | 3.6754 | -0.285122616406516 | 0.978 | 0.988 | 1.2326703 |
| **CMC2** | 8.5291 | -0.288235522368221 | 0.867 | 0.941 | 2.8605211 |
| **BOLA3** | 1.0081 | -0.337145292071777 | 0.938 | 0.977 | 3.3811112 |
| **NDUFC1** | 1.0359 | -0.334638786814066 | 0.953 | 0.984 | 3.4744150 |
| **LYPLA1** | 1.0722 | -0.292198370985621 | 0.968 | 0.991 | 3.5961308 |
| **CDCA7** | 2.0499 | -0.278259495449583 | 0.496 | 0.768 | 6.8751900 |
| **RABEP1** | 6.6929 | -0.287683644642252 | 0.962 | 0.987 | 2.2446711 |
| **RRAGA** | 9.4420 | 0.277775612238077 | 0.905 | 0.8 | 3.1666602 |
| **WDR46** | 1.0019 | -0.257042636684105 | 0.797 | 0.902 | 3.3604723 |
| **KDM1A** | 1.2235 | -0.285110991570528 | 0.829 | 0.929 | 4.1036226 |
| **HADHA** | 1.4664 | -0.296311835815409 | 0.992 | 0.996 | 4.9182165 |
| **UBL3** | 2.3326 | -0.296044716719892 | 0.802 | 0.912 | 7.8232386 |
| **PRMT1** | 1.3003 | -0.378729373325629 | 0.977 | 0.989 | 4.3612058 |
| **DERA** | 7.3460 | -0.270423394719903 | 0.823 | 0.922 | 2.4637128 |
| **SH3BGRL** | 4.1434 | 0.257583902295669 | 0.949 | 0.894 | 1.3896252 |
| **MRPL27** | 7.6466 | -0.289343530236873 | 0.987 | 0.994 | 2.5645410 |
| **RPL9** | 1.0041 | -0.321236278801538 | 1 | 1 | 3.3676872 |
| **APH1A** | 1.4137 | -0.25761360081563 | 0.939 | 0.975 | 4.7414865 |

| | | | | | |
|---|---|---|---|---|---|
| **APOC1** | 4.176 | -0.286781158259335 | 0.667 | 0.83 | 1.4008228 |
| **PPFIA1** | 4.387 | -0.383033706921836 | 0.998 | 0.998 | 1.4714536 |
| **DESI1** | 5.664 | -0.259989642593926 | 0.9 | 0.96 | 1.8996523 |
| **BLOC1S1** | 6.128 | 0.280492850622655 | 0.975 | 0.958 | 2.0554885 |
| **HIST1H2AC** | 1.204 | 0.419947328799693 | 0.744 | 0.489 | 4.0408591 |
| **IFNGR2** | 1.941 | 0.31873335088178 | 0.977 | 0.959 | 6.5105845 |
| **SKAP2** | 4.265 | 0.25239311555714 | 0.942 | 0.873 | 1.4306059 |
| **NIPSNAP2** | 5.019 | -0.255893927093094 | 0.925 | 0.962 | 1.6833686 |
| **MRPS23** | 1.313 | -0.296608999955542 | 0.98 | 0.988 | 4.4053741 |
| **CYGB** | 3.058 | 0.453125655925051 | 0.991 | 0.989 | 1.0258069 |
| **PRMT2** | 5.799 | -0.26763712064504 | 0.987 | 0.996 | 1.9451440 |
| **RPL36** | 1.292 | -0.262261386683214 | 1 | 1 | 4.3354575 |
| **TNS3** | 1.487 | 0.271315703261524 | 0.923 | 0.826 | 4.9897611 |
| **SYTL2** | 2.065 | -0.25549194439399 | 0.532 | 0.748 | 6.9273834 |
| **MTDH** | 8.923 | -0.297027333032464 | 1 | 0.999 | 2.9927103 |
| **RPL26L1** | 1.796 | -0.279632804345828 | 0.951 | 0.98 | 6.0242808 |
| **UGDH** | 3.575 | -0.293007718124904 | 0.728 | 0.864 | 1.1992846 |
| **DCXR** | 4.702 | -0.289094111214695 | 0.979 | 0.992 | 1.5769606 |
| **SLK** | 4.759 | 0.304295601405552 | 0.951 | 0.888 | 1.5963273 |
| **RPS23** | 6.865 | -0.325907963577097 | 1 | 1 | 2.3026831 |
| **NUDC** | 7.627 | -0.343115608685044 | 0.992 | 0.995 | 2.5581647 |
| **ZC2HC1A** | 8.571 | 0.259380809775787 | 0.904 | 0.784 | 2.8746557 |
| **SMARCA4** | 3.015 | -0.262711622561792 | 0.955 | 0.984 | 1.0113858 |
| **COL9A3** | 4.380 | -0.38396782476206 | 0.656 | 0.86 | 1.4692978 |
| **TMEM165** | 6.370 | 0.300447910998838 | 0.969 | 0.94 | 2.1365321 |
| **PDHA1** | 9.877 | -0.282719334040029 | 0.931 | 0.965 | 3.3128123 |
| **HNRNPC** | 1.320 | -0.331055872364205 | 0.992 | 0.998 | 4.4273119 |
| **TFG** | 1.514 | 0.270058315762888 | 0.994 | 0.982 | 5.0784258 |
| **COX7A2** | 2.282 | -0.276840545030746 | 0.998 | 0.999 | 7.6536594 |
| **RBM8A** | 6.230 | -0.343164241372428 | 0.99 | 0.997 | 2.0895052 |
| **RASSF8-AS1** | 9.352 | 0.277878762532677 | 0.896 | 0.77 | 3.1365302 |
| **HELLS** | 3.272 | -0.329402913062774 | 0.733 | 0.904 | 1.0976801 |

| | | | | | |
|---|---|---|---|---|---|
| **BNIP3** | 2.5802 | 0.642851206696573 | 0.977 | 0.953 | 8.6535555 |
| **METRN** | 2.3581 | -0.346269468987089 | 0.979 | 0.993 | 7.9087799 |
| **PPIH** | 3.6678 | -0.255738847252885 | 0.788 | 0.911 | 1.2301209 |
| **WDR1** | 5.1246 | 0.309481263057614 | 0.975 | 0.962 | 1.7186922 |
| **PPP1CC** | 1.0434 | -0.277374974569728 | 0.966 | 0.991 | 3.4993622 |
| **SRSF3** | 3.5530 | -0.323669565940143 | 0.991 | 0.996 | 1.1916327 |
| **KHSRP** | 3.7549 | -0.342871783828295 | 0.9 | 0.969 | 1.2593390 |
| **GLUL** | 5.5178 | 0.362188845466221 | 0.943 | 0.88 | 1.8505846 |
| **RNF181** | 9.8994 | 0.270874622424875 | 0.982 | 0.961 | 3.3200713 |
| **RECQL** | 1.3936 | 0.335586430264197 | 0.961 | 0.932 | 4.6738681 |
| **AGAP3** | 4.4379 | -0.259108412746847 | 0.918 | 0.956 | 1.4883839 |
| **CALU** | 5.2530 | 0.373550563780695 | 1 | 0.999 | 1.7617676 |
| **MTCH2** | 1.3929 | -0.267338234697554 | 0.938 | 0.982 | 4.6718370 |
| **C3orf14** | 1.5579 | -0.344935242737716 | 0.967 | 0.986 | 5.2249185 |
| **UBN1** | 2.4770 | 0.255725520084149 | 0.945 | 0.882 | 8.3075890 |
| **SPRY1** | 3.3951 | 0.374680241613022 | 0.738 | 0.521 | 1.1386573 |
| **IMP3** | 3.4565 | -0.258810381107539 | 0.929 | 0.964 | 1.1592673 |
| **UTP18** | 5.3991 | -0.282968099373573 | 0.934 | 0.969 | 1.8107643 |
| **PPIF** | 1.7141 | -0.260720651802021 | 0.879 | 0.954 | 5.7490534 |
| **NONO** | 4.2919 | -0.295935066733467 | 0.985 | 0.994 | 1.4394489 |
| **HNRNPF** | 4.9369 | -0.317956422998259 | 0.983 | 0.994 | 1.6557454 |
| **HSPE1** | 6.3063 | -0.391076010253849 | 0.999 | 1 | 2.1150161 |
| **CHAF1A** | 1.2413 | -0.301737023241724 | 0.582 | 0.797 | 4.1633769 |
| **SRFBP1** | 3.8364 | 0.321558061491689 | 0.905 | 0.828 | 1.2866616 |
| **TFDP1** | 7.3027 | -0.287695628593557 | 0.793 | 0.911 | 2.4492069 |
| **UBA2** | 8.3492 | -0.291154837203719 | 0.953 | 0.986 | 2.8001867 |
| **TUFM** | 1.1863 | -0.294059787576714 | 0.992 | 0.996 | 3.9788621 |
| **DEGS1** | 1.6849 | 0.272335854005758 | 0.905 | 0.792 | 5.6509835 |
| **HTRA1** | 2.1156 | 0.581143053171796 | 0.639 | 0.457 | 7.0956177 |
| **HNRNPM** | 1.7124 | -0.356101315064185 | 0.956 | 0.98 | 5.7431238 |
| **CEP250** | 5.4688 | 0.254781302488612 | 0.83 | 0.681 | 1.8341339 |
| **CENPW** | 9.9462 | -0.298387751088344 | 0.663 | 0.884 | 3.3357699 |

| | | | | | |
|---|---|---|---|---|---|
| **PHB2** | 2.1615 | -0.260939448060802 | 0.998 | 0.997 | 7.2492594 |
| **ECM1** | 1.7254 | 0.277237717299537 | 0.846 | 0.664 | 5.7869675 |
| **AURKAIP1** | 2.3613 | -0.269201085020069 | 0.998 | 1 | 7.9194855 |
| **BTG1** | 3.6127 | 0.595069602793852 | 0.994 | 0.987 | 1.2116330 |
| **DHCR24** | 1.0977 | -0.274720997431728 | 0.819 | 0.924 | 3.6815433 |
| **SDHB** | 9.1391 | -0.289606904200873 | 0.955 | 0.978 | 3.0650890 |
| **CSTB** | 1.8584 | -0.324653127683495 | 1 | 1 | 6.2327801 |
| **ZEB2** | 3.9751 | 0.308610443807237 | 0.983 | 0.941 | 1.3331940 |
| **HMGN2** | 6.5346 | -0.349242330201305 | 0.942 | 0.978 | 2.1915935 |
| **CTSH** | 1.1858 | -0.311757500949876 | 0.923 | 0.965 | 3.9770096 |
| **NASP** | 1.3043 | -0.407618778253299 | 0.975 | 0.987 | 4.3745738 |
| **PEPD** | 2.5000 | -0.268067080925261 | 0.898 | 0.947 | 8.3846112 |
| **SLC2A4RG** | 3.2278 | 0.295567390550773 | 0.962 | 0.919 | 1.0825635 |
| **RAI14** | 3.9914 | -0.264879564766627 | 0.924 | 0.984 | 1.3386486 |
| **C6orf48** | 8.5234 | 0.304088243807633 | 0.983 | 0.954 | 2.8586092 |
| **CLIC1** | 1.4900 | 0.304369121100328 | 0.996 | 0.996 | 4.9973030 |
| **ADIPOR1** | 1.6756 | -0.251736772261509 | 0.942 | 0.966 | 5.6197682 |
| **COPB1** | 7.8204 | 0.288801450288582 | 0.918 | 0.834 | 2.6228216 |
| **LBR** | 3.5119 | -0.25707018532835 | 0.648 | 0.814 | 1.1778526 |
| **AK2** | 3.5171 | -0.261779986126797 | 0.964 | 0.982 | 1.1795822 |
| **MCM7** | 2.2019 | -0.325273952483629 | 0.789 | 0.919 | 7.3847478 |
| **PCLAF** | 2.7024 | -0.331397931597238 | 0.682 | 0.906 | 9.0634420 |
| **DDX21** | 3.3573 | -0.281858949412841 | 0.95 | 0.982 | 1.1259895 |
| **CHMP2A** | 6.5830 | 0.261610180086181 | 0.987 | 0.982 | 2.2078159 |
| **LMNA** | 7.5678 | 0.429862549887606 | 0.999 | 0.999 | 2.5381143 |
| **PSME1** | 8.7736 | 0.284525976291402 | 0.972 | 0.947 | 2.9425142 |
| **PDE4B** | 1.2226 | 0.418767150033375 | 0.93 | 0.834 | 4.1003839 |
| **HNRNPD** | 3.6403 | -0.336773949942511 | 0.987 | 0.994 | 1.2209106 |
| **ECHS1** | 4.3404 | -0.269544141912425 | 0.989 | 0.995 | 1.4556951 |
| **MRPS18B** | 6.4391 | -0.250715719170577 | 0.98 | 0.99 | 2.1595475 |
| **HSPA4** | 2.0146 | -0.328441392788912 | 0.952 | 0.979 | 6.7568391 |
| **EIF1AX** | 3.2033 | -0.286469773843552 | 0.98 | 0.993 | 1.0743422 |

| | | | | | |
|---|---|---|---|---|---|
| **SLC2A3** | 1.4615 | 0.273603084178852 | 0.877 | 0.726 | 4.9018001 |
| **TTC3** | 1.7111 | 0.312277293058842 | 0.999 | 0.997 | 5.7389153 |
| **SRSF2** | 1.8368 | -0.293558593960482 | 0.983 | 0.99 | 6.1603995 |
| **EIF2AK2** | 6.0229 | 0.334455016365595 | 0.974 | 0.962 | 2.0199703 |
| **DDX5** | 7.3933 | 0.439579225874628 | 0.997 | 0.996 | 2.4795823 |
| **SCD** | 1.3191 | -0.308019641726856 | 0.988 | 0.997 | 4.4242846 |
| **UBE2N** | 2.1883 | -0.251646915617926 | 0.914 | 0.961 | 7.3392711 |
| **FUCA1** | 1.2157 | 0.293223742559011 | 0.76 | 0.557 | 4.0775248 |
| **FOSL1** | 3.0096 | 0.695036393699145 | 0.831 | 0.718 | 1.0093797 |
| **CFL1** | 3.0547 | 0.279789850536849 | 1 | 1 | 1.0245157 |
| **RAB6A** | 3.4119 | 0.259940843077338 | 0.972 | 0.935 | 1.1442869 |
| **UGCG** | 3.8727 | -0.252685630766314 | 0.572 | 0.754 | 1.2988446 |
| **EIF3B** | 4.5699 | -0.266372311736635 | 0.968 | 0.988 | 1.5326534 |
| **POLR2J3.1** | 3.9517 | 0.291955067221636 | 0.994 | 0.976 | 1.3253351 |
| **TUBA1B** | 8.4275 | -0.584191120282505 | 0.988 | 0.995 | 2.8264187 |
| **HMGN3** | 9.8650 | -0.252957038889752 | 0.972 | 0.985 | 3.3085437 |
| **SLC43A3** | 2.0862 | 0.364840987934262 | 0.985 | 0.969 | 6.9968141 |
| **LSM 5.00** | 4.4234 | -0.30736830363564 | 0.98 | 0.994 | 1.4835246 |
| **LMO4** | 8.0025 | 0.352432266751446 | 0.944 | 0.889 | 2.6838816 |
| **TXNRD1** | 1.3366 | 0.530834904280028 | 0.972 | 0.947 | 4.4827685 |
| **TMEM219** | 2.7156 | 0.26651039974147 | 0.987 | 0.974 | 9.1076621 |
| **HEY1** | 6.0074 | 0.350733842366352 | 0.785 | 0.628 | 2.0147778 |
| **MCM6** | 7.1373 | -0.252684163155373 | 0.533 | 0.761 | 2.3937222 |
| **SQSTM1** | 2.0607 | 0.372170732574402 | 0.999 | 0.999 | 6.9114425 |
| **DNAJC2** | 1.1109 | -0.270189668931321 | 0.965 | 0.985 | 3.7259539 |
| **NUP50** | 3.8343 | -0.260782154902166 | 0.823 | 0.91 | 1.2859542 |
| **RNF7** | 1.2111 | 0.25591355138686 | 0.993 | 0.987 | 4.0619834 |
| **CCT4** | 3.0264 | -0.323290360719077 | 0.989 | 0.995 | 1.0149970 |
| **SSRP1** | 6.3744 | -0.277604092975439 | 0.98 | 0.993 | 2.1378474 |
| **UQCRQ** | 1.7367 | -0.261834666357564 | 0.994 | 0.998 | 5.8247906 |
| **AHCYL1** | 1.8796 | 0.256999718940573 | 0.983 | 0.971 | 6.3040020 |
| **RPS19BP1** | 1.8750 | -0.270679648007534 | 0.991 | 0.996 | 6.2883930 |

| | | | | | |
|---|---|---|---|---|---|
| **DUSP4** | 2.3149 | 0.348476537218587 | 0.997 | 0.992 | 7.7639763 |
| **ATRX** | 3.7150 | 0.28467926116865 | 0.986 | 0.971 | 1.2459611 |
| **LGALS3BP** | 4.1515 | 0.413572981669509 | 0.995 | 0.995 | 1.3923460 |
| **KPNB1** | 4.9597 | -0.293237571242293 | 0.999 | 0.999 | 1.6633871 |
| **NIFK** | 7.9794 | -0.264734489220743 | 0.956 | 0.981 | 2.6761331 |
| **UBR5** | 1.4605 | -0.253684940948478 | 0.918 | 0.963 | 4.8983391 |
| **ATP5F1D** | 3.7762 | -0.296275787441703 | 0.996 | 0.998 | 1.2664776 |
| **SF3A3** | 6.8461 | -0.265418598620137 | 0.918 | 0.96 | 2.2960782 |
| **DNAJC8** | 9.5753 | -0.270594028774934 | 0.995 | 0.998 | 3.2113827 |
| **NME3** | 1.0602 | 0.25024956316936 | 0.967 | 0.925 | 3.5557042 |
| **RPS21** | 1.3038 | -0.288749991729349 | 1 | 1 | 4.3728943 |
| **EEA1** | 2.5683 | -0.2802518513135 | 0.911 | 0.964 | 8.6135772 |
| **PRKDC** | 4.9178 | -0.320137134629786 | 0.996 | 0.999 | 1.6493553 |
| **SSBP1** | 7.4007 | -0.260704350615466 | 0.998 | 0.998 | 2.4820713 |
| **MAD2L1** | 1.0259 | -0.266432325192894 | 0.669 | 0.877 | 3.4409909 |
| **FSTL1** | 1.8582 | -0.258384294412815 | 0.794 | 0.894 | 6.2320644 |
| **LINC00520** | 2.0039 | -0.348975325723368 | 0.259 | 0.508 | 6.7207150 |
| **TPX2** | 4.2704 | -0.395401284661771 | 0.694 | 0.903 | 1.4322160 |
| **RPS19** | 5.9800 | -0.251387072807881 | 1 | 1 | 2.0055763 |
| **MSMO1** | 4.5088 | 0.321058947094339 | 0.839 | 0.693 | 1.5121894 |
| **TPM3** | 1.2965 | -0.298137822307867 | 0.999 | 0.999 | 4.3482593 |
| **EID1** | 2.3549 | 0.291367587690555 | 0.999 | 0.998 | 7.8979672 |
| **CYHR1** | 3.2412 | 0.254885968974055 | 0.977 | 0.951 | 1.0870384 |
| **HNRNPA2B1** | 4.2265 | -0.375783808058896 | 1 | 1 | 1.4174858 |
| **PPDPF** | 2.8951 | 0.293495817522106 | 0.998 | 0.999 | 9.7096581 |
| **HOMER3** | 3.7752 | 0.255231305885041 | 0.974 | 0.965 | 1.2661280 |
| **MKI67** | 1.7950 | -0.409963643080514 | 0.566 | 0.826 | 6.0201607 |
| **PRDX6** | 5.2274 | -0.261021918669101 | 0.994 | 0.998 | 1.7531937 |
| **RUVBL1** | 7.0859 | -0.264191347344109 | 0.949 | 0.98 | 2.3764855 |
| **BRIX1** | 1.4703 | -0.254045963347343 | 0.922 | 0.961 | 4.9314117 |
| **CDC20** | 3.8061 | -0.299836312745444 | 0.483 | 0.73 | 1.2765142 |
| **RPL28** | 5.9789 | 0.309263488615684 | 1 | 1 | 2.0052156 |

| | | | | | |
|---|---|---|---|---|---|
| **DBF4** | 1.6846 | -0.285251055118306 | 0.809 | 0.918 | 5.6498909 |
| **G3BP2** | 1.9752 | -0.277087610527856 | 0.966 | 0.985 | 6.6247495 |
| **LSM 3.00** | 2.6100 | -0.262929369913343 | 0.958 | 0.986 | 8.7536742 |
| **TMEM59** | 7.5016 | 0.340709787355638 | 0.99 | 0.991 | 2.5159180 |
| **RBM28** | 1.7397 | -0.253406883968036 | 0.935 | 0.97 | 5.8347451 |
| **CDKN3** | 3.1374 | -0.255685485005914 | 0.641 | 0.863 | 1.0522491 |
| **ATP6V0B** | 7.4790 | -0.296150967462981 | 0.998 | 0.998 | 2.5083402 |
| **CLSPN** | 9.7728 | -0.319095757397658 | 0.61 | 0.815 | 3.2776142 |
| **AEBP1** | 1.0522 | -0.293238066389472 | 0.949 | 0.985 | 3.5290825 |
| **LRRC59** | 2.4612 | -0.25185804619069 | 0.979 | 0.993 | 8.2545023 |
| **FUS** | 2.4838 | -0.262504978116723 | 0.995 | 0.997 | 8.3302324 |
| **NOP58** | 3.7328 | -0.27838433056766 | 0.898 | 0.957 | 1.2519330 |
| **BZW1** | 1.9723 | 0.313881985455779 | 0.993 | 0.983 | 6.6148841 |
| **NUDT1** | 2.6201 | -0.264403208782872 | 0.974 | 0.99 | 8.7876187 |
| **CCT8** | 1.0457 | -0.29963050299196 | 0.995 | 0.998 | 3.5071167 |
| **TK1** | 4.2854 | -0.265475263271954 | 0.726 | 0.939 | 1.4372447 |
| **SUPT16H** | 4.2001 | -0.26218966520519 | 0.952 | 0.979 | 1.4086306 |
| **MRPL36** | 9.7550 | 0.297601817878079 | 0.994 | 0.992 | 3.2716649 |
| **ATP1B3** | 2.8493 | 0.297817025113877 | 0.993 | 0.988 | 9.5561521 |
| **RRBP1** | 9.5297 | 0.331818775893288 | 0.995 | 0.988 | 3.1960869 |
| **ANP32E** | 1.2229 | -0.265248454210882 | 0.856 | 0.942 | 4.1016444 |
| **TNPO1** | 3.1801 | 0.256008626524916 | 0.961 | 0.945 | 1.0665618 |
| **EEF1A1** | 4.8431 | 0.265097189644526 | 1 | 1 | 1.6243011 |
| **MCM4** | 1.1184 | -0.311094495485434 | 0.817 | 0.907 | 3.7510519 |
| **WLS** | 2.0930 | 0.276896409061691 | 0.893 | 0.817 | 7.0196367 |
| **ARL 2.00** | 2.8051 | 0.258423043343161 | 0.986 | 0.979 | 9.4078286 |
| **RPS6** | 3.4869 | -0.266584563653713 | 1 | 1 | 1.1694455 |
| **SFPQ** | 1.0102 | -0.338119360853688 | 0.983 | 0.994 | 3.3880458 |
| **CYBA** | 1.2410 | 0.32556772645439 | 0.993 | 0.992 | 4.1623182 |
| **SDCBP** | 2.5991 | -0.32880765444646 | 0.997 | 0.999 | 8.7169563 |
| **UBE2S** | 2.9234 | -0.374776268660625 | 0.99 | 0.995 | 9.8045359 |
| **AP002387.2** | 9.1706 | -0.27600611247149 | 0.852 | 0.914 | 3.0756424 |

| | | | | | |
|---|---|---|---|---|---|
| **PTTG1** | 9.6051 | -0.30136245245605 | 0.948 | 0.982 | 3.2213911 |
| **EMP3** | 1.7711 | -0.272681062942033 | 0.999 | 0.998 | 5.9401235 |
| **KRT18** | 4.3444 | 0.35792963080952 | 0.915 | 0.873 | 1.4570344 |
| **CLTA** | 1.2484 | 0.252495987847495 | 0.999 | 0.997 | 4.1871140 |
| **RACK1** | 5.8176 | -0.301911376506754 | 0.999 | 1 | 1.9511304 |
| **PSME2** | 4.1491 | 0.273949290781031 | 0.983 | 0.981 | 1.3915410 |
| **RPL26** | 9.4183 | -0.399043832311702 | 1 | 1 | 3.1587228 |
| **UBE2T** | 7.9582 | -0.259770014591444 | 0.766 | 0.904 | 2.6690359 |
| **GPNMB** | 1.9015 | 0.325878506386381 | 0.999 | 0.999 | 6.3774784 |
| **POLE4** | 2.0912 | 0.253015906134873 | 0.984 | 0.975 | 7.0136072 |
| **FADS1** | 1.7285 | 0.292356328692026 | 0.977 | 0.967 | 5.7973703 |
| **SQLE** | 2.1047 | 0.472383853550016 | 0.978 | 0.965 | 7.0588396 |
| **FXYD5** | 2.2567 | 0.261911089759433 | 0.998 | 0.993 | 7.5685962 |
| **PLIN2** | 3.9700 | 0.449843526836373 | 0.95 | 0.929 | 1.3314641 |
| **SRSF7** | 5.1230 | -0.31283078465796 | 0.981 | 0.991 | 1.7181788 |
| **PDIA3** | 1.2772 | 0.279152878351571 | 0.987 | 0.982 | 4.2835177 |
| **XRCC6** | 1.3391 | -0.254196373705761 | 0.991 | 0.997 | 4.4911999 |
| **SEC61G** | 2.2037 | 0.311886851491364 | 0.999 | 0.998 | 7.3908633 |
| **MARCKS** | 4.8666 | 0.31189907004646 | 0.939 | 0.918 | 1.6321844 |
| **HIST1H1C** | 4.0086 | 0.401751957846823 | 0.952 | 0.894 | 1.3444250 |
| **CKS2** | 5.0647 | -0.320437214468711 | 0.892 | 0.956 | 1.6986190 |
| **KPNA2** | 1.8296 | -0.261602206596474 | 0.893 | 0.949 | 6.1362513 |
| **EIF4A3** | 4.3481 | -0.281145535742665 | 0.917 | 0.94 | 1.4582789 |
| **SOD2** | 5.5149 | 0.287643187623964 | 0.656 | 0.501 | 1.8496118 |
| **TRIB3** | 8.4024 | 0.284755460088586 | 0.814 | 0.702 | 2.8180283 |
| **RPS10** | 1.6078 | -0.525007458073622 | 0.915 | 0.974 | 5.3925571 |
| **RPS17** | 1.9593 | -0.277879505688244 | 0.777 | 0.913 | 6.5713975 |
| **HIPK2** | 7.0249 | -0.26539789588571 | 0.869 | 0.916 | 2.3560173 |
| **CDC25B** | 1.5341 | 0.276370072537852 | 0.937 | 0.894 | 5.1452067 |
| **NAMPT** | 3.2614 | 0.308685416137057 | 0.972 | 0.957 | 1.0938311 |
| **UPP1** | 1.2162 | -0.251277568742028 | 0.949 | 0.973 | 4.0790314 |
| **GAPDH** | 3.8215 | 0.292105345762621 | 1 | 1 | 1.2816811 |

| | | | | | |
|---|---|---|---|---|---|
| **HIST1H1B** | 6.6983 | -0.389529375854405 | 0.546 | 0.695 | 2.2464983 |
| **RPL36A** | 1.4650 | -0.267291817658412 | 0.923 | 0.979 | 4.9134031 |
| **DBI** | 6.9297 | 0.285723438340482 | 1 | 1 | 2.3241086 |
| **HNRNPA3** | 1.8544 | -0.260071760452909 | 0.995 | 0.997 | 6.2194740 |
| **HIST1H4C** | 2.9684 | -0.564400030537216 | 0.968 | 0.969 | 9.9555737 |
| **CAPG** | 6.7852 | -0.48817407911155 | 0.998 | 0.999 | 2.2756354 |
| **TXNIP** | 7.1810 | 0.276406254807731 | 0.778 | 0.643 | 2.4083797 |
| **YWHAZ** | 2.9872 | 0.296674655955707 | 1 | 1 | 1.0018526 |
| **TMX4** | 9.6535 | 0.309463348290544 | 0.94 | 0.91 | 3.2376092 |
| **MAGED2** | 2.8630 | -0.273536511143527 | 0.932 | 0.959 | 9.6022267 |
| **HSP90AB1** | 7.1258 | -0.277822046305995 | 0.999 | 1 | 2.3898824 |
| **GDF15** | 3.0885 | 0.508783660042503 | 0.874 | 0.814 | 1.0358256 |
| **IGFBP5** | 1.4536 | 0.279175781281219 | 0.586 | 0.467 | 4.8751729 |
| **NUPR1** | 3.7012 | 0.35297065589182 | 0.672 | 0.572 | 1.2413326 |
| **IGFBP2** | 6.0372 | 1.19833291634074 | 0.874 | 0.882 | 2.0247705 |
| **ID3** | 2.3542 | -0.266280988282707 | 0.545 | 0.586 | 7.8957362 |
| **FAM89A** | 5.8443 | 0.289395935265607 | 0.989 | 0.988 | 1.9600899 |
| **MTRNR2L8** | 2.8426 | -0.26764352305008 | 0.992 | 0.994 | 9.5336278 |
| **MTRNR2L10** | 4.6241 | -0.324257412126074 | 0.968 | 0.966 | 1.5508367 |
| **ISG15** | 9.0240 | 0.528234733442954 | 0.973 | 0.979 | 3.0264725 |
| **HIST1H1D** | 2.9081 | -0.255226445860764 | 0.879 | 0.873 | 9.7533596 |
