## Supplemental Table 2 for "Disrupting cellular memory to overcome drug resistance"

| | | |
|---|---|---|
| Primers used to amplify lineage barcodes from 10X cDNA<br>sequence in red is the index sequence | GH_i5.1.1 | AATGATACGGCGACCACCGAGATCTACACCTAGCGCTACACTCTTTCCCTACACGACGCTCTTCCGATCT |
| | GH_i5.2.1 | AATGATACGGCGACCACCGAGATCTACACTCGATATCACACTCTTTCCCTACACGACGCTCTTCCGATCT |
| | GH_i7.3.1 | CAAGCAGAAGACGGCATACGAGATGTCTCTACGTGACTGGAGTTCAGACGTGTGCTCTTCCGATCTAGGACGAGCTGTACAAGTAGG |
| | GH_i7.3.2 | CAAGCAGAAGACGGCATACGAGATCAGATAGTGTGACTGGAGTTCAGACGTGTGCTCTTCCGATCTCTGGACGAGCTGTACAAGTAGG |
| | GH_i7.3.3 | CAAGCAGAAGACGGCATACGAGATTCTACGCAGTGACTGGAGTTCAGACGTGTGCTCTTCCGATCTTGCGGACGAGCTGTACAAGTAGG |
| | GH_i7.3.4 | CAAGCAGAAGACGGCATACGAGATCGACTCTGGTGACTGGAGTTCAGACGTGTGCTCTTCCGATCTGCTCGGACGAGCTGTACAAGTAGG |
| Primers used to amplify lineage barcodes from gDNA<br>sequence in red is the index sequence<br>blue sequence is a "Stagger sequece" so that | GHi5.2.1 | AATGATACGGCGACCACCGAGATCTACACTAGATCGCACACTCTTTCCCTACACGACGCTCTTCCGATCTNHNNNNTCGACTAAACGCGCTACTTG |
| | GHi5.2.2 | AATGATACGGCGACCACCGAGATCTACACCTCTCTATACACTCTTTCCCTACACGACGCTCTTCCGATCTNHNNNNCGTCGACTAAACGCGCTACTTG |
| | GHi5.2.3 | AATGATACGGCGACCACCGAGATCTACACTATCCTCTACACTCTTTCCCTACACGACGCTCTTCCGATCTNHNNNNGACTCGACTAAACGCGCTACTTG |
| | GHi5.2.4 | AATGATACGGCGACCACCGAGATCTACACAGAGTAGAACACTCTTTCCCTACACGACGCTCTTCCGATCTNHNNNNATAGTCTCGACTAAACGCGCTACTTG |
| | GHi5.2.5 | AATGATACGGCGACCACCGAGATCTACACGTAAGCAGACACTCTTTCCCTACACGACGCTCTTCCGATCTNHNNNNTACGTCGACTAAACGCGCTACTTG |
| | GHi5.2.6 | AATGATACGGCGACCACCGAGATCTACACACTGCGTAACACTCTTTCCCTACACGACGCTCTTCCGATCTNHNNNNCCTAATCGACTAAACGCGCTACTTG |
| | GHi5.2.7 | AATGATACGGCGACCACCGAGATCTACACAAGGAGTAACACTCTTTCCCTACACGACGCTCTTCCGATCTNHNNNNTCGACTAAACGCGCTACTTG |
| | GHi5.2.8 | AATGATACGGCGACCACCGAGATCTACACCTAAGCCTACACTCTTTCCCTACACGACGCTCTTCCGATCTNHNNNNGCTCGACTAAACGCGCTACTTG |
| | GHi5.2.9 | AATGATACGGCGACCACCGAGATCTACACCGTCTAATACACTCTTTCCCTACACGACGCTCTTCCGATCTNHNNNNCTGTCGACTAAACGCGCTACTTG |
| | GHi5.2.10 | AATGATACGGCGACCACCGAGATCTACACTCTCTCCGACACTCTTTCCCTACACGACGCTCTTCCGATCTNHNNNNTATCAGTCGACTAAACGCGCTACTTG |
| | GHi5.2.11 | AATGATACGGCGACCACCGAGATCTACACTCGACTAGACACTCTTTCCCTACACGACGCTCTTCCGATCTNHNNNNATGCTCGACTAAACGCGCTACTTG |
| | GHi5.2.12 | AATGATACGGCGACCACCGAGATCTACACTTCTAGCTACACTCTTTCCCTACACGACGCTCTTCCGATCTNHNNNNGGATATCGACTAAACGCGCTACTTG |
| | GHi7.2.1 | CAAGCAGAAGACGGCATACGAGATTAAGGCGAGTGACTGGAGTTCAGACGTGTGCTCTTCCGATCTGTCCTGCTGGAGTTCGTGAC |
| | GHi7.2.2 | CAAGCAGAAGACGGCATACGAGATCGTACTAGGTGACTGGAGTTCAGACGTGTGCTCTTCCGATCTAGGTCCTGCTGGAGTTCGTGAC |
| | GHi7.2.3 | CAAGCAGAAGACGGCATACGAGATAGGCAGAAGTGACTGGAGTTCAGACGTGTGCTCTTCCGATCTTCAAGTCCTGCTGGAGTTCGTGAC |
| | GHi7.2.4 | CAAGCAGAAGACGGCATACGAGATTCCTGAGCGTGACTGGAGTTCAGACGTGTGCTCTTCCGATCTCACTTAAGTCCTGCTGGAGTTCGTGAC |
| | GHi7.2.5 | CAAGCAGAAGACGGCATACGAGATCGACTCCTGTGACTGGAGTTCAGACGTGTGCTCTTCCGATCTTGTCCTGCTGGAGTTCGTGAC |
| | GHi7.2.6 | CAAGCAGAAGACGGCATACGAGATTAGGCATGGTGACTGGAGTTCAGACGTGTGCTCTTCCGATCTCTTGAGTCCTGCTGGAGTTCGTGAC |
| | GHi7.2.7 | CAAGCAGAAGACGGCATACGAGATATCGCTACGTGACTGGAGTTCAGACGTGTGCTCTTCCGATCTGTAGTCCTGCTGGAGTTCGTGAC |
| | GHi7.2.8 | CAAGCAGAAGACGGCATACGAGATCAGAGAGTGTGACTGGAGTTCAGACGTGTGCTCTTCCGATCTGATACTGTCCTGCTGGAGTTCGTGAC |
